# Can We Ever Develop an Ideal RNA Force Field? Lessons Learned from Simulations of UUCG RNA Tetraloop and Other Systems

**DOI:** 10.1101/2024.10.08.617195

**Authors:** Vojtěch Mlýnský, Petra Kührová, Martin Pykal, Miroslav Krepl, Petr Stadlbauer, Michal Otyepka, Pavel Banáš, Jiří Šponer

**Affiliations:** Institute of Biophysics of the Czech Academy of Sciences, Královopolská 135, 612 00 Brno, Czech Republic; Regional Center of Advanced Technologies and Materials, The Czech Advanced Technology and Research Institute (CATRIN), Palacký University Olomouc, Šlechtitelů 27, 779 00 Olomouc, Czech Republic; IT4Innovations, VSB – Technical University of Ostrava, 17. listopadu 2172/15, 708 00 Ostrava-Poruba, Czech Republic

## Abstract

Molecular dynamics (MD) simulations are an important and well-established tool for investigating RNA structural dynamics, but their accuracy relies heavily on the quality of the employed force field (*ff*). In this work, we present a comprehensive evaluation of widely used pair-additive and polarizable RNA *ff*s using the challenging UUCG tetraloop (TL) benchmark system. Extensive standard MD simulations, initiated from the NMR structure of the 14-mer UUCG TL, revealed that most *ff*s did not maintain the native state, instead favoring alternative loop conformations. Notably, three very recent variants of pair-additive *ff*s, OL3_CP_–gHBfix21, DESAMBER, and OL3_R2.7_, successfully preserved the native structure over a 10 × 20 µs timescale. To further assess these *ff*s, we performed enhanced sampling folding simulations of the shorter 8-mer UUCG TL, starting from the single-stranded conformation. Estimated folding free energies (ΔG°_fold_) varied significantly among these three *ff*s, with values of 0.0 ± 0.6 kcal/mol, 2.4 ± 0.8 kcal/mol, and 7.4 ± 0.2 kcal/mol for OL3_CP_–gHBfix21, DESAMBER, and OL3_R2.7_, respectively. The ΔG°_fold_ value from OL3_CP_–gHBfix21 was closest to experimental data, while the higher ΔG°_fold_ values from DESAMBER and OL3_R2.7_ were unexpected, suggesting an over- or underestimation of key interactions within the folded and mis(un)folded ensembles. These discrepancies led us to further test DESAMBER and OL3_R2.7_ *ff*s on additional RNA and DNA systems, where further performance issues were observed. Our results emphasize the complexity of accurately modeling RNA dynamics and suggest that creating an RNA *ff* capable of reliably performing across a wide range of RNA motifs remains extremely challenging. In conclusion, our study provides valuable insights into the capabilities of current RNA *ff*s and highlights key areas for future *ff* development.

## INTRODUCTION

Nucleic acid (DNA and RNA) molecules are involved in countless cellular processes, physiological as well as pathological ones. While DNA forms primarily canonical Watson-Crick double helices, RNA molecules adopt an astonishingly variable spectrum of three-dimensional folds. Many RNA molecules are dynamical, some even disordered, and most RNA molecules interact with diverse proteins in a cellular environment.^1–5^

Atomistic explicit solvent molecular dynamics (MD) simulations represent a genuine complement of experimental methods in studies of RNA structural dynamics offering very fine spatio-temporal resolution.^6–12^ However, MD simulations have also considerable limitations. First, the accessible timescale of MD simulations is much shorter than timescale of RNA structural dynamics in most physiological processes. This limitation can, in principle, be overcome by using diverse enhanced sampling simulation techniques,^13^ albeit these techniques bring additional and often significant approximations compared to standard unbiased MD simulations.^9^ Second, and perhaps more important, outcome of RNA MD simulations critically depends on capability of an empirical potential, i.e., force field (*ff*), to approximate the potential energy surface of real RNA molecules and their interactions with other molecules.^9, 14–22^ Thus, many biochemical processes involving RNA molecules are challenging for MD simulations. Neglecting limits of MD methodology may lead to overinterpretation of the simulation data, especially when only global metrices are reported and true atomistic details of the trajectories are not carefully monitored.^23^ Contemporary MD simulations cannot also reliably predict RNA structures.^9^ At present, MD techniques are most powerful when used to complement experimental studies that provide high-quality, atomic-resolution structural data as starting points for simulations.^9^

Assessment of *ff* performance, identification of problems and attempts to improve the *ff*s is an ongoing process. A genuine approach for (RNA) *ff* improvement is to compare the simulation data with atomistic experimental structures. The conventional method is to monitor structural stability of native interactions seen in the respective experimental structures available in Protein Data Bank (PDB, https://www.rcsb.org/).^24^ However, such analyses may be affected by the static (averaged) nature of the experimental structures and limited convergence of the simulation trajectories. It is sometimes difficult to separate structural instabilities caused by *ff* inaccuracies from real dynamics, which is masked in the experimental structures due to ensemble averaging. It can be illustrated by X-ray structure capturing binding of AU-rich single stranded RNA (ssRNA) element to the HuR C-terminal RNA recognition motif domain.^25^ Despite the relatively high nominal resolution of 1.9 Å, the most interesting part of the structure, the RNA binding, was poorly visible in electron densities. MD simulations helped to determine that the X-ray data arise from a mixture of different RNA binding patterns and registers.^25^

One of the main issues when parametrizing *ff*s is to ensure they work satisfactorily over a broad range of RNA systems. Taking into account structural diversity of RNA, it is an open question whether we can have a single universal *ff* capable to deal with all kinds of RNA structural dynamics. For example, we have recently carried out a first simulation study of a spontaneous binding of ssRNAs to RNA binding protein domains.^26^ To simulate the binding process, we had to radically scale down van der Waals interactions (not only base stacking, but also sugar – base and sugar – sugar interactions) of the common AMBER RNA OL3^27–30^ *ff*. The *ff* modification called stafix prevented spurious ssRNA self-interactions (essentially a structural collapse) but stafix obviously cannot be used for simulations of structured RNAs.^26^

In the last decade, a set of short RNA molecules has become a prominent benchmark to analyze the *ff* performance and to refine the *ff*s with the help of primary experimental data (NMR data or folding free energies).^15, 16, 18, 19, 31–54^ One of the key systems is the UUCG tetraloop (TL) which possesses well-defined native structure with canonical base pairing in the stems and characteristic noncanonical interactions involving sugar-base and base-phosphate (BPh) hydrogen bonds (H-bonds) in the loop.^15, 18, 34, 39, 42, 45, 48, 54–57^ For the UUCG TL both unambiguous experimental structure and measured folding free energies are available.^31–34, 55–57^

Common *ff* testing involves validating the overall structural stability of the native interactions during standard MD simulations.^22, 41, 58, 59^ More complicated benchmark setups involve calculation of folding free energies (and subsequent comparison with experiments) by enhanced sampling simulations. They enable to assess not only the structural stability of native conformations, but also the balance between the native conformation and the misfolded/unfolded ensembles.^15, 16, 18, 39, 42, 48, 60, 61^ The UUCG TL appears at first sight to be an easy target. It does not contain any RNA tertiary interactions (such as A-minor interactions^62^ and other ribose zippers and backbone interactions^63, 64^) which are fundamentally important in larger folded RNAs. Still, RNA *ff*s have been notoriously struggling to match the experimental data for the UUCG tetraloop.^15, 18, 39, 41, 45, 61^

In this paper, we provide a thorough assessment of the capability of RNA *ff*s to simulate the UUCG TL. Although the UUCG TL is usually included within benchmark set of testing systems by studies introducing new *ff* variants, it is not uncommon that the testing is done on a limited timescale and/or there is rather superficial structural analysis without convincing evidence that the native state was dominantly sampled. Here, we collected an extended set of standard MD simulations starting from the benchmark 2KOC NMR structure of the 14-mer UUCG TL with five base pair stem.^34^ Altogether, we investigated a set of fifteen *ff*s in a series of multiple (typically 10 × 20 µs) MD simulations. Our goal was to compare behavior of common RNA *ff*s (including polarizable ones) in structural description of native interactions which are defining the UUCG native structure. Our study confirms that the UUCG TL is a challenging system for MD simulations. Majority of the tested *ff*s irreversibly disrupted the native state and preferentially sampled alternative loop conformations. Nevertheless, we identified three very recent RNA *ff*s that provided stable structural description of the UUCG 2KOC 14-mer structure on our timescale. For these *ff* variants we performed a set of enhanced sampling folding simulations of the shorter 8-mer UUCG TL with two base-pair stem. The folding simulations revealed less satisfactory results and striking differences of predicted folding free energies among those *ff*s that performed well in standard simulations of the 14-mer UUCG TL. We then performed additional standard simulations on other nucleic acids systems for selected *ff*s to better understand their behavior. These simulations also revealed striking differences among the tested *ff*s. Altogether, we report results that are based on ∼2.3 ms of standard simulations and ∼1.1 ms of enhanced sampling simulations. In summary, our study provides unique insights into the performance of contemporary RNA *ff*s for the structural description of the common UUCG TL benchmark system and unveils important paths for the ongoing *ff* development. The results also underline the enormous intricacy of RNA *ff* parameterizations.

## METHODS

### System preparation and simulation protocols for pair-additive *ff*s

For standard simulations, we used the UUCG NMR structure (PDB ID 2KOC; selecting structure #13 which is characterized by the shortest distances of native H-bonds)^34^ for preparation of starting topology and coordinates of the r(ggcacUUCGgugcc) 14-mer, i.e., the UUCG TL with five base-pair stem. We used smaller r(gcUUCGgc) 8-mer motif for folding simulations (see the paragraph about enhanced sampling simulations below) which were initiated from one strand of A-RNA duplex.

For standard simulations of the 2KOC structure, we tested 13 different setups with recent pair-additive RNA *ff*s (Table 1). Several variants were prepared based on the common^65^ and widely used *ff*99bsc0χ_OL3_ (i.e., OL3) RNA *ff*.^27–30^ The OL3 *ff* was in some setups further adjusted by the van der Waals (vdW) modification of phosphate oxygens developed by Steinbrecher et al.^66^ in combination with refit of the affected α, γ, δ and ζ torsions parameterized by us.^16, 67^ This RNA *ff* version is abbreviated as OL3_CP_ henceforth and AMBER library file can be found in Supporting Information of Ref. ^16^. We used three variants of general H-bond fix (gHBfix) potentials,^18^ i.e., (i) gHBfix19 potential, where all –NH…N– base – base interactions are strengthened by 1.0 kcal/mol and all –OH…bO– and –OH…nbO– sugar – phosphate interactions are weakened by 0.5 kcal/mol,^18^ (ii) gHBfix_UNCG19_ version, where –NH…N– and –NH…O– base – base H-bonds are strengthened by 0.5 kcal/mol, sugar – phosphate interactions are weakened by 0.5 kcal/mol, sugar donor – base acceptor H-bonds are strengthened by 0.5 kcal/mol, and base donor – sugar acceptor and sugar – sugar H-bonds are weakened by 0.5 kcal/mol,^45^ and (iii) the latest optimized gHBfix version from 2021 (either gHBfix_opt_ or gHBfix21 subvariant).^54^ In the gHBfix_opt_ version, all RNA H-bond donor…H-bond acceptor interactions are modified, i.e., base donor – base acceptor, base donor – sugar acceptor, base donor – phosphate acceptor, sugar donor – base acceptor, sugar donor – sugar acceptor, and sugar donor – phosphate acceptor are adjusted specifically (see Ref. ^54^ for full description). The gHBfix_opt_ version was obtained by machine learning, but then the modification of sugar donor – sugar acceptor interactions was excluded because it caused spurious destabilization of A-minor type I interaction in simulations of RNA kink-turn motif. This led to the final gHBfix21 version.^54^ Both gHBfix_opt_ and gHBfix21 versions are very similar (see Table S1 in Supporting Information for complete list of interactions modified by gHBfix_opt_ and gHBfix21potentials) and are expected to provide essentially identical performance for the folded UUCG TL which does not contain any sugar donor – sugar acceptor interactions.^54^ Note that in this work some analyzed simulations were done with gHBfix_opt_ and some with gHBfix21 potentials but, for simplicity, only the gHBfix21 labelling is used henceforth (Table 1).

**Table 1:**
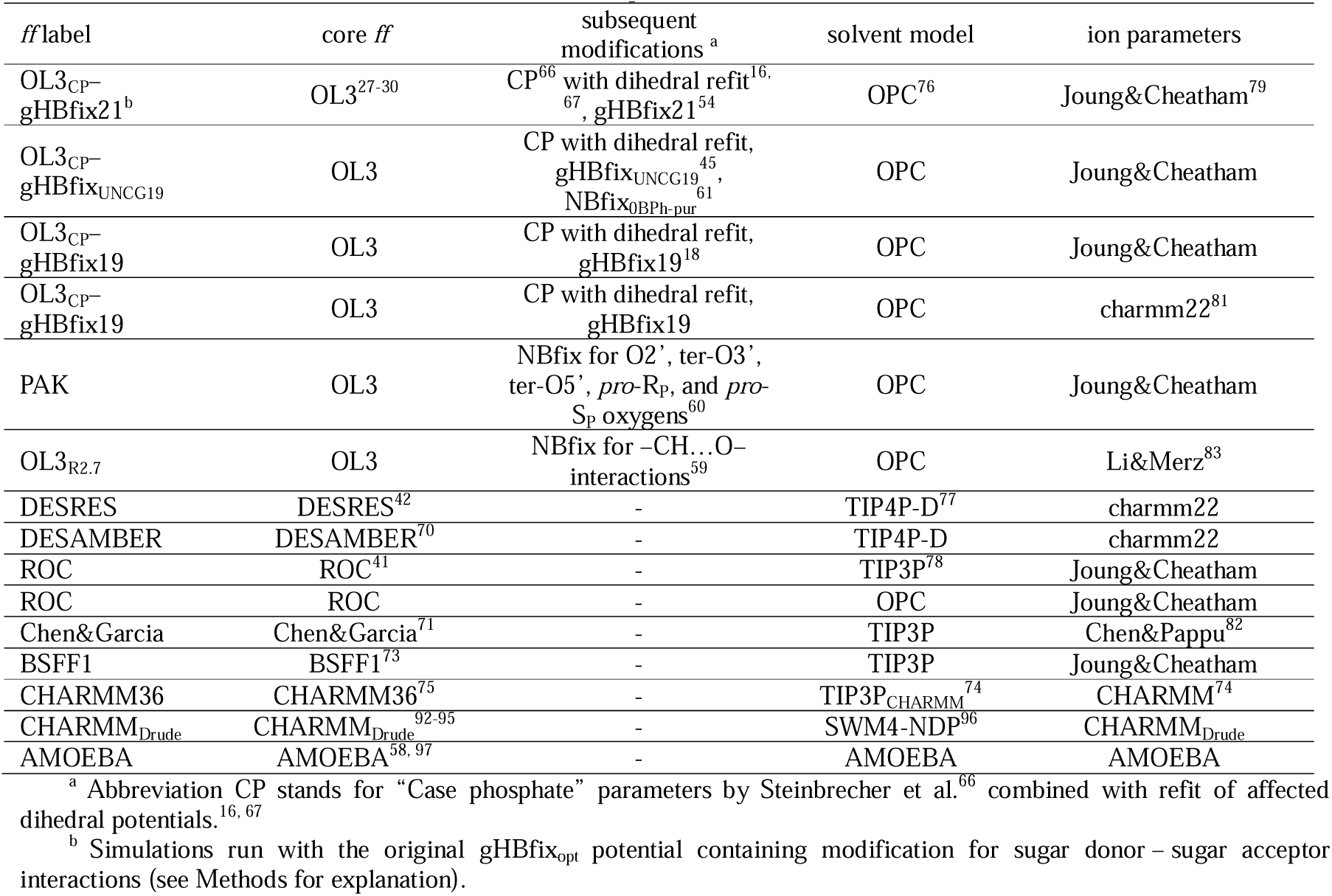
List of RNA *ff*s, water models and ion parameters used in MD simulations.

For the sake of completeness we note that simulations with the OL3_CP_–gHBfix_UNCG19_ *ff* also contained modified pairwise vdW parameters via breakage of combination (mixing) rules, i.e., the so-called nonbonded fix (NBfix) approach,^68^ for atoms involved in BPh interaction type 0 (0BPh).^69^ More specifically, we reduced the minimum-energy distance of Lennard-Jones potential (i.e., the *R_i,j_*parameter) for the –H8…O5’– pair, i.e., between H5 – OR atom types for purines (NBfix_0BPh-pur_; Table 1).^61^

The OL3 *ff* (the standard version without CP modification) was also tested with (i) adjusted vdW parameters of O2’, terminal O3’ and O5’, and non-bridging phosphate (*pro*-R_P_, *pro*-S_P_) oxygens by Yang et al. (PAK *ff*)^60^ and with (ii) very recently suggested general adjustment of pairwise vdW parameters between oxygens and nonpolar hydrogens, i.e., the NBfix applied for most –CH…O– interactions involving nonpolar hydrogens (Hydrogen Repulsion Modification; HRM), using the *R_i,j_* value of 2.7 Å (OL3_R2.7_ *ff*;^59^ Table 1).

Other tested pair-additive *ff*s include (i) reparameterized charges, nonbonded and dihedral parameters by Tan and co-workers (DESRES *ff*),^42^ (ii) subsequent further refinement of nonbonded parameters, charges and dihedrals by the same group (DESAMBER *ff*),^70^ (iii) comprehensive reparameterization of all AMBER dihedral parameters by Aytenfisu et al. (ROC *ff*),^41^ (iv) revised nonbonded and dihedral parameters by Chen and Garcia (Chen&Garcia *ff*),^71^ (v) optimized nonbonded parameters with an inclusion of grid-based energy correction map (CMAP) term^72^ by Li et al. (BSFF1 *ff*),^73^ and the latest CHARMM^74^ *ff* for RNA simulations (CHARMM36;^75^ Table 1).

Both 14-mer and 8-mer UUCG TLs were placed in a cubic box of either OPC^76^ (all OL3_CP_-based variants, OL3_R2.7_, ROC and PAK *ff*s), TIP4P-D^77^ (DESRES *ff*), or TIP3P^78^ (ROC and Chen&Garcia *ff*s) water molecules with minimal distance of 12 Å between the solute and the box border. KCl ions parametrized by Joung&Cheatham^79^ for the TIP4P-EW (OPC) or TIP3P water models were added to neutralize the system and to establish excess-salt ion concentration of ∼0.15 M. Simulations with DESRES *ff* were run with recommended^42, 80^ charmm22 ions^81^ and we further tested charmm22 ions in simulations with the OL3_CP_–gHBfix19 *ff*. Simulations with Chen&Garcia and OL3_R2.7_ *ff*s were run with recommended Chen&Pappu^82^ and Li&Merz^83^ ions, respectively. Note that both OPC and TIP3P water models were tested with the ROC *ff*. Whereas simulations with TIP3P water model were performed under excess-salt concentration of 0.1 M using NaCl ions (Joung&Cheatham parameters; as used in the original ROC paper^41^), we also tested performance of ROC *ff* with OPC water under excess-salt concentration of 0.15 M using KCl ions.

The tLeap program was used to generate initial files (with exception of DESAMBER, BSFF1 and CHARMM36 *ff*s; see below) and simulations were subsequently performed in AMBER18^84^ using the pmemd.MPI and pmemd.cuda^85^ programs for equilibration and production simulations, respectively. All MD simulations were run at T = 298 K with the hydrogen mass repartitioning^86^ allowing 4 fs integration time step. Long-range electrostatics were treated with particle mesh Ewald^87^ and the distance cut-off for Lennard-Jones interactions was set to 10 Å. The production simulations were performed in constant volume ensemble and temperature was regulated by Langevin thermostat^88^ (see Supporting Information of Ref. ^18^ for details about minimization and equilibration protocols). The production simulations were run for 20 μs and either 10 or 5 independent trajectories were obtained for each tested *ff*. In a few cases, we used trajectories from our previous papers (Table S2 in Supporting Information).

DESAMBER, BSFF1 and CHARMM36 simulations were performed in Gromacs2020.^89^ We used the recommended TIP4P-D, TIP3P and CHARMM-modified TIP3P (TIP3P_CHARMM_) water models for DESAMBER, BSFF1 and CHARMM36 simulations, respectively. KCl ions described by charmm22, Joung&Cheatham, and CHARMM parameters for DESAMBER, BSFF1 and CHARMM36 simulations, respectively, were added to neutralize the system and to establish excess-salt ion concentration of 0.15 M. We note that the simulation protocol in Gromacs2020 slightly differed from the one in AMBER18 due to differences in simulation codes. Namely, Gromacs simulations were performed in a rhombic dodecahedral box and bonds involving hydrogens were constrained using the LINCS algorithm.^90^ The cut-off distance for the direct space summation of the electrostatic interactions was 10 Å and the simulations were performed at 298 K using the stochastic velocity rescale thermostat^91^ (see Table 1 for summary of all tested *ff*s, water models and ion parameters).

### System preparation and simulation protocols for polarizable CHARMM_Drude_ and AMOEBA *ff*s

Ten pre-equilibrated structures of 14-mer UUCG TL prepared by the nonpolarizable OL3_CP_ *ff* were used as starting points for CHARMM_Drude_^92–95^ and AMOEBA^58, 97^ simulations. For the CHARMM_Drude_ *ff*, the structures were transformed into the polarizable model using the CHARMM software (version 44b1).^74^ During the conversion process, Drude particles were introduced for all heavy atoms and the lone pairs associated with each hydrogen acceptor. The OPC water molecules were converted into the polarizable SWM4-NDP model.^96^ After initial minimization and equilibration procedure using the NAMD 2.13 package,^98^ ten independent production simulations were performed at 298 K in OpenMM 8.0^99^ for the length of 5 μs. Drude Langevin integrator^100, 101^ was utilized with a timestep of 1 fs. The pressure was maintained at 1 bar utilizing the Monte Carlo barostat.^102^ The covalent bonds involving hydrogens were kept rigid using the SHAKE^103^ and SETTLE^104^ algorithms for solute and waters, respectively. A constraint of 0.2 Å was applied to limit the length of Drude-nuclei bonds. Electrostatic interactions were treated using the Particle-Mesh Ewald method (PME)^105^ with a 12 Å cutoff for the real space term. Non-bonded interactions were truncated at 12 Å using a switching function from 10 to 12 Å.

For the AMOEBA *ff*, pre-equilibrated UUCG TL structures were transferred into xyz files and minimized in 10 000 steps using the steepest descent method. The systems were then heated up to 298 K and equilibrated for the pressure of 1 bar. Stochastic velocity rescale thermostat^91^ and Monte Carlo barostat^102, 106^ with coupling constants of 0.1 ps were used to maintain temperature and pressure, respectively. The applied real-space cutoff for electrostatics and van der Waals was 7 and 12 Å, respectively. RESPA integrator^107^ was used with the integration step of 1 and 2 fs for equilibration and production simulations, respectively. Other control functions and parameters were set to their default values. Ten independent simulations were run for 5 μs in NVT ensemble using the GPU-accelerated Tinker code^108^ (see Table 1 for summary of all tested pair-additive and polarizable *ff*s).

### Enhanced sampling folding simulations

We used a combination of well-tempered metadynamics (MetaD)^109–111^ and replica exchange with solute tempering (REST2),^112^ i.e., the ST-MetaD method,^61^ for folding simulations of the r(gcUUCGgc) 8-mer TL. ST-MetaD simulations were performed with 12 replicas starting from unfolded single strands and were simulated in the effective temperature range of 298−497 K. We performed typically three independent ST-MetaD simulations for the tested *ff*s. The average acceptance rate was ∼30%. The *ε*RMSD metric^113^ was used as a biased collective variable using the reference native TL structure. ST-MetaD simulations were carried out using a GPU-capable version of GROMACS2018^89^ in combination with PLUMED 2.5^114, 115^ and run for 5 μs per replica; see Ref. ^61^ for further details about the ST-MetaD protocol and Table S2 in Supporting Information for a full list of standard and enhanced sampling simulations.

### Comparison with NMR data and Folding Experiments

Besides detailed analysis of the structural developments in the simulations of the 14-mer UUCG TL, the structural ensembles were also compared with the available primary NMR data.^34, 48^ We analyzed four NMR observables, i.e., (i) backbone 3J scalar couplings, (ii) sugar 3J scalar couplings, (iii) nuclear Overhauser effect intensities (NOEs), and (iv) ambiguous NOEs (ambNOEs) resulting from a sum of overlapping peaks. 3J scalar couplings were calculated via Karplus relationships, NOEs were obtained as averages over the N samples, and ambNOEs were calculated by summing the contribution from either two, three or four nuclei pairs and again averaged over the N samples (see Ref. ^19^ for details). Combination of all those analyzed NMR observables (calculated as weighted arithmetic mean) provided the total χ^2^ value for each MD simulation. In principle, the lower the total χ^2^ value, the better the agreement between experiment and predicted data from MD simulation. However, as most of considered NMR signals are either intranucleotide and/or from stem residues, local deviations of loop residue(s) from native conformation were typically not visibly reflected by the χ^2^ analysis (see the paragraph “Interpretation of Measured NMR signals of the 14-mer UUCG TL is not straightforward” in Results and Discussions).

ST-MetaD simulations of the 8-mer UUCG TL provided populations of the native structure and other conformations, which can be used for estimation of the folding free energy (ΔG°_fold_)^61^ and directly compared with available experimental data.^31–33^ Reference native structure of the 8-mer UUCG TL was taken from our previous work.^18^ The *ε*RMSD threshold separating the folded and un(mis)folded states was set at value of 0.7 (see Ref. ^61^ for details about ΔG°_fold_ estimations and convergence).

### Analysis and Definition of Important states

We identified number of alternative (non-native) substates that are significantly populated during MD simulation with certain *ff*s instead of the native TL state; see the paragraph “Description of alternative (misfolded) states of the UUCG TL identified in MD simulations” in Results and Discussions. Those were identified by visual inspection of all trajectories via VMD^116^ and PyMOL^117^ and characterized in detail by CPPTRAJ.^118^ Coordinates of characterized misfolded states are attached as PDB files in Supporting Information.

## RESULTS and DISCUSSION

We performed an extended set of standard MD simulations of one of the most common RNA benchmark structures, the 2KOC NMR structure^34^ of the 14-mer UUCG TL with five-base pair stem, using variety of RNA *ff*s including polarizable ones. We then carried out a set of folding (biased) simulations for selected *ff*s (those best performing for the folded TL) using the 8-mer UUCG TL. Finally, we ran standard simulations of some other systems, to better understand properties of specific *ff*s. In total, we analyzed 120 standard MD simulations of the UUCG 14-mer with a cumulative time of 2100 μs, fifteen ST-MetaD simulations of the UUCG 8-mer with 12 replicas and cumulative time of 900 μs, and additional 63 simulations of other systems with a cumulative time of 368 μs, resulting in almost 3.4 ms of simulation data (Tables S2 and S3 in Supporting Information). We first introduce the native UUCG TL structure with its key interactions and define all alternative states that are sampled in the simulations. Next, we describe the most common disruption and (in few cases) refolding pathways of the UUCG TL in simulations with different *ff*s. Then, we present UUCG 8-mer folding simulations for three *ff*s which are best performing in the standard simulations. Since we obtained striking differences in folding free energies among these *ff*s, we further inspect their performance using standard simulations for additional nucleic acid systems. Finally, we provide brief synthesis of the data and our perspective on the current state of the art and future development of RNA *ff*s.

### The native structure of UUCG TL

The 2KOC NMR structure^34^ contains five base-pair canonical stem (G_1_C_14_, G_2_C_13_, C_3_G_12_, A_4_U_11_ and C_5_G_10_ base pairs) and four U_6_, U_7_, C_8_ and G_9_ loop nucleotides (Figure 1). The UUCG TL *native* state contains *trans*-wobble G_9_U_6_ base pair which requires G_9_ to adopt an unusual *syn* conformation. The *trans*-wobble G_9_U_6_ base pair is stabilized by a G_9_(N1H)…U_6_(O2) H-bond and two U_6_(2’-OH)…G_9_(O6) and U_7_(2’-OH)…G_9_(N7) sugar – base H-bonds (Figure 1).^34, 55–57^ The U_7_ nucleotide is flipped out into solution and thus expected to be mobile. C_8_ is interacting with a U_6_ phosphate moiety and forms a C_8_(N4H)…U_6_(*pro*-R_P_) H-bond (the base – phosphate interaction type 7; 7BPh).^69^ The U_7_ and C_8_ nucleotides are also characterized by the unusual south-type ribose conformations (C2’-endo pucker; Figure 1).^34, 55–57^

**Figure 1:**
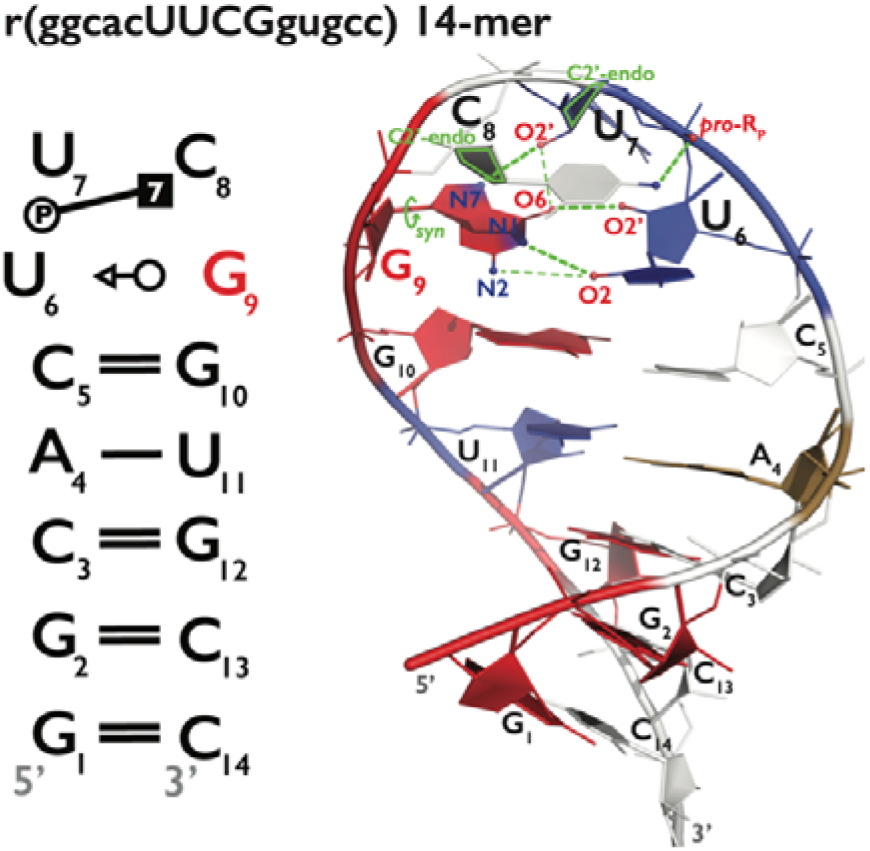
Secondary and tertiary structure of the 14-mer UUCG TL. Secondary structure annotation (left panel) follows the Leontis−Westhof nomenclature^119^ and residue labelling is consistent with the experimental structure (PDB ID 2KOC^34^). Both 7BPh interaction^69^ and key G_9_ nucleotide with *syn* conformation of *χ*_G9_ dihedral are highlighted. Panel on the right shows starting snapshot for MD simulations. A, C, G and U nucleotides are colored in sand, white, red, and blue, respectively. H- atoms, ions and water molecules are not shown for clarity. Key structural features of UUCG TL are colored in green and include *syn*-conformation of *χ*_G9_ dihedral and C2’-endo pucker of U7 and C8 riboses. Green dashed lines show the native H-bonds with involved heavy atoms explicitly labelled, i.e., G_9_(N1H)…U_6_(O2), U_6_(2’-OH)…G_9_(O6), U_7_(2’-OH)…G_9_(N7) and C_8_(N4H)…U_6_(*pro*-R_P_). Thin green dashed lines indicate two non-native H-bonds, i.e., G_9_(N2H)…U_6_(O2) and U_7_(2’-OH)…G_9_(O6) that are frequently established during MD simulations.

In previous simulation studies, we observed two alternative G_9_(N2H)…U_6_(O2) and U_7_(2’- OH)…G_9_(O6) H-bonds, which were often established when the native G_9_(N1H)…U_6_(O2) and U_7_(2’- OH)…G_9_(N7) H-bonds were weakened or broken (Figure 1). Those alternative H-bonds appear to assist in stabilization of the highly mobile G_9_ in the binding pocket and to maintain its *syn* conformation.^45^

Here, we used the following criteria to determine that the UUCG TL is sampling the *native* state: all four native G_9_(N1H)…U_6_(O2), U_6_(2’-OH)…G_9_(O6), U_7_(2’-OH)…G_9_(N7) and C_8_(N4H)…U_6_(*pro*-

R_P_) H-bonds are established with distance between proton donor and proton acceptor ≤ 3.5 Å; slightly higher cut-off of 3.7 Å was used for the C_8_(N4H)…U_6_(*pro*-R_P_) H-bond. Further, G_9_ is sampling the *syn* conformation defined by *χ*_G9_ dihedral G_9_(O4’)-G_9_(C1’)-G_9_(N9)-G_9_(C4) between -25° and 115°, Table 2).

**Table 2:** Summarized labelling and structural criteria used for characterization of alternative (misfolded) states occurring during MD simulations of the 14-mer UUCG TL.

| State | RMSD from start<sup>a</sup> | clustering criteria | Figure 2 label |
| --- | --- | --- | --- |
| *Native* | 0.9 ± 0.8 | G<sub>9</sub>(N1)-U<sub>6</sub>(O2) distance < 3.5 Å, U<sub>6</sub>(O2')-G<sub>9</sub>(O6) distance < 3.5 Å, U<sub>7</sub>(O2')-G<sub>9</sub>(N7) distance < 3.5 Å, C<sub>8</sub>(N4)-U<sub>6</sub>(*pro*-R<sub>p</sub>) distance < 3.7 Å; G<sub>9</sub>(O4')-G<sub>9</sub>(C1')-G<sub>9</sub>(N9)-G<sub>9</sub>(C4) dihedral <-25°,115°> | A) |
| *Sugar-base (N7) lost* | 1.0 ± 0.8 | at least one of G<sub>9</sub>(N1)-U<sub>6</sub>(O2) and G<sub>9</sub>(N2)-U<sub>6</sub>(O2) distances < 3.5 Å, U<sub>6</sub>(O2')-G<sub>9</sub>(O6) distance < 3.5 Å, U<sub>7</sub>(O2')-G<sub>9</sub>(N7) distance > 3.5 Å, C<sub>8</sub>(N4)-U<sub>6</sub>(*pro*-R<sub>p</sub>) distance < 3.7 Å, G<sub>9</sub>(O4')-G<sub>9</sub>(C1')-G<sub>9</sub>(N9)-G<sub>9</sub>(C4) dihedral <-25°,115°> | B) |
| *7BPh lost* | 1.0 ± 0.8 | at least one of G<sub>9</sub>(N1)-U<sub>6</sub>(O2) and G<sub>9</sub>(N2)-U<sub>6</sub>(O2) distances < 3.5 Å, U<sub>6</sub>(O2')-G<sub>9</sub>(O6) distance < 3.5 Å, U<sub>7</sub>(O2')-G<sub>9</sub>(N7) distance < 4.0 Å, C<sub>8</sub>(N4)-U<sub>6</sub>(*pro*-R<sub>p</sub>) distance > 3.7 Å, G<sub>9</sub>(O4')-G<sub>9</sub>(C1')-G<sub>9</sub>(N9)-G<sub>9</sub>(C4) dihedral <-25°,115°> | C) |
| *Both sugar-base (N7, O6) lost* | 1.1 ± 0.8 | at least one of G<sub>9</sub>(N1)-U<sub>6</sub>(O2) and G<sub>9</sub>(N2)-U<sub>6</sub>(O2) distances < 3.5 Å, U<sub>6</sub>(O2')-G<sub>9</sub>(O6) distance > 3.5 Å, U<sub>7</sub>(O2')-G<sub>9</sub>(N7) distance > 3.5 Å, C<sub>8</sub>(N4)-U<sub>6</sub>(*pro*-R<sub>p</sub>) distance < 3.7 Å, G<sub>9</sub>(O4')-G<sub>9</sub>(C1')-G<sub>9</sub>(N9)-G<sub>9</sub>(C4) dihedral <-25°,115°> | D) |
| *C<sub>8</sub> bulge + one sugar-base lost* | 3.0 ± 0.8 | at least one of G<sub>9</sub>(N1)-U<sub>6</sub>(O2) and G<sub>9</sub>(N2)-U<sub>6</sub>(O2) distances < 3.5 Å, at least one of U<sub>6</sub>(O2')-G<sub>9</sub>(O6) and U<sub>7</sub>(O2')-G<sub>9</sub>(N7) distances > 3.5 Å, C<sub>8</sub>(N4)-U<sub>6</sub>(*pro*-R<sub>p</sub>) distance > 5.0 Å, G<sub>9</sub>(O4')-G<sub>9</sub>(C1')-G<sub>9</sub>(N9)-G<sub>9</sub>(C4) dihedral <-25°,115°> | E) |
| *G<sub>9</sub> bulge (syn)* | 2.4 ± 0.6 | G<sub>9</sub>(N1)-U<sub>6</sub>(O2) distance > 5.0 Å, U<sub>6</sub>(O2')-G<sub>9</sub>(O6) distance > 5.0 Å, G<sub>9</sub>(O4')-G<sub>9</sub>(C1')-G<sub>9</sub>(N9)-G<sub>9</sub>(C4) dihedral <-25°,115°>, RMSD of loop nucleotides < 3.0 Å | F) |
| *G<sub>9</sub> bulge (anti, high-anti)* | 2.6 ± 0.7 | G<sub>9</sub>(N1)-U<sub>6</sub>(O2) distance > 5.0 Å, U<sub>6</sub>(O2')-G<sub>9</sub>(O6) distance > 5.0 Å, G<sub>9</sub>(O4')-G<sub>9</sub>(C1')-G<sub>9</sub>(N9)-G<sub>9</sub>(C4) dihedral either <-180°, -25°> or <115°,180°>, RMSD of loop nucleotides < 3.0 Å | G) |
| *G<sub>9</sub> back in pocket (anti, high-anti)* | $1.8 \pm 0.7$ | $G_9(N1)-U_6(O2)$ distance $< 5.2 \text{ \AA}$, at least one of the following two criteria was true: $U_6(O2')-G_9(O6)$ distance $< 3.7 \text{ \AA}$ and $C_8(N4)-U_6(pro-R_p)$ distance $< 4.2 \text{ \AA}$, $G_9(O4')-G_9(C1')-G_9(N9)-G_9(C4)$ dihedral either $<-180^\circ, -25^\circ>$ or $<115^\circ, 180^\circ>$, RMSD of loop nucleotides $< 3.0 \text{ \AA}$ | H) |
| *U<sub>6</sub> + U<sub>7</sub> + C<sub>8</sub> bulge* | $3.5 \pm 1.0$ | $G_9(N1)-U_6(O2)$ distance $> 5.0 \text{ \AA}$, at least one of $U_6(O2')-G_9(O6)$ and $U_7(O2')-G_9(N7)$ distances $< 4.0 \text{ \AA}$, $C_8(N4)-U_6(pro-R_p)$ distance $> 5.0 \text{ \AA}$; $G_9(O4')-G_9(C1')-G_9(N9)-G_9(C4)$ dihedral $<-25^\circ, 115^\circ>$ | I) |
| *Loop disrupted* | $4.2 \pm 1.2$ | RMSD of loop nucleotides $> 3.0 \text{ \AA}$; except when been included in the *U<sub>6</sub> + U<sub>7</sub> + C<sub>8</sub> bulge* state | J) |
| *Stem+loop disrupted* | $4.3 \pm 1.5$ | RMSD of all nucleotides $> 4.6 \text{ \AA}$ | K) |

<sup>a</sup> Averages (with errors as standard deviations) from all states belonging to the particular cluster from all simulations (considering all tested RNA *ffs*) are shown. RMSD was calculated for all heavy atoms of loop nucleotides, i.e., U<sub>6</sub>, U<sub>7</sub>, C<sub>8</sub> and G<sub>9</sub>. The equilibrated experimental structure (i.e., the starting MD snapshot) was used as the reference.

### Description of alternative (misfolded) states of the UUCG TL identified in MD simulations

Structural dynamics of the UUCG TL sampled by the tested RNA *ff*s was characterized by a variety of alternative (misfolded) loop conformations. We carefully inspected all MD trajectories and grouped these alternative conformations into ten different states (Table 2 and Figure 2). They were sampled during both disruption and refolding back to the *native* state of the UUCG TL. These states were ordered from those closest to the *native* state towards more disrupted structures.

**Figure 2:**
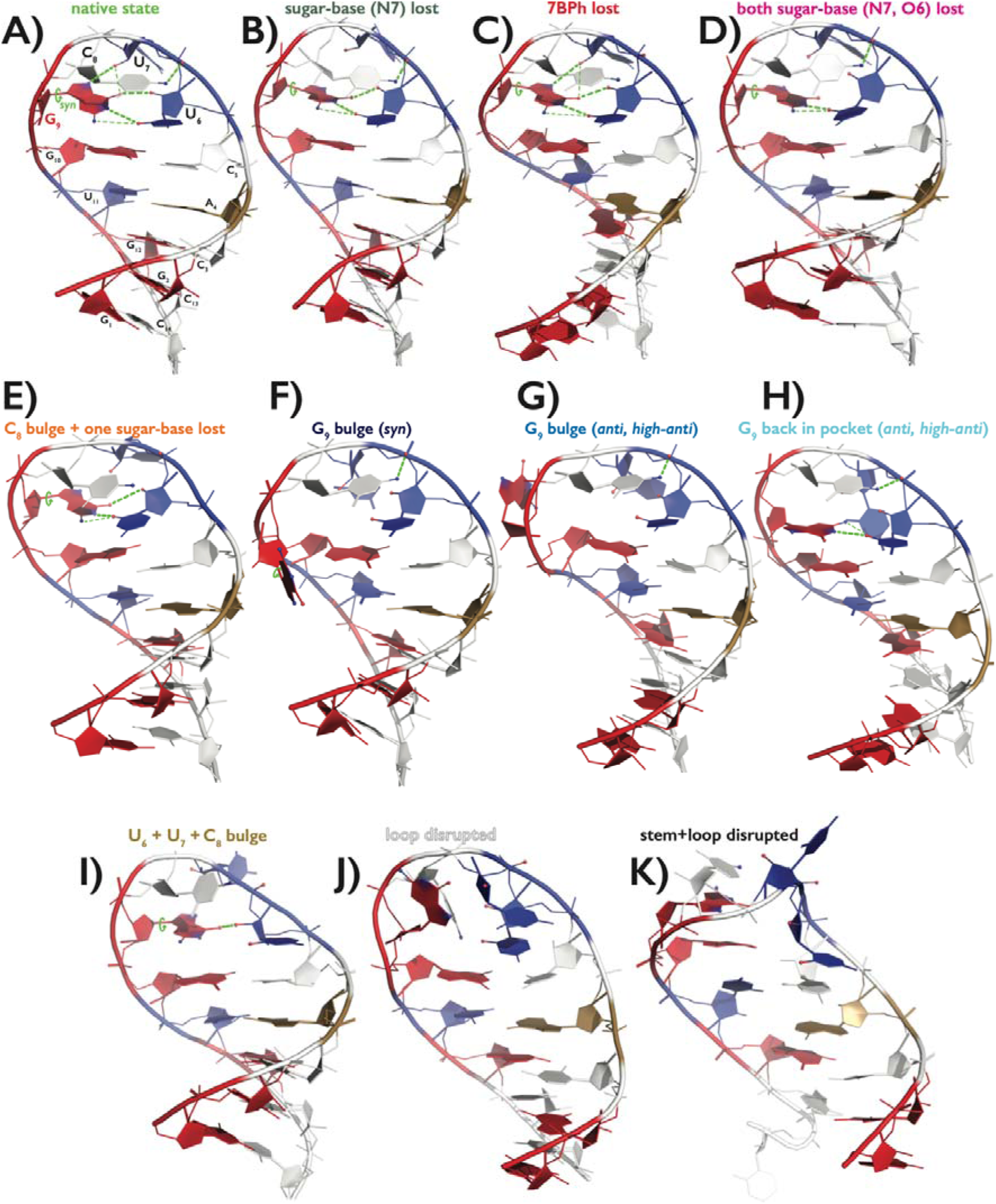
Representative snapshots of all important structural states frequently occurring during MD simulations of the 14-mer UUCG TL. See the legend to Figure 1 for nucleotide color scheme. Occurrence of key structural features that were monitored for characterization of each state, i.e., *syn* conformation of G_9_ nucleotide, G_9_(N1H)…U_6_(O2), G_9_(N2H)…U_6_(O2), U_6_(2’-OH)…G_9_(O6), U_7_(2’- OH)…G_9_(N7), and C_8_(N4H)…U_6_(*pro*-R_P_) H-bonds, is explicitly highlighted by green lines within each structure; see Table 2 for detailed description. Color labelling of the names of all states will be used later in the paper.

The first three states, labelled as *sugar-base (N7) lost*, *7BPh lost* and *both sugar-base (N7, O6) lost* were structurally close to the *native* state and their common feature was that the G_9_ remained in the binding pocket and maintained the *syn* conformation (Table 2 and Figure 2). Sampling of those states was usually reversible and most likely reflected the genuine dynamics within the loop due to thermal fluctuations. These states were also occurring as (final) intermediates upon successful refolding from disrupted structures back to the *native* state. Common feature of these states was the partial shift of G_9_ within the active side, characterized by formation of the alternative G_9_(N2H)…U_6_(O2) H-bond.^45^ The *sugar-base (N7) lost* state contained structures with the G_9_ in *syn* conformation, the *trans*-wobble G_9_U_6_ base pair formed (at least one of the native G_9_(N1H)…U_6_(O2) and alternative G_9_(N2H)…U_6_(O2) H- bonds was established) and the 7BPh interaction established by stable C_8_(N4H)…U_6_(*pro*-R_P_) H-bond. Further, the native U_6_(2’-OH)…G_9_(O6) sugar – base H-bond was formed but the other U_7_(2’- OH)…G_9_(N7) sugar – base H-bond was lost (the distance between U_7_(O2’) and G_9_(N7) being greater than 3.5 Å). The *7BPh lost* state also contained structures with G_9_ in *syn* and *trans*-wobble G_9_U_6_ base pair formed. For this state, both native U_6_(2’-OH)…G_9_(O6) and U_7_(2’-OH)…G_9_(N7) sugar – base H- bonds were formed while the 7BPh interaction was lost (i.e., the distance between C_8_(N4) and U_6_(*pro*- R_P_) exceeded 3.7 Å). Finally, the *both sugar-base (N7, O6) lost* state was also characterized by G_9_ in *syn* and *trans*-wobble G_9_U_6_ base pair formed. The 7BPh interaction was established but now both native sugar – base H-bonds were lost, i.e., the distances between the U_6_(O2’) and U_7_(O2’) hydroxyl oxygens and G_9_(O6, N7) acceptors were greater than 3.5 Å; Table 2).

Other four states, termed as *C_8_ bulge + one sugar-base lost*, *G_9_ bulge (syn)*, *G_9_ bulge (anti, high-anti)*, and *G_9_ back in pocket (anti, high-anti)* could be categorized as partial disruption of the loop (Table 2 and Figure 2). They involved a broader variety of conformations within each cluster with details often dependent on the particular *ff*. Generally, these states assembled structures with either C_8_ or G_9_ flipped out and other misfolded structures with G_9_ back in the binding pocket with nonnative *anti*/high*-anti* conformation of *χ*_G9_ dihedral. Occurrence of these states during MD simulation could be followed by further structural deformations with complete loss of the native UUCG TL fold. The *C_8_ bulge + one sugar-base lost* state still included structures with G_9_ in *syn* and *trans*-wobble G_9_U_6_ base pair formed. However, the 7BPh interaction was lost, C_8_ more displaced (i.e., distance between C_8_(N4) and U_6_(*pro*-R_P_) was greater than 5.0 Å), and one of the native sugar – base H-bonds was lost, i.e., at least one of the distances between U_6_ (O2’) and G_9_(O6) and between U_7_(O2’) and G_9_(N7) was greater than 3.5 Å. The *G_9_ bulge (syn)* state was composed from structures with G_9_ in *syn* conformation but repelled from the binding pocket, i.e., both distances between G_9_(N1) and U_6_(O2) and between U_6_(O2’) and G_9_(O6) were greater than 5.0 Å while RMSD of loop nucleotides (involving all heavy atoms from U_6_, U_7_, C_8_ and G_9_ nucleotides) was less than 3.0 Å. Comparably, the *G_9_ bulge (anti, high-anti)* state also involved structures with G_9_ repelled from the binding pocket (with similar criteria as for the *G_9_ bulge (syn)* state) but G_9_ was sampling either *anti* or *high-anti* values of *χ*_G9_ dihedral. Note that both the *G_9_ bulge (syn)* and *G_9_ bulge (anti, high-anti)* states included structures independently of the status of the 7BPh interaction (i.e., both states contained structures where the 7BPh interaction could be both established and weakened/disrupted). The *G_9_ back in pocket (anti, high-anti)* state encompassed variety of structures, where G_9_ approached back into the binding pocket but in *anti*/high*- anti* orientation of *χ*_G9_ dihedral, distance between G_9_(N1) and U_6_(O2) was less than 5.2 Å, at least one of distances between U_6_ (O2’) and G_9_(O6) and between C_8_(N4) and U_6_(*pro*-R_P_) was less than 3.7 Å and 4.2 Å, respectively, and RMSD of loop nucleotides was less than 3.0 Å. G_9_ could reform the base pair with U_6_ but in a variety of nonnative orientations, including formation of nonnative *trans* Watson-Crick/Hoogsteen^119^ GU base pair with the formation of U_6_(N3H)…G_9_(O6/N7) H-bond. Majority of structures belonging to this state had at least one of the 7BPh interaction and the U_6_(2’-OH)…G_9_(O6) H-bond established.

Remaining three characterized states, termed as *U_6_ + U_7_ + C_8_ bulge*, *loop disrupted*, and *stem+loop disrupted*, implicated more severe disruptions of the loop (Table 2 and Figure 2). The *U_6_ + U_7_ + C_8_ bulge* state involved a broad variety of structures where G_9_ maintained the *syn* conformation and remained in the binding pocket but the *trans*-wobble G_9_U_6_ base pair was completely disrupted (the distance between G_9_(N1) and U_6_(O2) was grerater than 5.0 Å). The 7BPh interaction was lost as the C_8_ was dislocated (the distance between C_8_(N4) and U_6_(*pro*-R_P_) was greater than 5.0 Å) and at least one of distances between U_6_(O2’) and G_9_(O6) and between U_7_(O2’) and G_9_(N7) was less than 4.0 Å. In summary, this state involved quite disrupted loop structures with the G_9_ maintaining its native conformation, often stabilized by one native sugar – base interaction. The *loop disrupted* state was defined just by RMSD of loop nucleotides greater than 3.0 Å except of structures that were already characterized, i.e., there was a partial overlap between this state and the previous one as some structures belonging to the *U_6_ + U_7_ + C_8_ bulge* state could also have RMSD of loop nucleotides greater than 3.0 Å. Some structures belonging to the *loop disrupted* state could have one native interaction established (typically the 7BPh interaction) as only RMSD value was used for definition of structures belonging to this state. Finally, the *stem+loop disrupted* state involved structures with RMSD of all residues greater than 4.6 Å (including all heavy atoms from the 14 nucleotides). This criterion overrode all other state definitions and includes structures not only with full displacement of loop nucleotides but usually also those where the canonical loop-closing C_5_G_10_ base pair (or even more stem base pairs) was disrupted.

### OL3_CP_ with gHBfix potentials, DESRES, DESAMBER and OL3_R2.7_ *ff*s revealed fully or reasonably stable UUCG TL dynamics

The standard^65^ OL3 RNA *ff*^27–30^ (including its versions with modified phosphate oxygens^66^)^38, 67^ struggles with structural description of the UUCG TL, resulting in irreversible loss of the native state typically within first tens to hundreds of ns of standard simulations.^15, 18, 59, 72^

Significant improvement was achieved in simulations combining OL3 with gHBfix potentials (Figure 3). We here compared the behavior of gHBfix21,^54^ gHBfixUNCG19^45^ and gHBfix19,^18^ all applied in combination with the OL3_CP_^27–30, 66, 67^ RNA *ff* variant. OL3_CP_–gHBfix21 simulations revealed entirely stable dynamics of the UUCG TL.^54^ The *native* state was lost in two out of ten OL3_CP_– gHBfix_UNCG19_ simulations and in all ten OL3_CP_–gHBfix19 simulations on the 20 μs-long timescale (Figure 3). We also tested charmm22 ion parameters (see Ref. ^80^ for the explanation) in OL3_CP_– gHBfix19 simulations (see Methods and Table 1). The overall sampling of the *native* state was still rather unsatisfactory, although with one refolding event and one stable trajectory (Figure 3). A common feature of OL3_CP_–gHBfix simulations was that sampling of the *native* state was accompanied with short reversible occurrences of the *sugar-base (N7) lost* and *7BPh lost* states, i.e., reversible weakening of the native U_7_(2’-OH)…G_9_(N7) and C_8_(N4H)…U_6_(*pro*-R_P_) H-bonds. Reversible distortions of the loop occurred via flip of either C_8_ (from the *7BPh lost* state to the *C_8_ bulge + one sugar-base lost* state) or G_9_ (from the *G_9_ bulge (syn)* state to the *G_9_ bulge (anti, high-anti)* state) out of its binding conformation; refolding back to the native state was possible (Figure 3). Note that the occurrence of the *C_8_ bulge + one sugar-base lost* state at the very end of sim #7 with the gHBfix21 was reversible and we actually observed refolding back to the native state at ∼20.4 µs upon prolongation of the simulation.^54^

**Figure 3:**
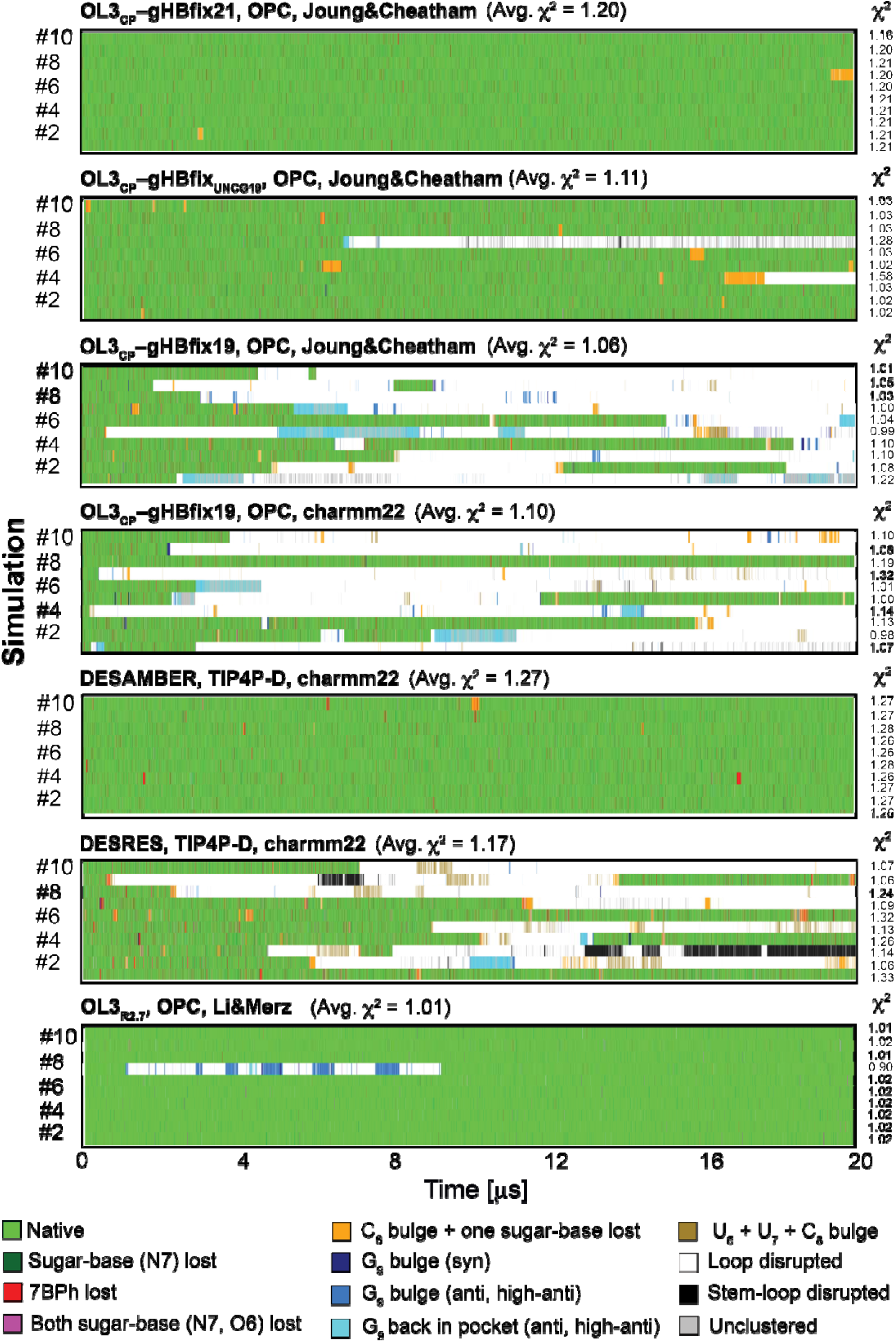
Structural dynamics of the 14-mer UUCG TL obtained by various RNA *ff*s in 20 μs-long standard MD simulations. Each panel is labelled by the applied *ff*, water model and ion parameters (see Table 1). Panels show evolution of characterized states highlighted by different colors; see Figure 2 for representative snapshots and Table 2 for detailed description. Comparison with the NMR data is shown next to each tested RNA *ff* as total *χ*^2^ values, see Methods. They were calculated for each simulation (values on the right) and as averages (over ten simulations as standard deviations).

Simulations with DESRES *ff* provided quite good description of the UUCG TL (Figure 3).^80^ Comparably to OL3_CP_–gHBfix19 simulations, loss of the *native* state occurred via flip of either C_8_ or G_9_ out of its binding conformation, i.e., via *C_8_ bulge + one sugar-base lost*, *G_9_ bulge (syn)* or *G_9_ bulge (anti, high-anti)* states. Refolding back to the *native* state was also possible and four out of 10 simulations maintained the native state after 20 μs (Figure 3). DESAMBER *ff* achieved fully stable dynamics of the 14-mer UUCG TL, where (similarly to the OL3_CP_–gHBfix21 simulations) sampling of the dominant *native* state was alternating with very short visits of the *sugar-base (N7) lost* and *7BPh lost* states (Figure 3).

Finally, fully stable structural description of the UUCG TL was also achieved with the OL3_R2.7_ *ff*. We evidenced one flip of G_9_ out of the binding pocket in sim #7, followed by sampling of *G_9_ bulge (anti, high-anti)*, *G_9_ back in pocket (anti, high-anti)* and *loop disrupted* states. This lasted for ∼8 μs, but then the *syn* conformation of *χ*_G9_ dihedral was restored, G_9_ returned to the binding pocket and the *native* state was neatly re-established (Figure 3). Interestingly, the *7BPh lost* state, i.e., the reversible weakening of the native C_8_(N4H)…U_6_(*pro*-R_P_) H-bond (Figure 2) occurred significantly less frequently than in simulations with OL3_CP_–gHBfix21 and DESAMBER *ff*s (both the latter having comparable performance; Figure 3). Hence, the 7BPh interaction in the UUCG TL is probably indirectly stabilized by the OL3_R2.7_ *ff* due to reduction of –CH…O– steric conflicts in the loop.^59^

In summary, OL3_CP_–gHBfix21, DESAMBER and OL3_R2.7_ *ff*s provide entirely stable structural description of the UUCG TL during our standard 10 × 20 µs simulations.

### Other pair-additive RNA *ff*s provided unstable description of the UUCG TL

Figure 4 summarizes data for the other pair-additive RNA *ff*s, namely PAK, Chen&Garcia, ROC, BSFF1, and CHARMM36 *ff*s. Simulations with PAK *ff* revealed initially reasonable structural description of the UUCG TL, but the *native* state was eventually lost in all five 20 μs-long simulations (Figure 4). Although the overall sampling of structural states was comparable with OL3_CP_–gHBfix19 and DESRES *ff*s (discussed in previous paragraph), we did not see any successful refolding back to the *native* state from more disrupted states, i.e., those where G_9_ left the binding site (Figure 4).

**Figure 4:**
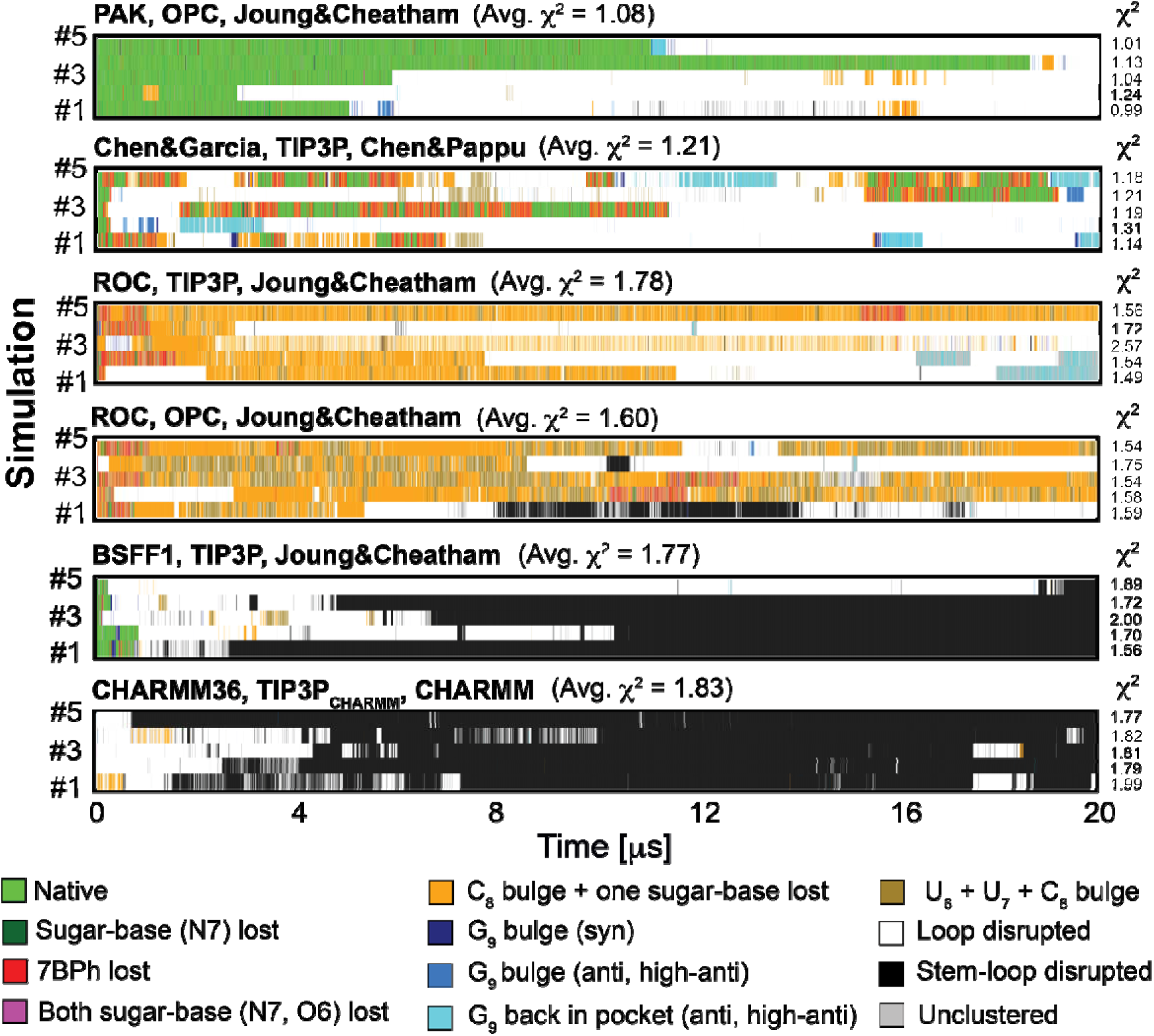
Structural dynamics of the 14-mer UUCG TL obtained by another set of pair-additive RNA *ff*s. Each panel shows evolution of the characterized states in five 20 μs-long standard MD simulations. See legend of Figure 3 for more details.

Simulations with Chen&Garcia *ff* provided strikingly different outcome in comparison with all other pair-additive *ff*s. The overall population of the *native* state was rather low (in agreement with the previous study using the shorter 10-mer UUCG TL).^15^ The *native* state could be reestablished rather easily from a variety of more disrupted structures but was then again swiftly lost. We observed major dynamics of the loop characterized by sampling of all the classified misfolded states and additional ones belonging to the *loop disrupted* state. In another words, UUCG TL simulated by the Chen&Garcia *ff* was characterized by sustained dynamics of loop nucleotides coupled with stable base pairing within the stem. Perturbation of the *native* state was typically initiated via loss of native C_8_(N4H)…U_6_(*pro*-R_P_) H-bond (*7BPh lost* state) followed by flip of C_8_ out of the binding site (*C_8_ bulge + one sugar-base lost* state) and then typically reached states with complete repositioning of other loop nucleotides.

The ROC *ff* did not provide a good structural description of the UUCG TL, regardless of its combination with either TIP3P (as originally suggested)^41^ or OPC water (Table 1). In agreement with the original work^41^ we observed abrupt and irreversible loss of the *native* state within first few hundreds of ns. Disruptions started with the loss of native C_8_(N4H)…U_6_(*pro*-R_P_) H-bond (*7BPh lost* state) and were followed by repositioning of either one, two or all three pyrimidine loop nucleotides (U_6_, U_7_ and C_8_). Hence, the UUCG TL with ROC *ff* was characterized by frequent sampling of *C_8_ bulge + one sugar-base lost* and *U_6_ + U_7_ + C_8_ bulge* states (Figure 4) where native H-bonds were lost (Figure 2). Although we also detected states with G_9_ flipped out to solvent, G_9_ typically maintained its *syn* conformation of *χ*_G9_ dihedral and remained within the binding pocket despite loss of all native H-bonds (the *U_6_ + U_7_ + C_8_ bulge* state).

With BSFF1 *ff* the *native* state was lost in all five simulations within the first few hundreds of ns. This was followed by complete rearrangement of loop nucleotides with weakening of stem base pairs during later stages of the simulations (Figure 4). Loop disruption was initiated by reversible flips of the G_9_ out of the binding pocket (the *G_9_ bulge (syn)* state). Once G_9_ lost its *syn* conformation (the *G_9_ bulge (anti, high-anti)* state) additional irreversible repositioning of the remaining loop nucleotides occurred. Then the disruption propagated towards the stem where the closing C_5_G_10_ base pair was broken (i.e., the *stem-loop disrupted* state was sampled; Figures 2 and 4).

Significant UUCG TL deformations occurred with the CHARMM36 *ff*. We observed abrupt loss of all native interactions and flip of G_9_ out of the binding pocket within first few ns of simulations, just after the initial equilibrations (Figure 4). The relocation of G_9_ was immediately followed by loss of remaining native interactions and resulted in complete distortion of the native arrangement. We also detected frequent loss of canonical base pairing in the stem with possible occurrence of completely unfolded structures in later stages of the simulations.

In summary, our extensive testing revealed that the structural description of UUCG *native* state is a significant challenge for several RNA *ff*s. Apart from PAK *ff* which revealed some improvement in structural stability of the *native* state in comparison with standard OL3 *ff*, all remaining *ff*s summarized in Figure 4 were characterized by abrupt loss of the UUCG native arrangement and sampling of alternative states. We observed that only Chen&Garcia *ff* was able to intermittently reestablish the *native* state from alternative misfolded states though only for very short times (typically tens up to hundreds of ns). PAK, ROC, BSFF1 and CHARMM36 *ff*s were characterized by irreversible loss of the native arrangement. BSFF1 and CHARMM36 *ff*s even sampled structures with broken canonical base pairs within the stem during later stages of MD simulations.

### Polarizable *ff*s also face challenges in accurate description of the UUCG TL structure

We included two available polarizable RNA *ff*s in our testing set of simulations. Although both CHARMM_Drude_ and AMOEBA *ff*s are used with GPU-accelerated versions of particular MD engines, i.e., OPEN-MM and Tinker-HP, we still obtained at least an order of magnitude slower performance (using Nvidia RTX3080ti GPU cards) in comparison with pair-additive *ff*s utilized by either pmemd.cuda^85^ in AMBER or GPU accelerated GROMACS code. Thus, presented results by polarizable *ff*s are based on shorter 5 μs-long MD simulations.

Both CHARMM_Drude_ and AMOEBA *ff*s revealed comparable behavior in structural description of the UUCG TL characterized initially by reversible loss of the *native* state and frequent sampling of alternative *7BPh lost*, *C_8_ bulge + one sugar-base lost* and *U_6_ + U_7_ + C_8_ bulge* states (Figure 5), i.e., those states where G_9_ remained in the binding pocket in its native *syn* conformation of the *χ*_G9_ dihedral (Figure 2). In other words, reversible loss of the native state started with breakage of the native C_8_(N4H)…U_6_(*pro*-R_P_) H-bond (*7BPh lost* state). It was followed by disruption of native sugar – base interaction and further repositioning of either one, two or all three pyrimidine loop nucleotides (U_6_, U_7_, and C_8_). However, we did not see any successful refolding back to the *native* state (on the limited 5 μs- long timescale) by both polarizable *ff*s once G_9_ left the pocket and lost its *syn* conformation, which occurred in four and five simulations (out of ten) with CHARMM_Drude_ and AMOEBA *ff*s, respectively (Figure 5).

**Figure 5:**
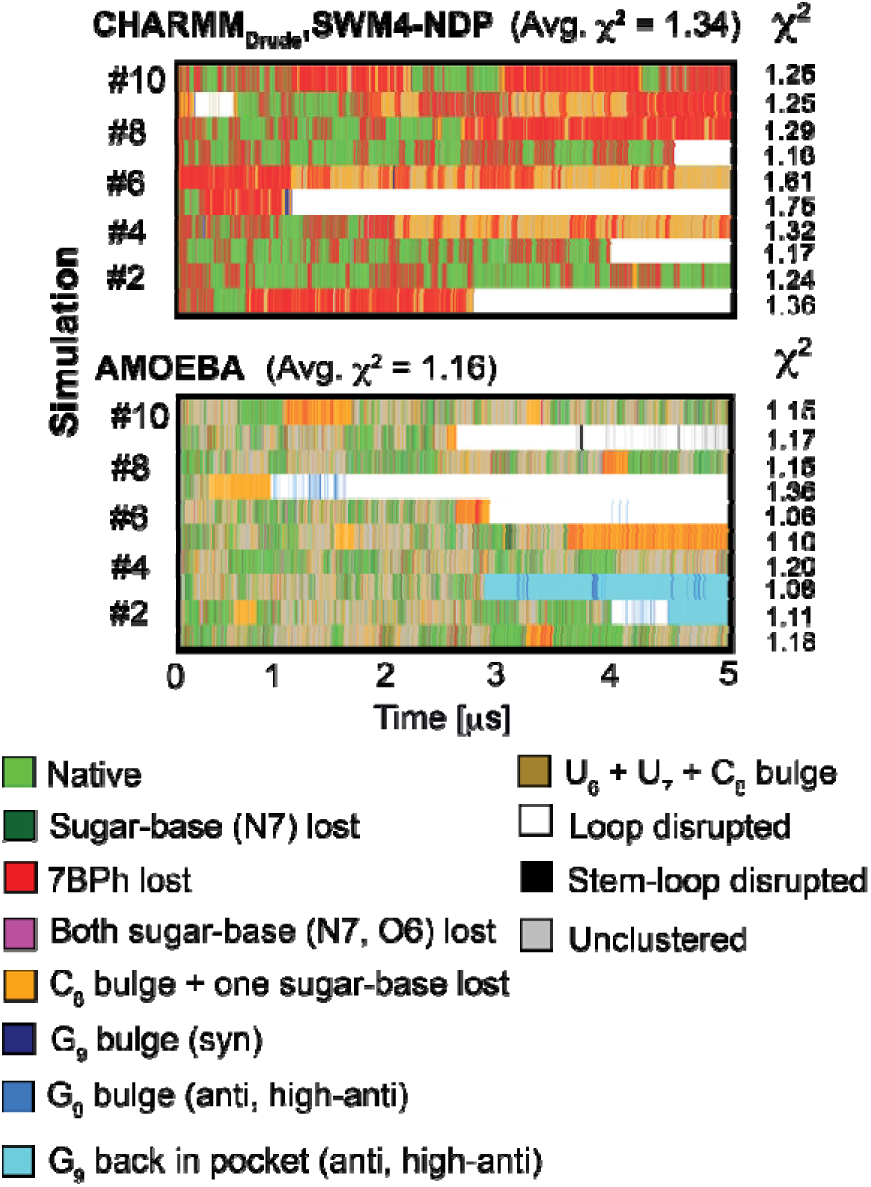
Structural dynamics of the 14-mer UUCG TL obtained by polarizable RNA *ff*s. Each panel shows evolution of characterized states in ten 5 μs-long simulations. See legend in Figure 3 for more details

Specific development was seen in sim #9 and sims #2 and #3 with CHARMM_Drude_ and AMOEBA *ff*s, respectively. We observed abrupt loop disruption during CHARMM_Drude_ sim #9 characterized by repositioning of all three U_6_, U_7_ and C_8_ pyrimidine loop nucleotides (from *7BPh lost*, via *C_8_ bulge + one sugar-base lost* towards *loop disrupted* state). However, G_9_ remained in position sampling its native *syn* conformation but U_6_ left the binding pocket and all H-bonds with G_9_ were broken. This unusual state falls rather ‘accidentally’ within clustering criteria of *G_9_ bulge (syn)* state (see dark-blue hit on Figure 5 and state definition in Table 2). As the G_9_ remained in the pocket in its *syn* state, refolding back to the *native state* was possible and, in fact, we detected successful refolding and complete reestablishment of all native interaction after ∼600 ns of MD sim #9. Nevertheless, *7BPh lost* and *C_8_ bulge + one sugar-base lost* states were dominantly sampled during later stages of this simulation (Figure 5).

Other unusual states were frequently sampled near the end of sims #2 and #3 using the AMOEBA *ff*. These states are marked as *G_9_ back in pocket (anti, high-anti)* state (see cyan marks in Figure 5 and clustering criteria in Table 2) and, indeed, are characterized by repositioning of G_9_ back to the binding pocket with noncanonical *anti/high-anti* conformation of *χ*_G9_ dihedral. Intriguingly, all native H-bond interactions were reestablished with G_9_ in *high-anti* state, which required visibly deeper insertion of G_9_ into the binding pocket, noncanonical C2’-endo sugar pucker of G_9_ and other major deformations along the sugar-phosphate backbone (see Figure S1 in Supporting information for direct comparison of the *native* state and this unusual ‘native-like’ state).

In summary, both tested polarizable RNA *ff*s struggled with description of the UUCG *native* state. We observed frequent refolding back to the *native* state from partially disrupted structures, but we did not detect any successful restoration of the native arrangement once G_9_ left the binding pocket and lost its canonical *syn* conformation. The *native* state was not dominantly populated as alternative loop arrangements were favored by both polarizable *ff*s.

### Interpretation of measured NMR signals of the 14-mer UUCG TL is not straightforward

Besides the detailed structure monitoring and clustering given above we also analyzed available 63 backbone 3J-couplings, 33 sugar 3J-couplings, 253 NOEs and 27 ambNOEs signals used to refine the NMR structure.^34, 48^ For each signal, we calculated differences between predicted (MD data) and experimentally observed values and all of them were subsequently combined (see Methods), which resulted in total *χ*^2^ value for each MD simulation (Figures 3, 4 and 5). Calculated *χ*^2^ values from all simulations by different RNA *ff*s were distributed between 0.90 to 2.57, where values around and below 1.0 usually indicate good agreement with the experiment (considering experimental errors). In general, *ff*s providing reasonably stable UUCG TL dynamics revealed *χ*^2^ values typically from ∼1.0 to ∼1.2 whereas those struggling to sample the *native* state provided higher *χ*^2^ values > ∼1.5. However, this was true for cases where larger structural distortions often affecting canonical base pairing in the stem were detected. On the other hand, we found out that extensive sampling of structures rather close to the *native* state (i.e., *sugar-base (N7) lost*, *7BPh lost* and *both sugar-base (N7, O6) lost* states) and even those considered as more disrupted states (including the *loop disrupted* state with RMSD of loop nucleotides greater than 3.0 Å; see Table 2) could also provide low total *χ*^2^ values. In fact, we evidenced several examples that irreversible loss of the *native* state and substantial sampling of states with disrupted native loop arrangement provided lower total *χ*^2^ values than simulations where the *native state* was the dominant sampled structure (Figures 3, 4 and 5). Furthermore, we observed cases where two simulations (even with the same *ff*) sampled comparable structural states (including the *native* state, close to native and more disrupted states) and revealed quite different total *χ*^2^ values (Figures 3, 4 and 5).

In summary, available NMR structural data for the UUCG TL must be used carefully for assessing performance of *ff*s during MD simulations. Disagreement between predicted and measured NMR signals revealed bigger structural problems in a simulation but did not correlate well with local dynamics within the loop, i.e., with (ir)reversible loss of the *native* state and sampling of alternative loop conformations. We identified that repositioning of one (or even two) loop nucleotide (i.e., minor loop distortion) resulted in just few violated NMR signals. Those discrepancies usually vanished upon averaging of the data using the entire set of 376 signals during calculation of the total *χ*^2^ value. Hence, our results confirm that the structural interpretation of NMR data for dynamic RNA molecules (even for small systems like the 14-mer UUCG TL) is rather complex.^48^ To precisely evaluate loop behavior during MD simulations of UUCG TL using NMR data, it is necessary to focus on specific NMR signals from loop nucleotides, rather than relying solely on a comparison between predicted and measured signals across all available data (the total *χ*^2^ value).

Folding simulations of the 8-mer UUCG TL revealed that the OL3_CP_–gHBfix21 *ff* currently provides the most accurate folding free energy estimate. The OL3_CP_–gHBfix21, DESAMBER and OL3_R2.7_ RNA *ff*s achieved a fully satisfactory performance for the structural description of the 14-mer UUCG TL in standard simulations. Thus, we used the 8-mer UUCG TL to test these three *ff*s using ST-MetaD folding simulations initiated from single-strand A-form conformation (see Methods). The 8- mer having an A-RNA stem with only two base pairs is an ideal target for such comparison as its structures in solution are in a dynamic temperature-dependent equilibrium between folded and unfolded states.^31–33^ The *ff*s should describe well not only the *native* state, but also its balance with the misfolded/unfolded ensemble.^61^ We performed three independent ST-MetaD simulations for all three tested *ff*s and obtained staggering difference between the calculated ΔG°_fold_ energies. Averages over the three simulations predicted ΔG°_fold_ of 0.0 ± 0.6 kcal/mol with OL3_CP_–gHBfix21, 2.4 ± 0.8 kcal/mol with DESAMBER and 7.4 ± 0.2 kcal/mol with OL3_R2.7_. Thus, the ΔG°_fold_ energy obtained with the OL3_CP_–gHBfix21 combination predicted the highest population of the native state at 298 K (Table 3) and the calculated ΔG°_fold_ energy is quite close to the available experimental data (reported values are between -1.3 and -0.7 kcal/mol).^31–33^ The present OL3_CP_–gHBfix21 result is comparable with a single simulation reported earlier (ΔG°_fold_ of 0.5 ± 0.1 kcal/mol).^54^ For overall comparison, we also calculated ΔG°_fold_ energies (single ST-MetaD runs) with two other *ff*s that provided rather reasonable description of the 14-mer UUCG TL. For the OL3_CP_–gHBfix_UNCG19_ *ff* we obtained ΔG°_fold_ of 1.9 ± 0.1 kcal/mol^61^ and for the DESRES *ff* 5.1 ± 0.1 kcal/mol (Table 3).

**Table 3:**
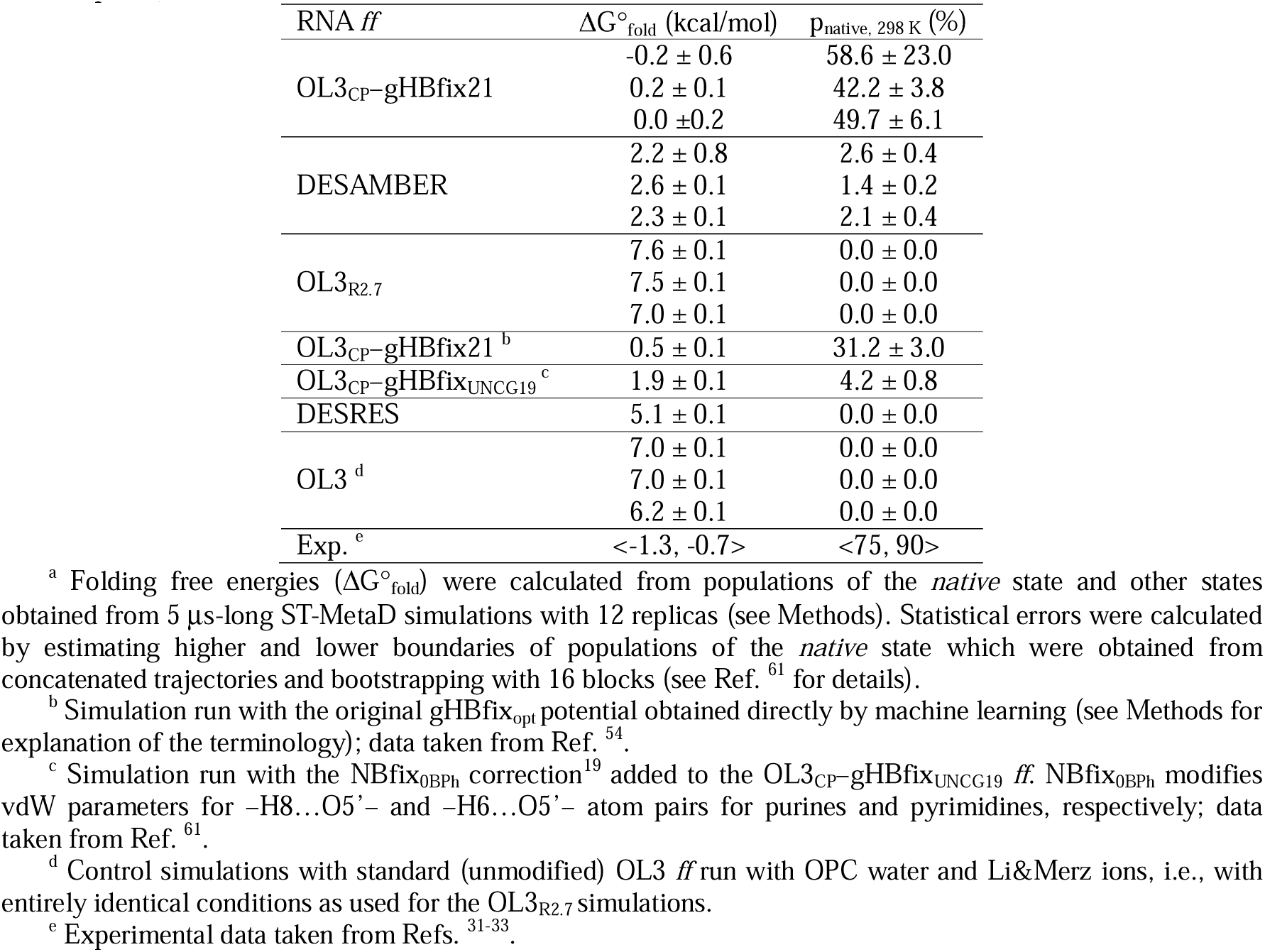
Calculated folding free energies (ΔG°_fold_) and expected populations of the *native* state at 298 K (p_native, 298 K_) for the 8-mer UUCG TL.^a^.

Taking all the data together, we observed strikingly different performance of the OL3_R2.7_ *ff* in standard simulations of the 14-mer and folding simulations of the 8-mer UUCG TL (Figure 3 and Table 3). To further inspect this point, we performed ST-MetaD simulations with the standard (unmodified) OL3 *ff* using identical water, ion and other setups as used with the OL3_R2.7_ *ff*. With OL3 we obtained ΔG°_fold_ of 6.7 ± 0.2 kcal/mol (average over three simulations, Table 3). Comparison of calculated ΔG°_fold_ energies obtained from OL3_R2.7_ and control OL3 simulations indicates that the NBfix modification of the –CH…O– interactions (the Hydrogen Repulsion Modification; HRM)^59^ does not bring any thermodynamic stabilization of the *native* state and could even have slightly opposite effect. To validate the convergence of estimated ΔG°_fold_ energies we monitored the number of folding events in continuous (demultiplexed) replicas from all ST-MetaD simulations and evidenced comparable behavior of folding events for all tested *ff*s (Figures S2-S16 in Supporting Information). Hence, we suggest that the unexpectedly high ΔG°_fold_ energies obtained with the OL3_R2.7_ *ff* are not caused by the lack of folding events, but rather reflect an imbalance between folded (native) and misfolded/unfolded ensemble, i.e., the *native* state is energetically disfavored in comparison with other states.

In summary, despite comparable results in standard simulations of the 14-mer UUCG TL (Figure 3), we obtained strikingly different outcome for the OL3_CP_–gHBfix21, DESAMBER and OL3_R2.7_ RNA *ff*s in folding simulations of the 8-mer UUCG TL. The OL3_CP_–gHBfix21 combination provided population of the *native* state rather consistent with experimental data. Obviously, this result should not be over-interpreted as folding energy of the UUCG TL was directly included in the training dataset of this *ff.* The DESAMBER *ff* underestimates the thermodynamic stability of the UUCG TL, however, in line with the standard simulations, it shows a clear improvement over the DESRES *ff*. In the DESAMBER paper the authors stated that RNA hairpins are thermodynamically under-stabilized by 3- 5 kcal/mol (see Supporting Information of Ref. ^70^) which is consistent with the data reported here. The discrepancy could be caused, for example, by underestimated stability of the A-RNA stem or by some other reasons.^14–16, 18, 39^ Interestingly, while the original DESRES *ff* paper suggests that thermodynamic stability of A-RNA duplexes is overestimated,^42^ the DESAMBER *ff* paper suggests that this *ff* should give rather correct thermodynamics for the A-RNA duplexes.^70^ The OL3_R2.7_ data is entirely counterintuitive (Table 3). While OL3_R2.7_ dramatically improves the *kinetic* stability of the folded UUCG TL compared to standard OL3 *ff*, it, at the same time, provides even slightly lower *thermodynamic* stability (higher ΔG°_fold_) than the basic OL3 (Table 3). It is a clear indication that the HRM introduced in Ref. ^59^ may over-stabilize some potentially spurious structures in the un(mis)folded ensemble, most likely via the modified –CH…O– interactions.

### Additional simulations with DESAMBER and OL3_R2.7_ *ff*s

The obtained data motivated us to perform additional simulations with DESAMBER and OL3_R2.7_ RNA *ff*s. See Supporting Information for full description of all these simulations.

Results from both standard MD and ST-MetaD simulations demonstrate that the DESAMBER *ff* is improved over its DESRES predecessor for the UUCG TL. It was shown in the original papers that both *ff*s provide in general good results for small RNA model systems.^42, 70^ However, independent testing revealed that DESRES^42^ *ff* struggles with some folded RNAs with tertiary interactions, namely, it collapsed structures of RNA kink-turn 7 (Kt-7) and L1 stalk ribosomal RNA (L1-stalk rRNA).^18^ Loss of the Kt-7 motif was also reported for the PAK *ff* (L1-stalk rRNA was not tested).^18^

Our evaluation of the DESAMBER *ff* gives the following picture. The DESAMBER *ff* still struggles with the kink-turn, as it has problems with the key A-minor tertiary interaction between the stems of the kink-turn (Figure S17 in Supporting information). It appears to ultimately lead to kink-turn unfolding and rearrangement into essentially an A-RNA-like structure with changed base pairing (Figure S18 in Supporting Information). However, there is visible improvement of DESAMBER *ff* over DESRES *ff* in description of the GA base pairs in the noncanonical stem of the Kt-7 and loss of the kinked structure is considerably slower compared to DESRES *ff*. On the other hand, for the L1- stalk rRNA the DESAMBER *ff* provides essentially the same results as DESRES *ff*, i.e., a swift collapse of the folded structure on ∼500 ns time scale (Figure S19 in Supporting information), indicating persisting problems with RNA tertiary interactions.

We then tested the OL3_R2.7_ *ff* (and other OL3-HRM setups; see Supporting Information)^59^ on several nucleic acids systems. For canonical A-RNA the OL3_R2.7_ *ff* caused moderate reduction of the A-RNA inclination and roll (Tables S4 and S5 in Supporting information). The OL3_R2.7_ *ff* did not improve the simulated ensembles of RNA tetranucleotides compared to standard OL3, i.e., it resulted into considerable sampling of spurious intercalated RNA structures (see Supporting Information for details).^15, 18, 37, 38, 43^ The OL3_R2.7_ *ff* further deepened the over-compaction of rU_5_ ssRNA compared to standard OL3 (Figure S20 in Supporting Information). Note that ssRNAs such as rU_5_ are known to be (severely) over-compacted when using the OL3 *ff*.^26, 120^ Thus, ssRNA-specific stafix *ff* modification has been parametrized to enable stable simulations of ssRNA/protein complexes.^26^ The compaction of rU_5_ by OL3_R2.7_ *ff* indicates that some additional imbalances are introduced by the HRM, likely due to sampling of over-stabilized –CH…O– interactions. This was ultimately confirmed by simulations of guanine quadruplexes (GQs). When simulating the Human Telomeric RNA (TERRA) GQ 3IBK^121^ with UUA propeller loops we observed formation of spurious –CH…O– interactions between the loop backbone and the GQ stem (Figures S21-S23 in Supporting information). Since the HRM should, in principle, be transferable also to the DNA *ff*s, we have further simulated analogous Human Telomeric (Htel) DNA GQ 1KF1^122^ with TTA propeller loop, combining HRM with the OL21^123^ DNA *ff*. The formation of spurious –CH…O– interactions was even more visible for DNA GQ and some such interactions occurred even within the GQ stem (Figures S24-S27 in Supporting information). We also tested “softer” version of the HRM using *R_i,j_* value of 2.8 Å^59^ for both RNA and DNA GQs (see Supporting Information for details), but the results remained virtually unchanged. In summary, the data indicate that while the HRM can ease steric constrictions in tightly packed parts of RNA structures, it can also lead to formation of networks of spurious –CH…O– interactions (H-bonds) in parts of RNA (and DNA) that have sufficient conformational freedom.

## CONCLUDING REMARKS

We provide a comprehensive assessment of contemporary RNA force fields (*ff*s) primarily using the challenging UUCG tetraloop (TL) RNA system.

Our study employs standard brute-force molecular dynamics (MD) simulations of the UUCG TL folded state to evaluate how well the *ff*s reproduce its native structure. Additionally, enhanced sampling folding simulations allow us to examine whether the *ff*s can accurately predict the ΔG°_fold_ free energies. As our results are based primarily on one specific RNA motif, our objective was not to declare a single ‘best’ RNA *ff*. Even within the UUCG TL system, the ranking of *ff*s differs depending on whether the evaluation is based on standard simulations of the folded state or on folding simulations from the unfolded state. The data indicate that rigorous testing of RNA *ff*s is a complex task that cannot be adequately addressed through a few anecdotal simulations or without an in-depth analysis of the trajectories.^15, 18^ In the future, we plan systematic tests of diverse *ff*s using additional benchmark systems, namely folded RNA structures with common tertiary interactions and protein-RNA complexes^124^. We also acknowledge that we were unable to comprehensively test all RNA *ff*s currently available in the literature (see, e.g., ^125–127^). However, any other RNA *ff* can be straightforwardly tested using the set of benchmark systems and calculations presented in this paper.

Most of the tested pair-additive and both polarizable *ff*s did not maintain the native structure of the UUCG TL. Intriguingly, we identified three recent pair-additive *ff* variants, i.e., OL3_CP_–gHBfix21,^54^ DESAMBER^70^ and OL3_R2.7_,^59^ which provided fully stable native conformation of the 2KOC 14-mer UUCG TL^34^ in 10 × 20 µs simulations. We subsequently tested these *ff*s in folding simulations of the 8-mer UUCG TL and obtained strikingly different results. While OL3_CP_–gHBfix21 predicted folding free energy quite consistent with experimental data, the DESAMBER and especially OL3_R2.7_ *ff* variants provided higher ΔG°_fold_ values.

The good performance of OL3_CP_–gHBfix21 was not unexpected given its machine learning-based development and inclusion of the UUCG TL in its training set.^54^ This *ff* has also been tested on a set of other RNA systems and so far no adverse effects of the gHBfix21 modification were found.^53^ However, it does not mean that problems will not emerge in future for insofar untested systems. The DESAMBER *ff* shows an improvement over its DESRES^42^ predecessor in both standard and folding simulations of the UUCG TLs but it has persistent issues^18^ with correct description of RNA kink-turn and L1-stalk rRNA systems, indicating imbalances in description of RNA tertiary interactions and potentially also some bias into the A-RNA conformation (see Supporting Information for details). The trickiest was analysis of the very recent OL3_R2.7_ *ff*. It introduces a substantial reduction of Lennard-Jones repulsion for numerous –CH…O– atom pairs using the NBfix approach^68^ (Hydrogen Repulsion Modification; HRM)^59, 128^ for the standard AMBER OL3 *ff*. It is based on an empirical observation that the compact folded stem-loop motifs contain multiple *very close* –CH…O– contacts which, when described by standard AMBER nonbonded parameters,^27^ leads to steric clashes (i.e., the *ff* hydrogens are ‘too large’).^19, 45, 59^ In addition, such close –CH…O– interactions are omnipresent in nucleic acids crystal structures.^59^ Indeed, the OL3_R2.7_ *ff* variant, which universally reduces the respective non-polar H…O *R_i,j_* Lennard-Jones parameters to 2.7 Å, neatly corrected standard simulations of the UUCG TL. However, the HRM did not improve the predicted UUCG TL ΔG°_fold_ value compared to the standard OL3 *ff*; the folding free energy in fact even slightly increased. More importantly, it can cause undesired side effects, i.e., the HRM can lead to formation of networks of spurious –CH…O– interactions in parts of RNA (and DNA) molecules that have sufficient conformational freedom (See Supporting Information for details).

The present results fully expose the difficulties in parametrization of RNA *ff*s. The HRM is based on very solid experimental and quantum chemical evidence.^59^ It works well for the folded UUCG TL, consistent with the findings of the original study,^59^ as some –CH…O– clashes indeed arise when the standard Lennard-Jones parameters^27^ are applied. In fact, we previously identified these clashes and performed tests on the UUCG TL using a reduction of vdW radii for non-polar hydrogens.^45^ Subsequently, we introduced a general NBfix correction for the intranucleotide 0BPh interaction,^69^ i.e., for the –H8…O5’– and –H6…O5’– atom pairs.^19^ In this correction, we opted for a conservative tuning of parameters to avoid potential negative effects on other RNA and DNA systems.^19^ Our present simulations show that besides positive effects for the folded UUCG TL, the HRM^59^ does not improve the description of the UUCG TL folding process and may even lead to issues when transferred to other systems. What is the primary cause of the limited transferability of HRM is presently not clear and will be further investigated. It could be due to the fundamental approximation of the basic Lennard-Jones plus atom-centered point charge model utilized by the contemporary pair-additive *ff*s, which affects especially the short-range repulsion region of the molecular interactions.^129, 130^ It limits direct transferability of quantum-chemical calculations of molecular interactions into the *ff* modifications.

In this context, the advantage of the basic OL3 *ff* is that it is not overparametrized in favor of specific systems and describes a relatively broad range of RNAs reasonably well. OL3 includes two key corrections of the original AMBER *ff*. The χ_OL3_ refinement^30^ prevents formation of spurious RNA ladder-like structures^131, 132^ which would otherwise constitute the global RNA structure minimum. The bsc0 α/γ refinement^29^ prevents occurrence of spurious γ-*trans* backbone states in A-RNA duplexes. Although the γ-*trans* states would not be dominantly populated without the bsc0 correction, they could be relatively long-living (dozens on ns) and reduce the A-RNA helical twist.^133^

The present data suggest that the introduction of any new RNA *ff* should ideally be accompanied by tests on a diverse set of RNA motifs, including single strands (tetranucleotides, hexanucleotides), tetraloops, A-RNA duplexes, kink-turn motifs, L1-stalk rRNA, and, where possible, other systems. Such testing would help evaluate how well the *ff* captures RNA structural dynamics across a broad spectrum of systems, highlighting its specific strengths and limitations. Given the inherent simplicity of the *ff* formalism, we remain skeptical that it is feasible to develop a universally accurate RNA *ff*. The significant van der Waals modification we recently had to implement for simulations involving protein-ssRNA complexes^26^ and opening-closing dynamics of DNA Holliday junction^134^ suggests that the complexity of nucleic acid structures, interactions, and dynamics is too vast to be comprehensively captured by a single *ff* variant. In the absence of a universal RNA *ff*, it can be justifiable to adapt atomistic *ff*s in a system-specific or goal-specific manner, even at the expense of full transferability, to address particular simulation challenges. There are certainly RNA systems that cannot be stably simulated by current *ff*s, even with system-specific adjustments.^23^ Perhaps, future developments in polarizable *ff*s may offer greater flexibility in parameterization, potentially allowing for a single *ff* variant to capture a broader range of RNA structural dynamics. However, current evidence, including the present results and other recent studies,^135–140^ suggests that achieving the right balance in polarizable nucleic acid *ff*s remains a formidable challenge.

In conclusion, while there have been significant advancements in RNA *ff* development, substantial hurdles still persist in achieving fully reliable and accurate modeling of the diverse structural dynamics of RNA systems.

## ASSOCIATED CONTENT

### Supporting Information

The Supporting Information is available free of charge via the Internet at http://pubs.acs.org/ and containing details about simulations of additional systems, supporting tables and figures to the article (PDF). Coordinates of the identified most common misfolded states of UUCG TL and examples of key sampled structural states of RNA and DNA GQs are also attached (zipped PDBs).

### Data Availability Statement

Precalculated MD datasets and NMR signals for UUCG TL are available at the URL: https://ida.4sims.eu.

## Supporting information

Supporting Information for the Article

## ACKNOWLEDGMENT

This work was supported by the Czech Science Foundation to V.M., M.K., P.S. and J.S. (grant number 23-05639S). This research also received the support of EXA4MIND, a European Uniońs Horizon Europe Research and Innovation programme under grant agreement N° 101092944 (M.O. and P.B). Views and opinions expressed are however those of the author(s) only and do not necessarily reflect those of the European Union or the European Commission. Neither the European Union nor the granting authority can be held responsible for them. P.K., M.P., M.O. and P.B. were also supported by ERDF/ESF project TECHSCALE (No. CZ.02.01.01/00/22_008/0004587).

