## Supporting Information for the Article for "Can We Ever Develop an Ideal RNA Force Field? Lessons Learned from Simulations of UUCG RNA Tetraloop and Other Systems"

### SUPPORTING RESULTS

**MD simulations of additional RNA and DNA motifs.** To get further insights into performance of selected *ff*s, we have carried out standard and enhanced sampling simulations for additional nucleic acids systems. These simulations are summarized in Table S3.

**Starting structures and simulation setup.** The initial coordinates of r(CAAU) and r(CCCC) tetranucleotides (TNs), and r(UUUUU) pentanucleotide (PN) were prepared using Nucleic Acid Builder of AmberTools14[1] as one strand of an A-form duplex. Starting topologies and coordinates of the remaining systems were taken from X-ray crystallography structures, and were prepared from the particular experimental structures by using the tLEaP module of AMBER22[2] program package (see Supporting Information of Ref. [3] for details about structure preparation). These systems include two canonical RNA duplexes (PDB ID 1QC0;[4] only ten canonical base pairs were considered, and 1RNA[5]), RNA Kink-turn 7 (Kt-7, PDB ID 1S72;[6] residues 76-83 and 91-101), and ribosomal L1 stalk RNA motifs from *Thermus thermophilus* (*T.t.*) and *Haloarcula marismortui* (*H.m.*); (L1-stalk rRNAs, PDB IDs 3U4M[7] and 5ML7[8]). Finally, two guanine quadruplexes (GQs) were simulated, namely the parallel-stranded human telomeric RNA and DNA GQs with propeller loops, PDB IDs 3IBK[9] and 1KF1[10], respectively. The 5-bromouracil in 3IBK[9] structure of RNA GQ was replaced by uracil.

RNA TNs, PN and duplexes were simulated with $OL3_{R2.7}$ *ff*,[11] whereas Kt-7 and L1-stalk rRNAs with DESAMBER *ff*.[12] Solvent models, ion concentrations and parameters, equilibration and simulation protocols were similar to those applied in UUCG TL simulations (see Methods and Table 1 in the main text). The RNA and DNA GQs were solvated in a truncated octahedral box of OPC[13] waters with the border of the box being at least 10 Å from the GQ. Channel $K^+$ ions present in the crystal structures of GQs were kept in place while additional $K^+$ and $Cl^-$ ions were added to neutralize the system and reach the final 0.15M KCl excess salt concentration using Li&Merz ion parameters.[14]

RNA and DNA GQs were first simulated using the standard RNA OL3[15-18] and DNA OL21[19] *ff*s, respectively, to obtain reference simulations. Then we have tested the effect of adding the NBfix van der Waals (vdW) adjustment for –CH…O– interactions (Hydrogen Repulsion Modification; HRM) suggested by Raguette et al.[11] We note that although the modification was originally proposed for RNA, it should be fully transferrable also to DNA molecules. When preparing the simulations we spotted a minor ambiguity of the HRM definition. According to the text of the original paper, Raguette et al. modified only minimum-energy distances of Lennard-Jones potential (i.e., the $R_{i,j}$ parameters) for the –CH…O– interactions.[11] In contrast to the text description, the published script for topology modification in Supporting Information of the original article[11] introduces also slight changes (decreases) of depths of the potential well ($\varepsilon_{i,j}$ parameters) for interactions involving nonbridging oxygens (O2 atom type) as they have slightly higher $\varepsilon$ parameter than the O3', O5' and O4' oxygens (OS atom types) in the standard AMBER *ff*.[15] Hence, we decided to test four HRM adjustments in simulations of RNA and DNA GQs: (i) the $OL3_{R2.7}$ and analogously modified $OL21_{R2.7}$ DNA variant (see below), (ii) the $OL3_{R2.8}$ and analogous $OL21_{R2.8}$ DNA variant, (iii) the $OL3_{R2.7.2}$ and $OL21_{R2.7.2}$ variants, which differ from the R2.7 modification by keeping the original $\varepsilon$ values for –CH…O– pairs as in the standard *ff*, and similarly, (iv) the $OL3_{R2.8.2}$ and $OL21_{R2.8.2}$, variants, which are like R2.8, but keep $\varepsilon$ values of the standard *ff*. Furthermore, our OL21 DNA implementation of the HRM encompasses the same hydrogens as the original modification of the OL3 RNA *ff*,[11] and in addition to that, it includes the second hydrogen on C2' (placed instead of the 2'-OH group in RNA), as well as the methyl hydrogens of thymine nucleotides.

All $OL3_{R2.7}$, $OL3_{R2.7.2}$, $OL3_{R2.8}$, $OL3_{R2.8.2}$, $OL21_{R2.7}$, $OL21_{R2.7.2}$, $OL21_{R2.8}$ and $OL21_{R2.8.2}$ simulations were performed in AMBER22,[2] whereas DESAMBER simulations were carried out in GROMACS2020.[20] All systems were simulated under standard MD simulation protocols (see Methods in the main text) with exception of RNA TNs, where we used the standard replica exchange solute tempering (REST2) protocol[21] executed at 298 K (the reference replica) with 8 replicas. The scaling factor (λ) values ranged from 1.0 to 0.601700871, corresponding to the effective solute temperatures ranging from 298 K to ~500 K. Further details about REST2 settings can be found elsewhere.[3] See Table S3 for summary of all additional MD simulations.

**Comparison with experiments and conformational analysis.** MD conformational ensembles from REST2 simulations of TNs were compared with available data from solution experiments.[22, 23] We used and analyzed separately four NMR observables, i.e., (i) backbone 3J scalar couplings, (ii) sugar 3J scalar couplings, (iii) nuclear Overhauser effect intensities (NOEs), and (iv) the absence of specific peaks in NOE spectroscopy (uNOEs). Their combination provided the total $\chi^2$ value (see Methods in the main text for more details). Dominant conformers occurring in reference (unbiased) replicas were identified by clustering based on an algorithm introduced by Rodriguez and Laio[24] in combination with the εRMSD metric[25] (see Ref. [3] for more details).

Structural behavior of r(UUUUU) PN in $OL3_{R2.7}$ *ff* was monitored by calculating the solvent-accessible surface area (SASA) and compared with control OL3 simulation. Behavior of folded RNA and DNA systems was assessed against their starting X-ray structures and/or compared against control simulations.

**Results of additional MD simulations with DESAMBER *ff*.** We have carried out an additional set of standard MD simulations with DESAMBER[12] *ff* using Kink-turn 7 (Kt-7) and ribosomal L1 stalk RNA (L1-stalk rRNA) motifs from *Thermus thermophilus (T.t.)* and *Haloarcula marismortui (*Table S3). Kink-turn is a recurrent RNA motif commonly found in ribosomes and some other folded RNAs. It induces a sharp bend, or kink, between two RNA helices (stems).[26] Kink-turns are characterized by highly conserved asymmetrical topology, where one strand is longer and forms a distinctive bulge of three or more nucleotides. This bulge is flanked by two stems: a non-canonical stem containing at least two adjacent *trans* Hoogsteen/Sugar-edge AG base pairs and a canonical stem formed by *cis* Watson-Crick GC base pairs. The sharp bend between the two stems is stabilized by an A-minor interaction, either type 0 or type I.[27] There is also a critically important interaction formed between the 2'-OH group of the first nucleotide in the bulge and the adenine in the AG base pair nearest to the kink. Overall, the global stability of the Kink-turn motif relies on simultaneous formation of multiple non-canonical RNA interactions, making it an excellent benchmark system for RNA *ff* testing. For our simulations, we selected the structure of Kink-turn 7 (Kt-7) from the large ribosomal subunit of *Haloarcula marismortui*. The same Kink-turn structure was used as a benchmark in our preceding papers.[3, 28] The Kt-7 sequence closely resembles a consensus Kink-turn motif, making it a suitable candidate for evaluating performance of RNA *ff*s. The earlier DESRES *ff* provided highly unstable trajectories for this system.[3]

We observed visible instability of the A-minor interaction in the Kt-7 simulations with the DESAMBER *ff*. All five independent simulations revealed that the starting A-minor I interaction transitioned into A-minor type 0 arrangement – a realistic development for the Kt-7 when taken out of the ribosomal context. However, the A-minor 0 interaction was subsequently quite unstable as we observed reversible disruptions of the A-minor 0 interaction (Figure S17). These disruptions started as soon as the A-minor 0 was first formed. Globally, the disrupted state was coupled with reduced degree of kinking. In other words, its presence caused Kt-7 to slightly straighten. In three (out of five) simulations, the structure subsequently fluctuated between the native A-minor 0 and a major population of the disrupted state for the rest of the simulations. In the remaining two simulations, the A-minor 0 instability and the associated reduced kink further progressed into a permanent degradation (unkinking) of the kink-turn motif (Figure S17). The unkinking events occurred at ~7.4 μs and ~4.2 μs in the respective simulations and the RNA molecule subsequently further (within few μs) rearranged into essentially an A-RNA-like structure, with changes of base pairing (Figure S18). The A-RNA-like structure was maintained till the end of both simulations. This is likely an unrealistic development which has no experimental support and indicates that the DESAMBER *ff* might be generally somewhat biased in favor of an A-form RNA. Given the clear and consistent instability of the A-minor 0 interaction, we expect that unkinking would eventually occur in the other three simulations on longer simulation timescales. We note that the A-minor interaction is the most prevalent RNA tertiary interaction in folded RNA molecules,[27] highlighting the importance of its accurate representation by RNA *ff*s. Nevertheless, the kink-turn description by the DESAMBER[12] *ff* is visibly improved compared to the DESRES[29] *ff*, since with DESRES we observed also instability of the non-canonical AG base pairs in the non-canonical stem while the disintegration of the kink-turn structure was much faster, occurring on a time scale of ~200 ns.[3] Still, despite the improvement, proper description of the key A-minor interaction remains challenging for the DESAMBER *ff* while the *ff* may also be somewhat biased towards the A-form RNA. Other tests will be needed to clarify the latter point.

The L1-stalk is a well-conserved and dynamic peripheral component of the large ribosomal subunit that aids in the release of deacylated tRNA from the ribosome during the elongation phase.[30, 31] Its rRNA part features numerous non-canonical interactions and important (recurrently occurring in folded RNA molecules) RNA motifs, making it another appealing benchmark system for *ff* evaluations. It is a very different RNA molecule compared to the commonly used RNA simulation benchmarks. Specifically, the L1-stalk rRNA contains two kink-turns, a TL, and a structured internal loop.[7] Collectively, these RNA motifs create a complex, six-layer nucleotide platform composed of stacked noncanonical base pairs and triplets. At the apex of this conserved platform lies the binding site for deacylated tRNAs.[32] Previous simulations demonstrated that the L1-stalk rRNA from *T.t.* can be stably modeled using the standard OL3 RNA *ff*[33] but not with the DESRES *ff* which caused its complete structural disintegration on the scale of hundreds of nanoseconds.[3] Thus, we tested performance of the DESAMBER[12] *ff* on the L1-stalk rRNA from *T.t.*. We once again observed rapid (timescale ~0.5 μs) large-scale structural disintegration of the L1-stalk rRNA structure with a loss of many RNA interactions, resulting in major unfolding of its characteristic structure in all four simulations (Figure S19). It is a nearly identical global structural degradation as previously observed with the DESRES[29] *ff*.[3] We subsequently tested the performance of DESAMBER *ff* on the L1-stalk rRNA from *H.m.*. Compared to the L1-stalk rRNA from *T.t.*, the one from *H.m.* contains a different TL (GCUA as opposed to UCCG), and one of the kink-turns (Kt-78) is heavily modified, with the AG base pairs typical for the motif replaced by different base pairs and triplets. The structural collapse of the L1-stalk rRNA from *H.m.* in all four simulations using the DESAMBER *ff* was even faster, on the scale of tens of nanoseconds. The loss of the rRNA fold was very similar to the *T.t.* structure albeit more extensive, with both kink-turns affected and the nucleotide platform damaged by the end of 1 μs-long MD simulations. In comparison, the L1-stalk rRNA from *H.m.* was entirely stable in all simulations using either the standard OL3 *ff* combined with SPC/E water model or the $OL3_{CP}$–gHBfix21 *ff* combined with OPC water model (and with the $NBfix_{0BPh}$ modification,[34] which, however is not assumed to affect these simulations significantly). For more details see Methods in the main text and Table S3. In summary, for structural description of the L1-stalk rRNAs, the performance of DESAMBER *ff* is not improved with respect to its DESRES predecessor.

**Results of additional MD simulations with $OL3_{R2.7}$ *ff* for single strand and duplex RNAs.** We tested the $OL3_{R2.7}$[11] *ff* for r(CAAAU) and r(CCCC) TNs, and r(UUUUU) PN. It was shown that both TNs have high propensity to form spurious intercalated structures with standard OL3 *ff*[23, 35, 36] that disagree with NMR data.[22, 23] In another study, the longer PN simulated with the standard OL3 *ff* sampled compacted structures with numerous intra-RNA interactions which are also deemed to be unrealistic.[37] Here, we performed REST2 simulations with $OL3_{R2.7}$ *ff* for the TNs and observed that the intercalated structure was the dominant conformation (~60%) sampled in the reference (unbiased) replica for both TNs (see, e.g., Ref.[38] for structural snapshots of intercalated states). This is comparable with what was reported for the standard OL3 *ff* (see, e.g., Table 6 in Ref. [3]). Dominant sampling of spurious intercalated structures with $OL3_{R2.7}$ *ff* disagrees with experiments, which is reflected by high calculated total $\chi^2$ values of 6.5 and 3.9 for r(CAAU) and r(CCCC), respectively (see Methods in the main text).

Since there are no NMR data for the longer r(UUUUU) PN, we instead calculated the solvent accessible surface area (SASA) during standard MD simulation with $OL3_{R2.7}$ *ff* and compared it with the control OL3 simulation. The $OL3_{R2.7}$ *ff* induced visibly higher compaction of the RNA compared to the standard OL3 *ff* (Figure S20). Note that the r(UUUUU) PN was shown to be excessively compacted already in simulations with the standard OL3 *ff* due to large-scale spurious RNA self-interactions, causing major issues in simulations of protein-RNA complexes involving ssRNAs.[37] The modified $OL3_{R2.7}$ *ff* further exacerbates the over-compaction.

We also tested the $OL3_{R2.7}$ *ff* on two RNA duplexes, i.e., $r(GCACCGUUGG)_2$ decamer (excised from the PDB ID 1QC0[4] structure) and $r(UUAUAUAUAUAUAA)_2$ tetradecamer (PDB ID 1RNA[5]). We performed two 5 μs-long standard MD simulations of each duplex and observed a stable behavior, which resulted in averaged RMSD values (calculated from the starting structures and skipping the terminal base pairs) of 1.2 ± 0.2 Å and 1.8 ± 0.4 Å for the decamer and tetradecamer, respectively. We also analyzed structural parameters of the simulated A-RNA helices. The Tables S4 and S5 show that the $OL3_{R2.7}$ *ff* variant provides reasonable description of A-RNA duplexes, nevertheless, there is a moderate reduction of inclination and roll compared to the experimental structures. Note that the global helical base pair step parameter inclination and local base pair step parameter roll are mathematically interconnected.[39] Interestingly, the $OL3_{R2.7}$ *ff* appears to reduce inclination and roll also compared to the standard OL3 *ff*. In particular, for the decamer, we identified decrease of averaged values by ~2.7° and ~1.1° for the inclination and roll, respectively. For the tetradecamer the decrease was ~4.1° and ~2.1° for the inclination and roll, respectively (see Table S4 in Supporting Information of Ref. [3] for results from standard OL3 simulations).

**Simulations of parallel-stranded dimer TERRA RNA GQ 3IBK. Comparison of simulations with standard OL3 *ff* and OL3 augmented by HRM variants.** We carried out a series of standard simulations of the 3IBK structure on 3x2 μs time scale. Adding of the HRM (for all its four variants we tested) to OL3 deteriorated the description of the two 3IBK GQ UUA propeller loops. Namely, addition of the HRM setups led to excessive formation of spurious –CH…O– interactions that bent the loop backbone into rather bizarre shapes and/or pushed the loop towards the GQ groove.

The shape of the two loops in the 3IBK crystal structure is similar, with a minor difference in base stacking (Figure S21). One loop contains stacked U:A:U bases and in the other loop the first U is shifted to a more perpendicular position to the remaining A:U stack. Both loops contain a few native –CH…O– contacts. While the GQ stem remained entirely stable, the starting loop conformations transformed into other geometries in all simulations. With the standard OL3 *ff*, the two U's were oriented into the bulk solvent, without a tendency to stack with the other bases. The A could be oriented into the solvent, too, or it was stacked atop the 5′-quartet, which was the most common conformation. On our simulation time scale, this state was the final geometry. The conformations sampled in these simulations were very similar to those sampled using the SPC/E water model.[40] We do not claim that the standard OL3 *ff* is flawless as description of GQ loops is in general challenging (including the lack of appropriate NMR primary data in loop regions in solution experiments and possible role of crystal packing in X-ray crystallography studies).[41, 42] It is possible that stacking of the A on the 5′-quartet may reflect some overstabilization of base stacking with the AMBER *ff* nonbonded terms.[43]

All tested HRM setups shifted the balance towards the structure with A stacked on the 5′-quartet, as this conformation occurred faster than with the OL3 *ff*. Although these structures looked at first sight similar to those sampled with the OL3 *ff*, the loop backbone was located visibly closer to the GQ groove with OL3 augmented by HRM variants. Simulations with all HRM setups sampled several closely related conformations characterized by the presence of a network of short non-native –CH…O– interactions between the loop backbone and the GQ stem backbone (Figure S22). Further, the loops were rigidified. Besides the conformation with stacked A (i.e., before its formation), other sampled loop conformations also displayed close contacts between loop and stem backbones and/or sharp backbone turns within the loops (Figure S23). We often observed conformations with –CH…O– interactions between nucleotide's ribose hydrogens and the consecutive nucleotide's O4′/phosphate oxygens. We hypothesize that formation of such contacts, many of which are 1…5 and 1…6 interactions, i.e., between atoms separated by 4 and 5 covalent bonds, respectively, may misbalance the parametrization of dihedral angles.

**Simulations of parallel-stranded human telomeric DNA GQ 1KF1. Comparison of simulations with standard DNA OL21 *ff* and with OL21 augmented by HRM variants.** Short –CH…O– interactions can be found also in some DNA structures[44] and the HRM should in principle be fully transferable to DNA *ff*s. We thus simulated the 1KF1 parallel-stranded GQ, which is a DNA equivalent of the RNA TERRA GQ, again on the 3×2μs time scale. We used the latest OL21[19] DNA *ff* as the core *ff*. As for the RNA GQ, adding the HRM (for all its four variants we tested) again visibly deteriorated the simulations, and in fact the worsening was even considerably larger than for the 3IBK RNA GQ. The DNA TTA propeller loops are more flexible than the RNA UUA propeller loops, which are rigidified by H-bonds involving the ribose 2′-OH group.[9] Simulations of the 1KF1 structure with OL21-HRM populated states with bent backbone stabilized by short non-native –CH…O– interactions even more than the OL3-HRM 3IBK simulations. We also observed excessive sticking of the loops into the groove. Furthermore, visible structural deteriorations now propagated also into the GQ stem.

The three TTA loops in the 1KF1 DNA GQ X-ray structure are very similar. They contain a T:A:T stack of bases, which is somewhat similar to the U:A:U stack found in one loop of the 3IBK RNA GQ. There are a few –CH…O– interactions located near the main backbone turn in the middle (Figure S24); the ones defining the loop's structural integrity are: (i) the interaction between the methyl group and/or H6 of the second T with its own phosphate, (ii) first T's sugar H2′ and/or H3′ bind to A's phosphate, and (iii) the A forms a base – phosphate interaction type 0 (0BPh)[45] intra-nucleotide interaction. All three G-quartets remained stable in the simulations. However, we observed formation of spurious –CH…O– interactions inside the GQ stem in simulations with OL21-HRM *ff*s (with all variants). Specifically, the phosphate group in the backbone could flip and form a H-bond with preceding sugar hydrogen and sometimes even with its own sugar hydrogen (Figure S25). To our opinion such interactions should not occur in GQ stems.

The loop regions were even more problematic. The typical T:A:T stack transformed into other conformations in all the simulations. With the OL21 *ff*, we observed various types of sampled loop conformations. They included formation of a T:T stack, T:A stack, T:T:A stack, stacking of the first T or A on the 3′- or 5′-quartet, respectively, or turning of a base into the groove; the numerous available conformations illustrate the flexibility of the DNA loop. Such conformations were already observed and are well documented in previous studies using the SPC/E water model and older AMBER *ff*s.[41, 46] Augmentation of OL21 by HRM variants did not help in stabilization of the native –CH…O– interactions present in the native loop of the 1KF1 DNA GQ structure. We again observed a wide range of loop conformations with various base stacks, but the backbone was affected by excessive formation of spurious –CH…O– interactions in simulations with OL21-HRM. They resulted into formation of weird sharp turns in the backbone and/or sticking of the loop into the GQ groove (Figures S26 and S27). The typical cause of the bending was formation of –CH…O– bond between a sugar hydrogen and the preceding or succeeding phosphate or O4′ atom. The loop collapse into the groove could happen regardless of whether any base was oriented towards the groove (then the base was trapped there) or all the bases were oriented towards the solvent (examples of key sampled structural states are provided as attached PDB files).

#### SUPPORTING TABLES

**Table S1:** List of interactions (H-bonds) that are modified by gHBfix21 and $gHBfix_{opt}$ potentials.[a]

| H-bonds | | gHBfix21 correction (kcal/mol) | | $gHBfix_{opt}$ correction (kcal/mol) | |
|---|---|---|---|---|---|
| Donors | Acceptors | Support | Weakening | Support | Weakening |
| NH (base) | N (base) | 0.3 | - | 0.3 | - |
| NH (base) | O (base) | 0.8 | - | 0.8 | - |
| NH (base) | O4' (sugar) | - | 1.0 | - | 1.0 |
| NH (base) | O2' (sugar) | - | 1.0 | - | 1.0 |
| NH (base) | bO (phosphate) | - | 1.0 | - | 1.0 |
| NH (base) | nbO (phosphate) | 0.1 | - | 0.1 | - |
| 2'-OH (sugar) | N (base) | 0.8 | - | 0.8 | - |
| 2'-OH (sugar) | O (base) | 0.9 | - | 0.9 | - |
| 2'-OH (sugar) | O4' (sugar) | - | 1.0 | - | 1.0 |
| 2'-OH (sugar) | O2' (sugar) | - | - | - | 1.0 |
| 2'-OH (sugar) | bO (phosphate) | - | 1.5 | - | 1.5 |
| 2'-OH (sugar) | nbO (phosphate) | - | 1.5 | - | 1.5 |

[a] See Refs. [3, 28] for details.

**Table S2**: Overview of all standard and enhanced sampling MD simulations carried out for the UUCG TL systems.[a]

| System | *ff* | Method | Runs [b] | Reps.[c] | Length [μs] |
|---|---|---|---|---|---|
| 14-mer UUCG TL | $OL3_{CP}$–gHBfix21 [d] | sMD | 10 | 1 | 20 |
| | $OL3_{CP}$–$gHBfix_{UNCG19}$ | | 10 | 1 | 20 |
| | $OL3_{CP}$–gHBfix19 | | 10 | 1 | 20 |
| | $OL3_{CP}$–gHBfix19 [e] | | 10 | 1 | 20 |
| | PAK | | 5 | 1 | 20 |
| | $OL3_{R2.7}$ | | 10 | 1 | 20 |
| | DESRES [f] | | 10 | 1 | 20 |
| | DESAMBER | | 10 | 1 | 20 |
| | ROC | | 5 | 1 | 20 |
| | ROC [g] | | 5 | 1 | 20 |
| | Chen&Garcia | | 5 | 1 | 20 |
| | BSFF1 | | 5 | 1 | 20 |
| | CHARMM36 | | 5 | 1 | 20 |
| | $CHARMM_{Drude}$ | | 10 | 1 | 5 |
| | AMOEBA | | 10 | 1 | 5 |
| 8-mer UUCG TL | $OL3_{CP}$–gHBfix21 | ST-MetaD | 3 | 12 | 5 |
| | DESAMBER | | 3 | 12 | 5 |
| | $OL3_{R2.7}$ | | 3 | 12 | 5 |
| | $OL3_{CP}$–gHBfix21 [h] | | 1 | 12 | 5 |
| | $OL3_{CP}$–$gHBfix_{UNCG19}$ [i] | | 1 | 12 | 5 |
| | DESRES | | 1 | 12 | 5 |
| | OL3 | | 3 | 12 | 5 |

[a] See Methods in the main text for details about system and *ff* labelling.
[b] Number of independent simulations for standard (sMD) and enhanced sampling (ST-MetaD) simulations.
[c] Number of replicas in ST-MetaD simulations (it is one for all sMD simulations).
[d] Data taken from Ref. [28].
[e] Simulations run with different ion parameters (see Methods in the main text).
[f] Data taken from Ref. [47].
[g] Simulations run with a different water model (see Methods in the main text).
[h] Simulation run with the original $gHBfix_{opt}$ settings (see Methods in the main text); data taken from Ref. [28].
[i] Simulation run with the $NBfix_{0BPh}$ setting, i.e., with modified vdW parameters for –H8…O5'– and –H6…O5'– atom pairs for purines and pyrimidines, respectively; data taken from Ref. [48].

**Table S3**: Summary of MD simulations of additional RNA and DNA systems.[a]

| System | Seq./PDB | *ff* | Method | Runs[b] | Reps.[c] | Length [μs] |
|---|---|---|---|---|---|---|
| r(CAAU) | CAAU | $OL3_{R2.7}$ | REST2 | 1 | 8 | 10 |
| r(CCCC) | CCCC | $OL3_{R2.7}$ | | 1 | 8 | 10 |
| r(UUUUU) | UUUUU | $OL3_{R2.7}$ [d] | sMD | 3 | 1 | 2 |
| | | OL3 [d,e] | | 3 | 1 | 2 |
| 1QC0 duplex | 1QC0 | $OL3_{R2.7}$ | | 2 | 1 | 5 |
| 1RNA duplex | 1RNA | $OL3_{R2.7}$ | | 2 | 1 | 5 |
| Kt-7 | 1S72 | DESAMBER | | 5 | 1 | 20 |
| L1-stalk *T.t.* | 3U4M | DESAMBER | | 4 | 1 | 1 |
| L1-stalk *H.m.* | 5ML7 | DESAMBER | | 4 | 1 | 1 |
| | | OL3 [f] | | 4 | 1 | 1 |
| | | $OL3_{CP}$–gHBfix21 [g] | | 4 | 1 | 1 |
| RNA GQ | 3IBK | OL3 | | 3 | 1 | 2 |
| | | $OL3_{R2.7}$ | | 3 | 1 | 2 |
| | | $OL3_{R2.7.2}$ | | 3 | 1 | 2 |
| | | $OL3_{R2.8}$ | | 3 | 1 | 2 |
| | | $OL3_{R2.8.2}$ | | 3 | 1 | 2 |
| DNA GQ | 1KF1 | OL21 | | 3 | 1 | 2 |
| | | $OL21_{R2.7}$ | | 3 | 1 | 2 |
| | | $OL21_{R2.7.2}$ | | 3 | 1 | 2 |
| | | $OL21_{R2.8}$ | | 3 | 1 | 2 |
| | | $OL21_{R2.8.2}$ | | 3 | 1 | 2 |

[a] See Supporting Information *Starting structures and simulation setup* paragraph for details about each system.
[b] Number of independent simulations for standard (sMD) and enhanced sampling (REST2) simulations.
[c] Number of replicas in REST2 simulations (it is 1 for all sMD simulations).
[d] r(UUUUU) PN simulations were performed with standard OL3 and $OL3_{R2.7}$ *ff*s, OPC water and Joung&Cheatham ions.
[e] Data taken form Ref. [37].
[f] Simulations run with SPC/E water model.
[g] Simulations run in addition with the $NBfix_{0BPh}$ setting, i.e., with modified vdW parameters for –H8…O5'– and –H6…O5'– atom pairs for purines and pyrimidines, respectively (see Ref. [34] for details).

**Table S4:** Averaged values of global helical parameters (and their standard deviations) from two 5 μs $OL3_{R2.7}$ simulations of the $r(GCACCGUUGG)_2$ decamer excised from the 1QC0[4] structure and of the $r(UUAUAUAUAUAUAA)_2$ tetradecamer 1RNA.[5] [a]

| | Helical rise [Å] | Helical incl. [°] | Tip [°] | Helical twist [°] | Major width [Å] | Minor width [Å] | X-disp. [Å] | Y-disp [Å] |
|---|---|---|---|---|---|---|---|---|
| 1QC0 (Exp.) | 2.7 | 15.2 | 2.1 | 32.3 | 18.5 | 13.1 | -4.4 | 0.0 |
| $OL3_{R2.7}$ | 2.9 ± 0.1 | 11.5 ± 0.5 | -0.4 ± 0.1 | 32.8 ± 1.3 | 18.3 ± 0.7 | 13.4 ± 0.5 | -3.7 ± 0.1 | 0.0 ± 0.0 |
| | 2.9 ± 0.1 | 11.4 ± 0.5 | -0.4 ± 0.1 | 32.8 ± 1.2 | 18.3 ± 0.7 | 13.4 ± 0.5 | -3.7 ± 0.1 | 0.0 ± 0.0 |
| 1RNA (Exp.) | 2.6 | 18.8 | -0.4 | 32.7 | 18.3 | 13.3 | -4.1 | -0.1 |
| $OL3_{R2.7}$ | 2.7 ± 0.1 | 14.4 ± 0.6 | 0.1 ± 0.2 | 34.2 ± 1.6 | 18.3 ± 0.7 | 13.3 ± 0.5 | -3.4 ± 0.1 | 0.0 ± 0.1 |
| | 2.7 ± 0.1 | 14.5 ± 0.6 | 0.0 ± 0.1 | 33.8 ± 1.3 | 18.3 ± 0.7 | 13.3 ± 0.5 | -3.4 ± 0.1 | 0.0 ± 0.0 |

[a] Simulation were run with the $OL3_{R2.7}$ *ff*, OPC water model and 0.15 M KCl (Li&Merz parameters; see Methods in the main text). The first two base pairs from each termini were excluded from the calculations to prevent end effects. Standard deviations were calculated using block average over 1000 snapshots (saved every 10 ps).

**Table S5:** Averaged values of base pair and local base pair step helical parameters (and their standard deviations) from two 5 μs $OL3_{R2.7}$ simulations of the r(GCACCGUUGG)$_2$ decamer excised from the 1QC0[4] structure and of the r(UUAUAUAUAUAUAA)$_2$ tetradecamer 1RNA.[5] [a]

| | Shear [Å] | Stretch [Å] | Stagger [Å] | Buckle [°] | Propeller [°] | Opening [°] | Tilt [°] | Shift [Å] | Slide [°] | Rise [Å] | Roll [°] | Twist [°] | Fraying (top, bottom) [b] [%] | |
|---|---|---|---|---|---|---|---|---|---|---|---|---|---|---|
| 1QC0 (Exp.) | 0.1 | -0.3 | 0.1 | 1.2 | -12.5 | 1.4 | -1.2 | -0.1 | -1.7 | 3.2 | 8.1 | 30.8 | - | - |
| $OL3_{R2.7}$ | 0.0 ± 0.0 | 0.0 ± 0.0 | 0.0 ± 0.0 | 0.4 ± 0.2 | -13.3± 0.6 | -0.3 ± 0.1 | 0.2 ± 0.1 | 0.0 ± 0.0 | -1.5 ± 0.1 | 3.3 ± 0.1 | 6.5 ± 0.3 | 31.4 ± 1.2 | 19.5 | 8.1 |
| | 0.0 ± 0.0 | 0.0 ± 0.0 | 0.0 ± 0.0 | 0.4 ± 0.2 | -13.3 ± 0.5 | -0.3± 0.1 | 0.2 ± 0.1 | 0.0 ± 0.0 | -1.5 ± 0.1 | 3.3 ± 0.1 | 6.5 ± 0.3 | 31.4 ± 1.2 | 9.7 | 13.5 |
| 1RNA (Exp.) | -0.1 | -0.2 | 0.0 | 1.3 | -18.8 | -0.4 | 0.1 | 0.1 | -1.3 | 3.3 | 10.0 | 30.5 | - | - |
| $OL3_{R2.7}$ | -0.1 ± 0.2 | -0.1 ± 0.1 | 0.1 ± 0.0 | -0.2 ± 0.5 | -16.2 ± 0.8 | 0.0 ± 1.0 | -0.1 ± 0.2 | 0.0 ± 0.1 | -1.3 ± 0.1 | 3.2 ± 0.1 | 8.6 ± 0.3 | 32.2 ± 1.6 | 42.2 | 22.9 |
| | 0.0 ± 0.0 | 0.0 ± 0.0 | 0.1 ± 0.0 | 0.0 ± 0.2 | -16.4 ± 0.6 | 0.4 ± 0.1 | 0.0 ± 0.1 | 0.0 ± 0.0 | -1.3 ± 0.1 | 3.2 ± 0.1 | 8.6 ± 0.4 | 31.8 ± 1.2 | 28.2 | 29.1 |

[a] Simulations were run with the $OL3_{R2.7}$ *ff*, OPC water model and 0.15 M KCl (Li&Merz parameters; see Methods in the main text). The first two base pairs from each termini were excluded to prevent end effects and standard deviations were calculated using block average over 1000 snapshots (saved every 10 ps).

[b] Top and bottom base pairs are $G_1C_{20}$ and $G_{10}C_{11}$ for the 1QC0 duplex and $U_1A_{28}$ and $A_{14}U_{15}$ for the 1RNA duplex, respectively.

#### SUPPORTING FIGURES

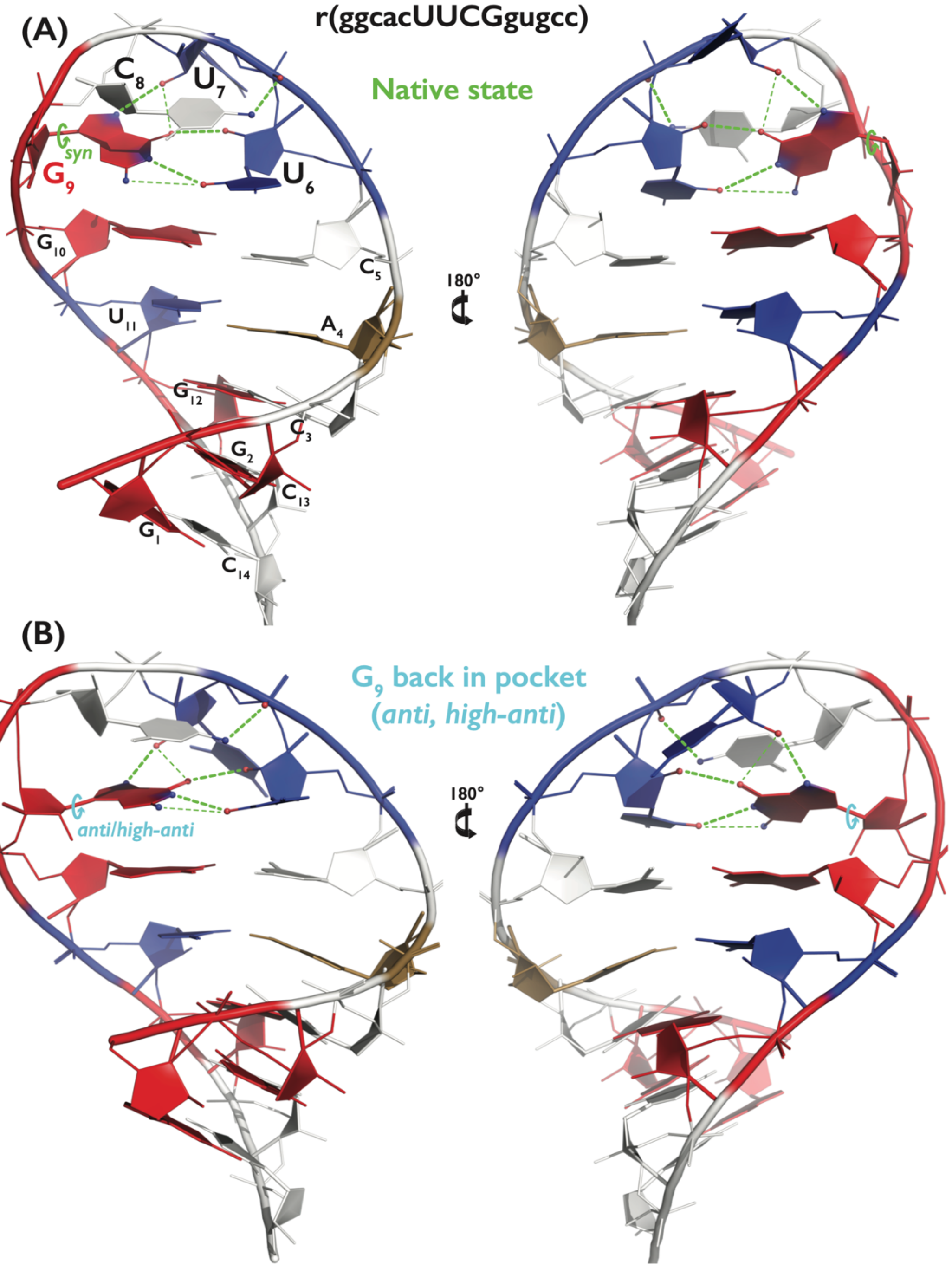

**Figure S1:** Representative snapshots of the 14-mer UUCG TL in the *native state* (A) and in the unusual 'native-like' state belonging to the broader *$G_9$ back in pocket (anti, high-anti)* state (B) sampled by the AMOEBA *ff*. A, C, G and U nucleotides are colored in sand, white, red, and blue, respectively. H-atoms, ions and water molecules are not shown for clarity. Key conformation of the $\chi_{G9}$ dihedral is colored in green (*syn*) and cyan *(anti, high-anti)* for *native* and *$G_9$ back in pocket (anti, high-anti)* states, respectively. Thick green dashed lines show native H-bonds, i.e., $G_9$(N1H)…$U_6$(O2), $U_6$(2'-OH)…$G_9$(O6), $U_7$(2'-OH)…$G_9$(N7) and $C_8$(N4H)…$U_6$(*pro*-$R_P$) interactions. Thin green dashed lines indicate two alternative H-bonds, i.e., $G_9$(N2H)…$U_6$(O2) and $U_7$(2'-OH)…$G_9$(O6), that are frequently established during MD simulations. Note that the sugar-phosphate backbone around $G_9$ and $G_{10}$ residues is significantly distorted in this AMOEBA-specific *$G_9$ back in pocket (anti, high-anti)* state.

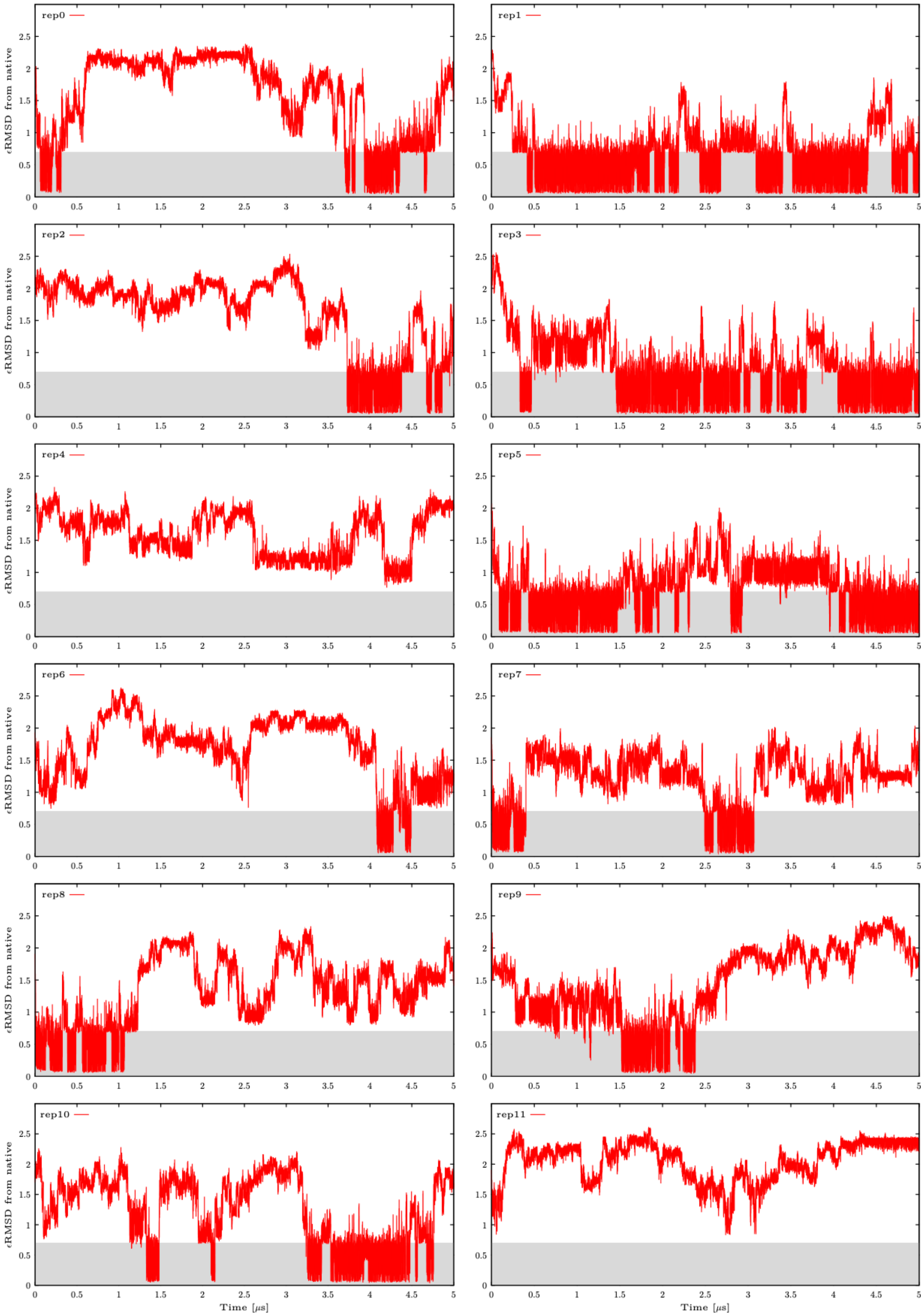

**Figure S2:** Calculated εRMSD from the native state in all twelve continuous (demultiplexed) replicas from the first ST-MetaD simulation of 8-mer UUCG TL in OL3$_{CP}$–gHBfix21 *ff*. εRMSD values were calculated every 50 ps. The shaded area highlights states close to the reference (native state, εRMSD less than 0.7).

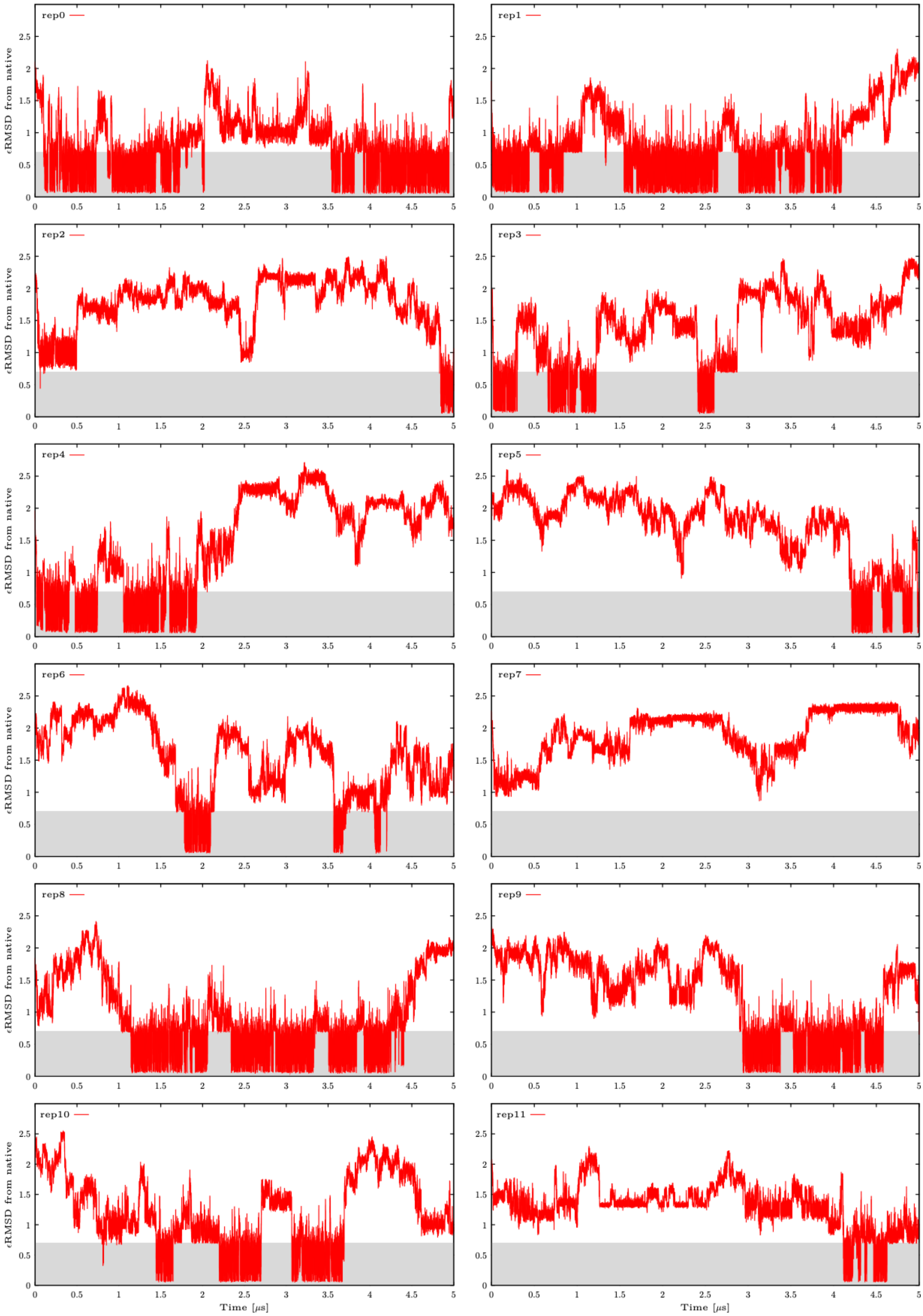

**Figure S3:** Calculated εRMSD from the native state in all twelve continuous (demultiplexed) replicas from the second ST-MetaD simulation of 8-mer UUCG TL in OL3$_{CP}$–gHBfix21 *ff*. See Figure S2 for more details.

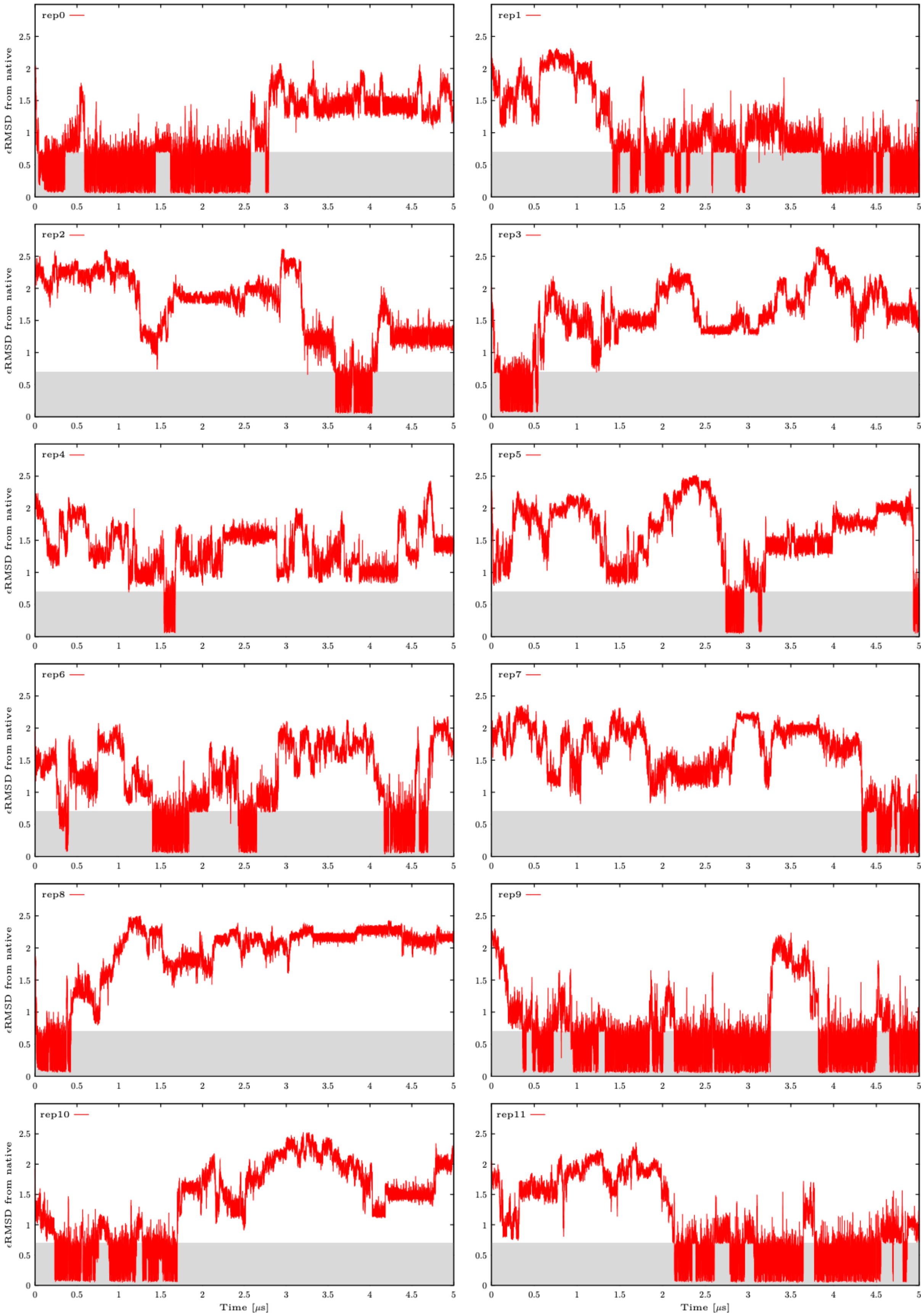

**Figure S4:** Calculated εRMSD from the native state in all twelve continuous (demultiplexed) replicas from the third ST-MetaD simulation of 8-mer UUCG TL in OL3$_{CP}$–gHBfix21 *ff*. See Figure S2 for more details.

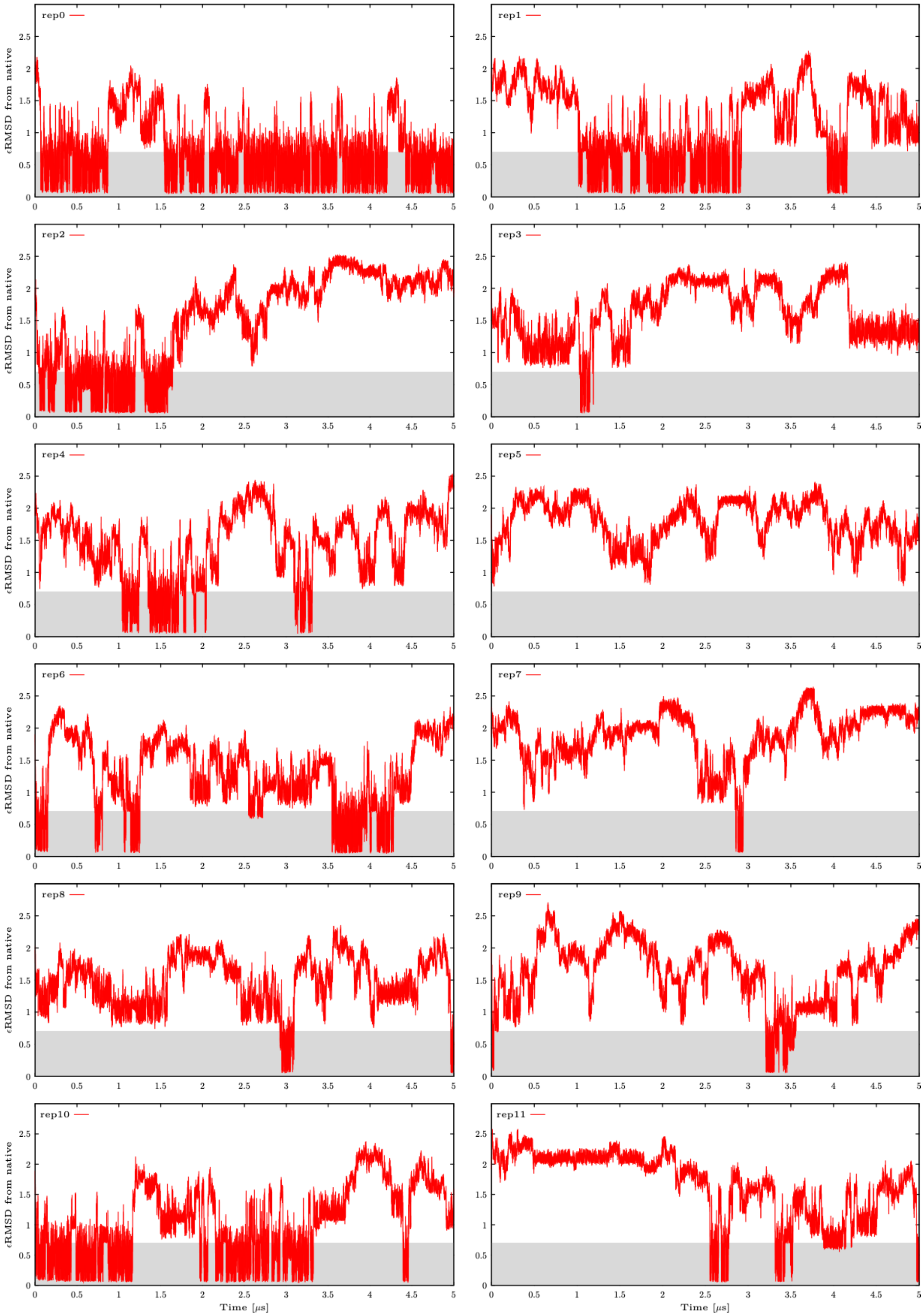

**Figure S5:** Calculated εRMSD from the native state in all twelve continuous (demultiplexed) replicas from the first ST-MetaD simulation of 8-mer UUCG TL in DESAMBER *ff*. See Figure S2 for more details.

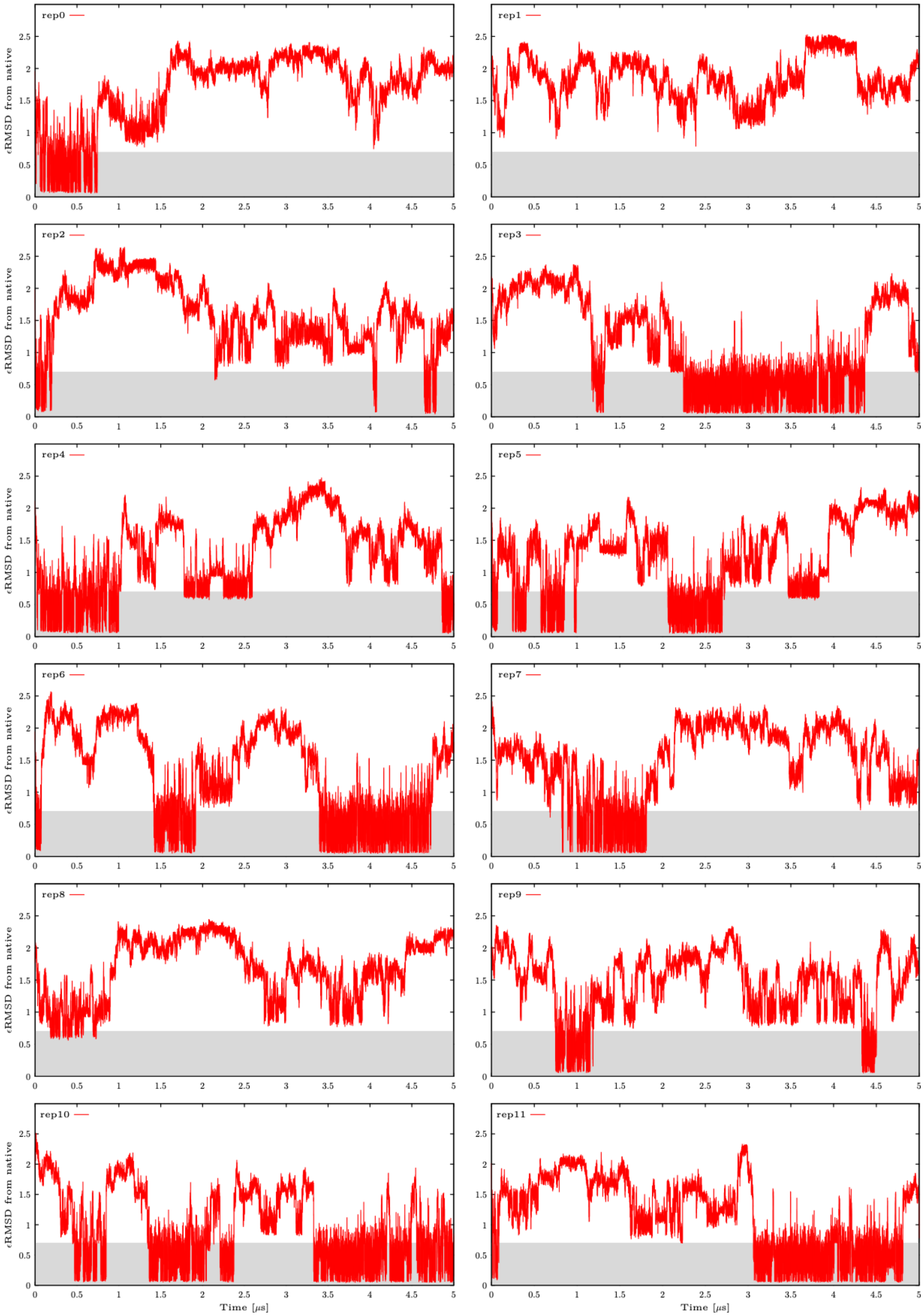

**Figure S6:** Calculated εRMSD from the native state in all twelve continuous (demultiplexed) replicas from the second ST-MetaD simulation of 8-mer UUCG TL in DESAMBER *ff*. See Figure S2 for more details.

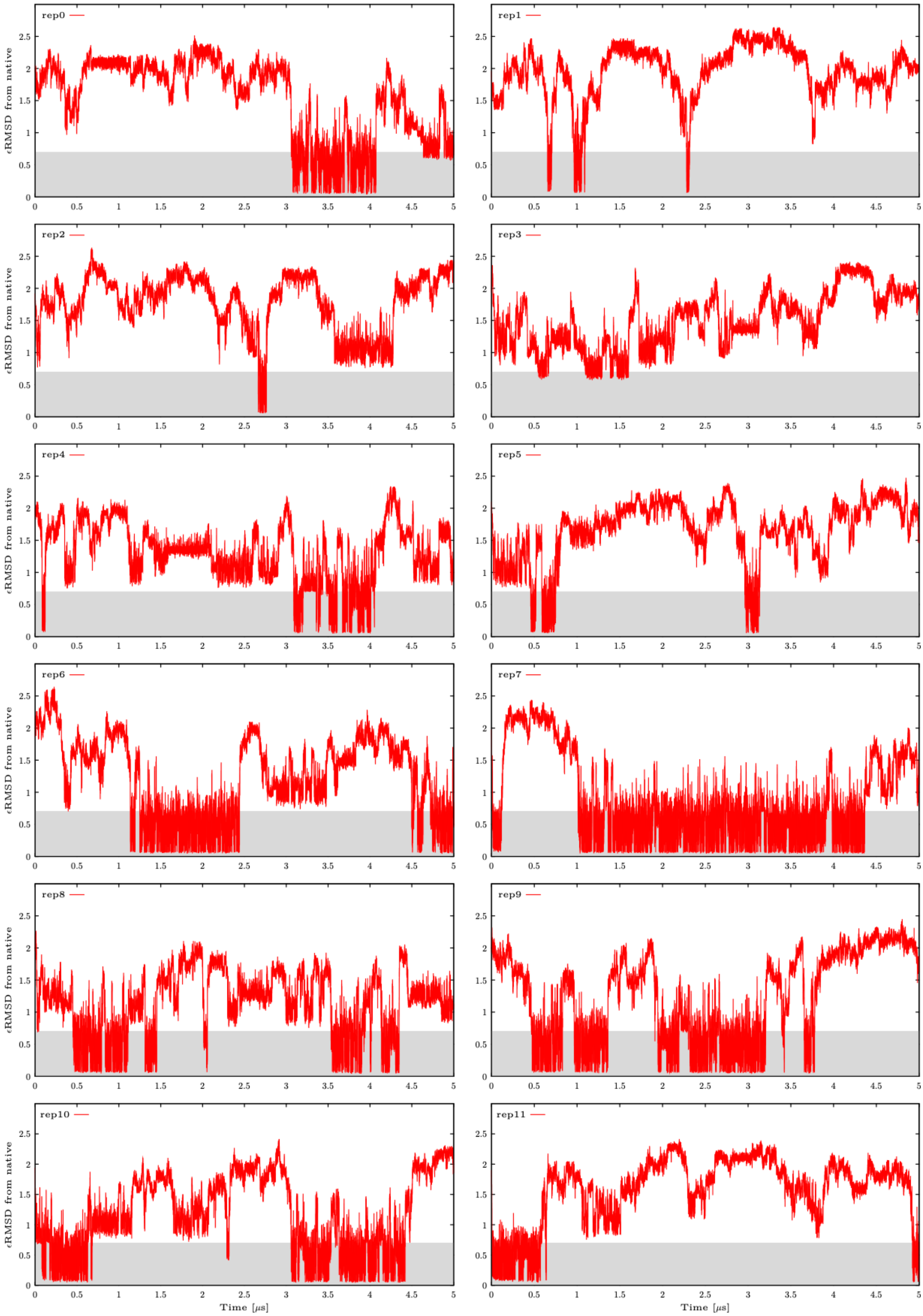

**Figure S7:** Calculated εRMSD from the native state in all twelve continuous (demultiplexed) replicas from the third ST-MetaD simulation of 8-mer UUCG TL in DESAMBER *ff*. See Figure S2 for more details.

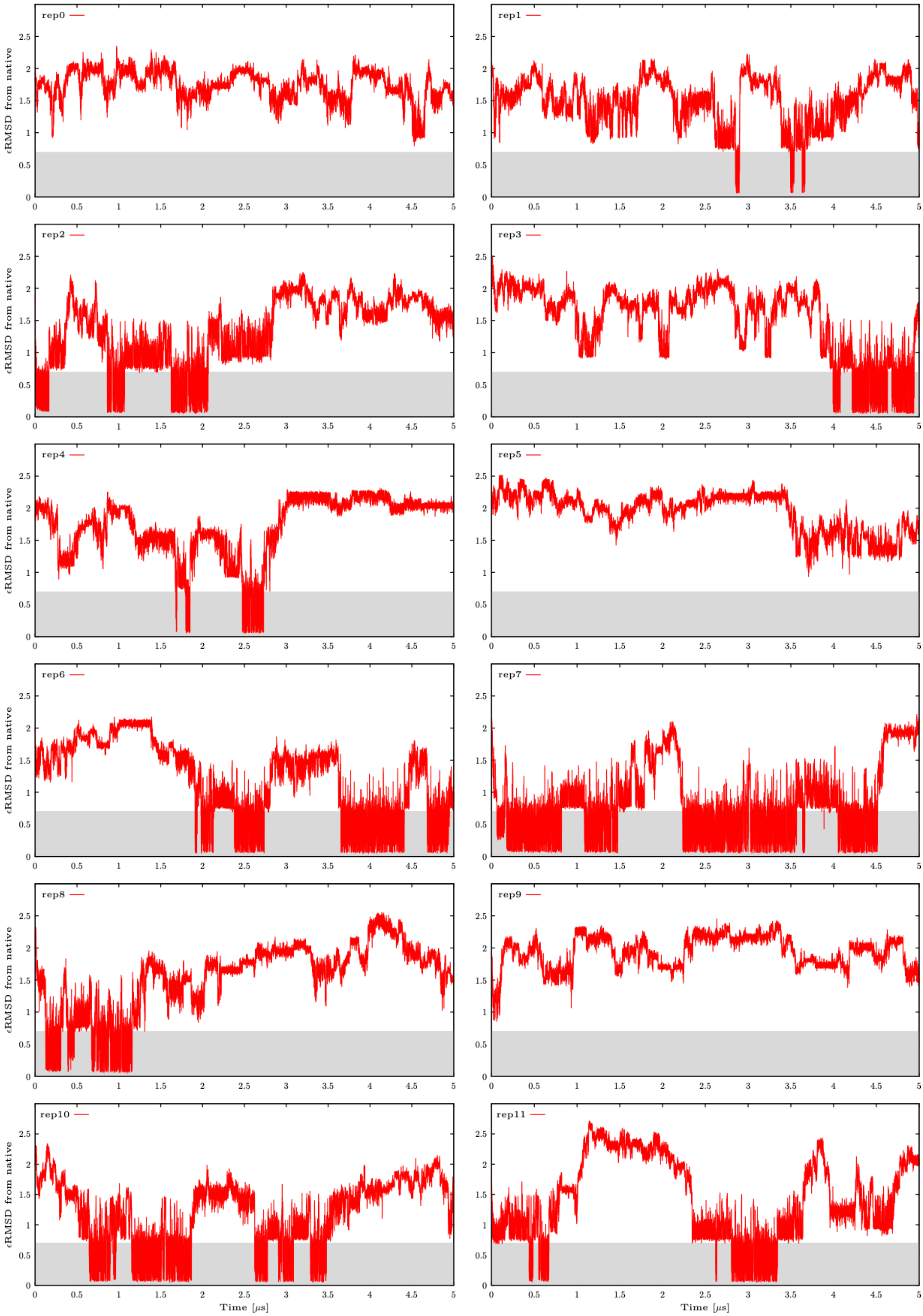

**Figure S8:** Calculated εRMSD from the native state in all twelve continuous (demultiplexed) replicas from the first ST-MetaD simulation of 8-mer UUCG TL in $OL3_{R2.7}$ *ff*. See Figure S2 for more details.

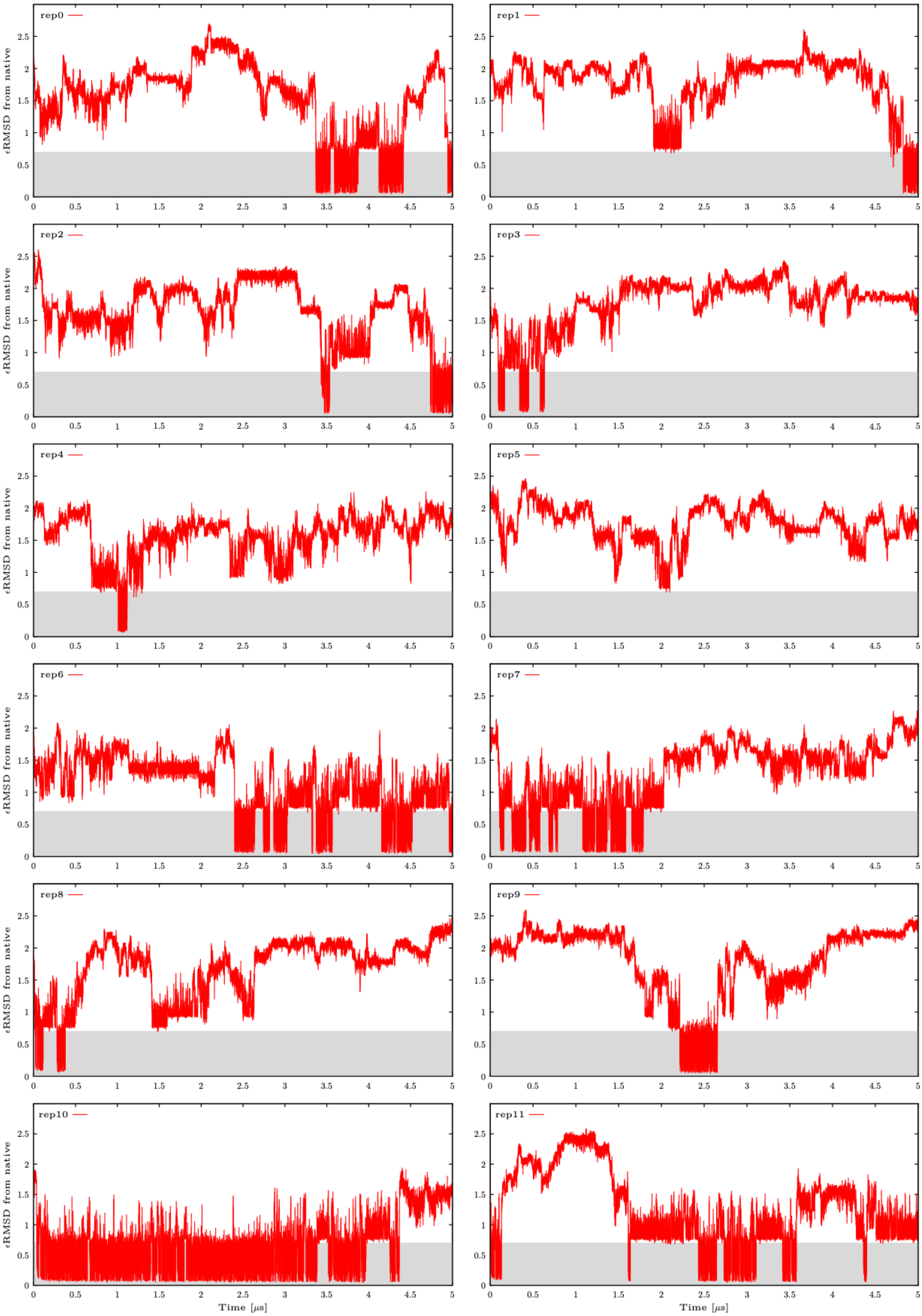

**Figure S9:** Calculated εRMSD from the native state in all twelve continuous (demultiplexed) replicas from the second ST-MetaD simulation of 8-mer UUCG TL in $OL3_{R2.7}$ *ff*. See Figure S2 for more details.

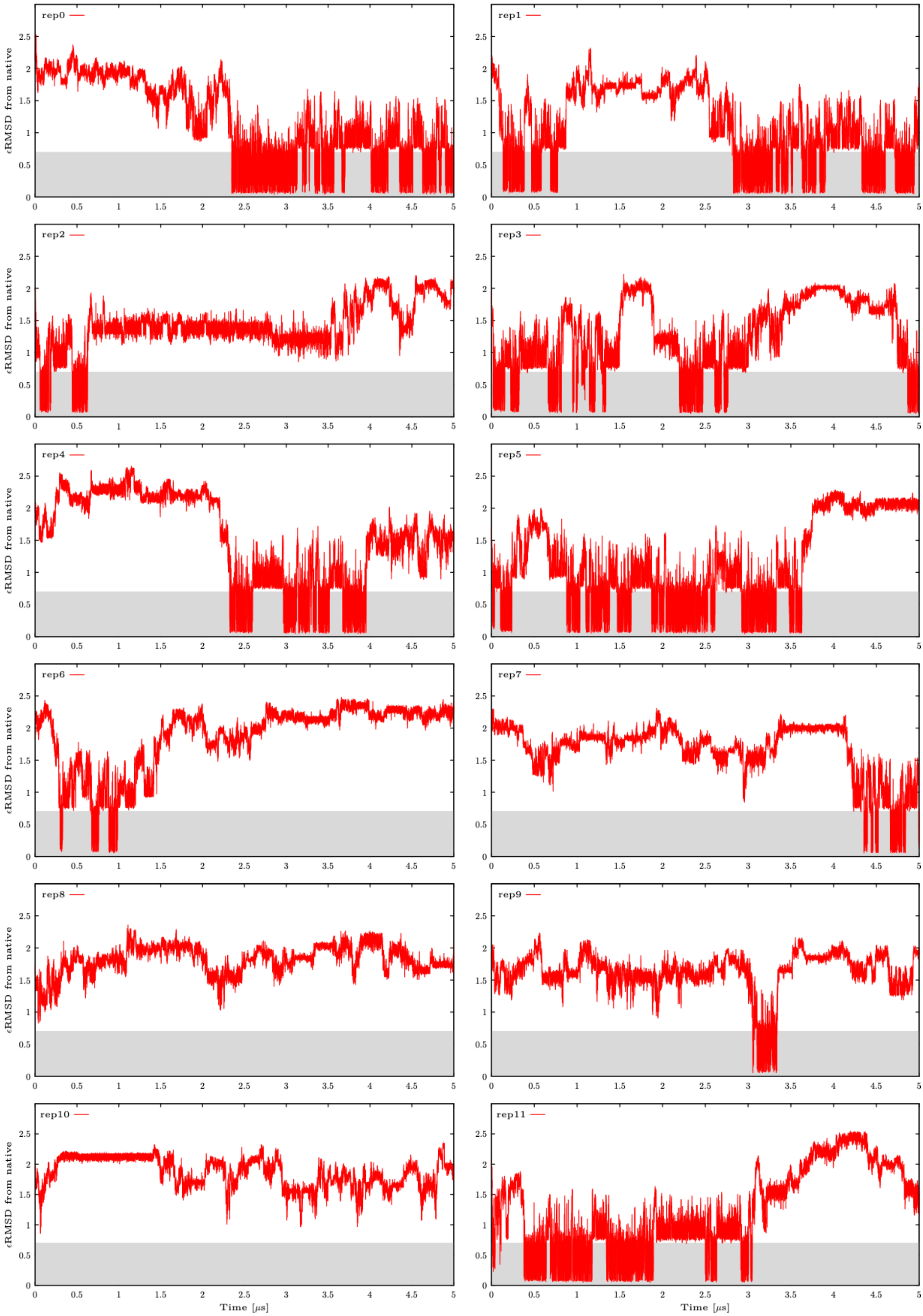

**Figure S10:** Calculated εRMSD from the native state in all twelve continuous (demultiplexed) replicas from the third ST-MetaD simulation of 8-mer UUCG TL in OL3$_{R2.7}$ *ff*. See Figure S2 for more details.

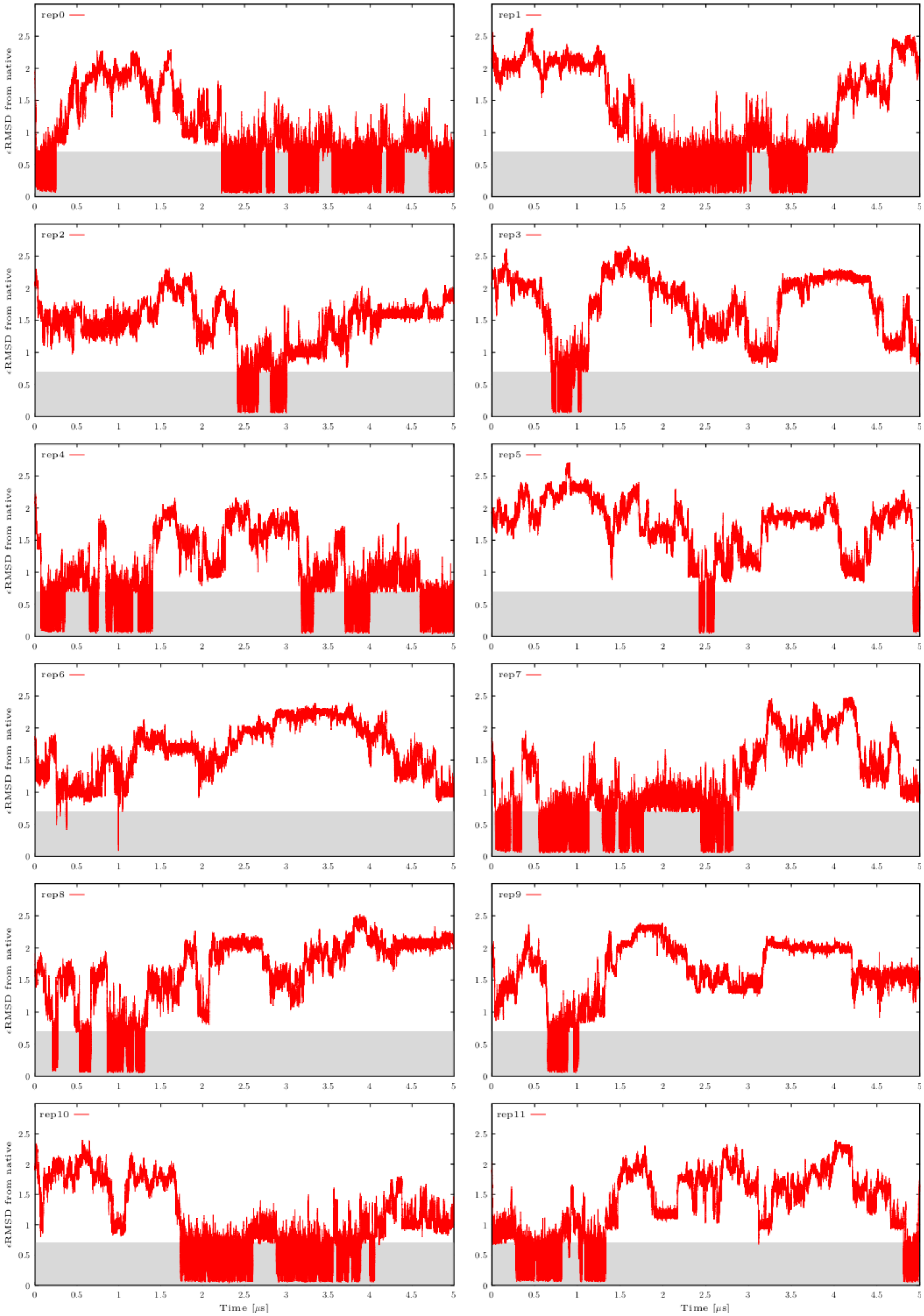

**Figure S11:** Calculated *ε*RMSD from the native state in all twelve continuous (demultiplexed) replicas from the ST-MetaD simulation of 8-mer UUCG TL in OL3$_{CP}$–gHBfix$_{UNCG19}$ *ff*. Data taken from Ref. [48]. See Figure S2 for more details.

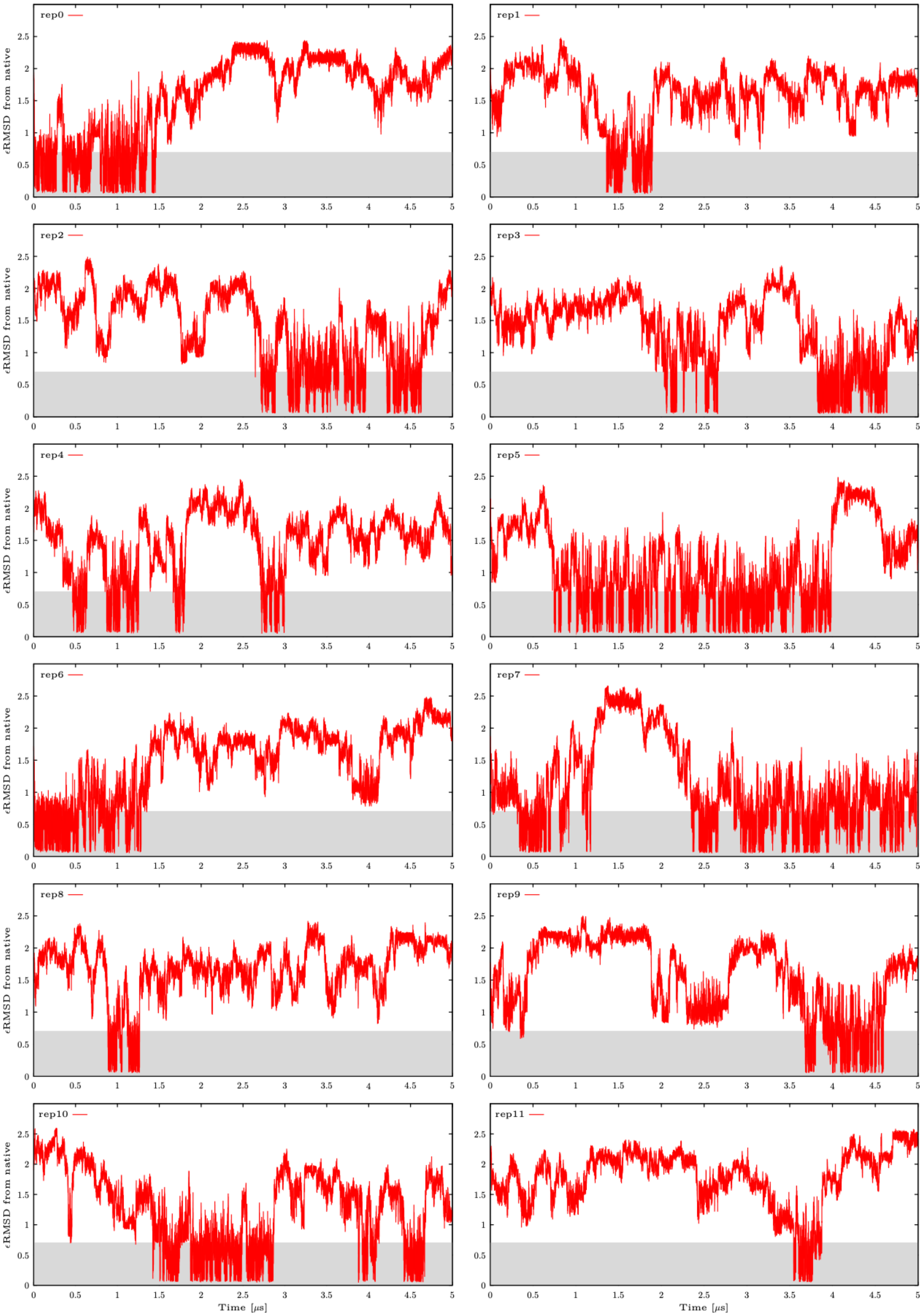

**Figure S12:** Calculated εRMSD from the native state in all twelve continuous (demultiplexed) replicas from the ST-MetaD simulation of 8-mer UUCG TL in DESRES *ff*. See Figure S2 for more details.

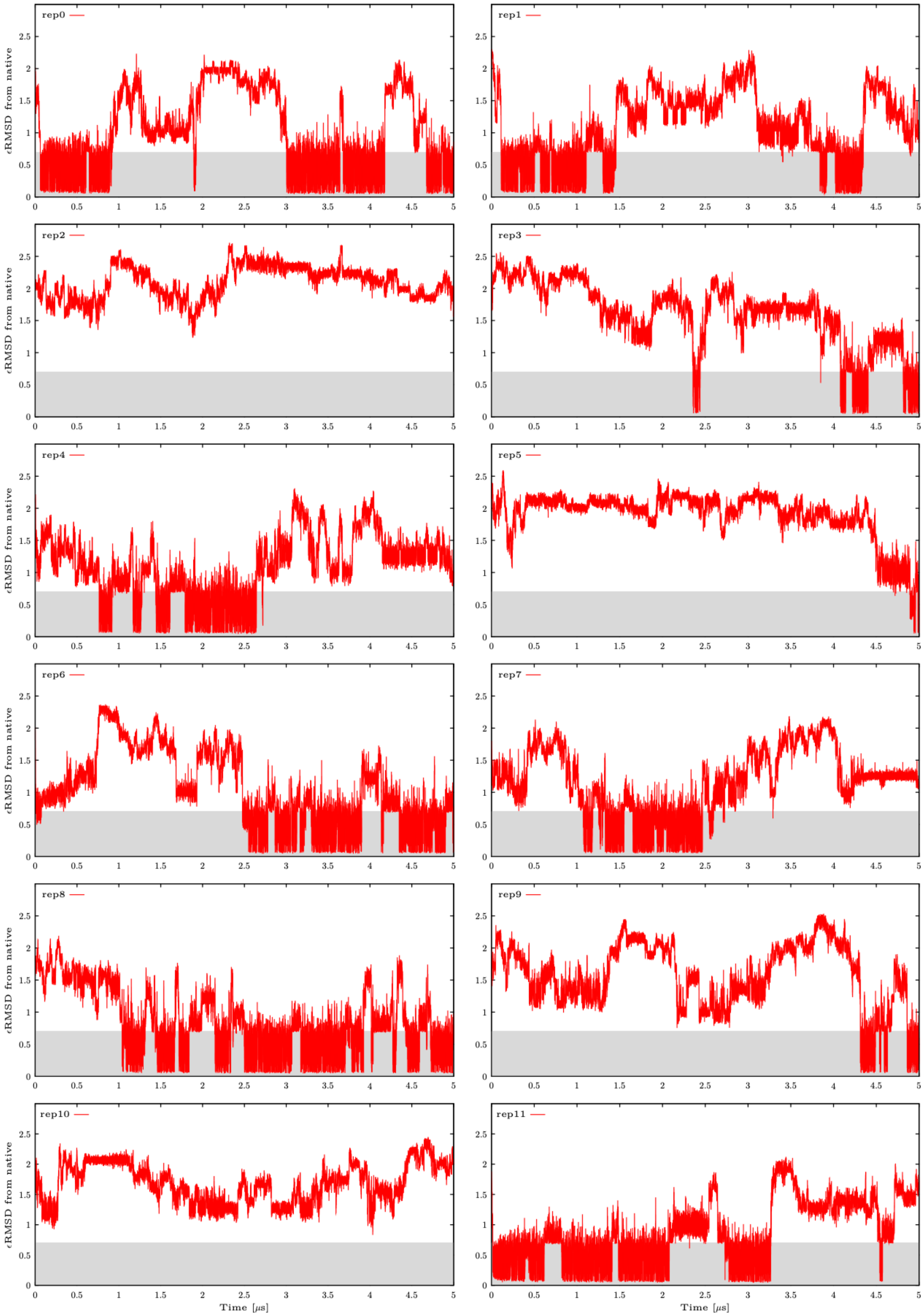

**Figure S13:** Calculated εRMSD from the native state in all twelve continuous (demultiplexed) replicas from the ST-MetaD simulation of 8-mer UUCG TL in $OL3_{CP}$–gHBfix21 *ff* (the original $gHBfix_{opt}$ settings were used; see Methods in the main text). Data taken from Ref. [28]. See Figure S2 for more details.

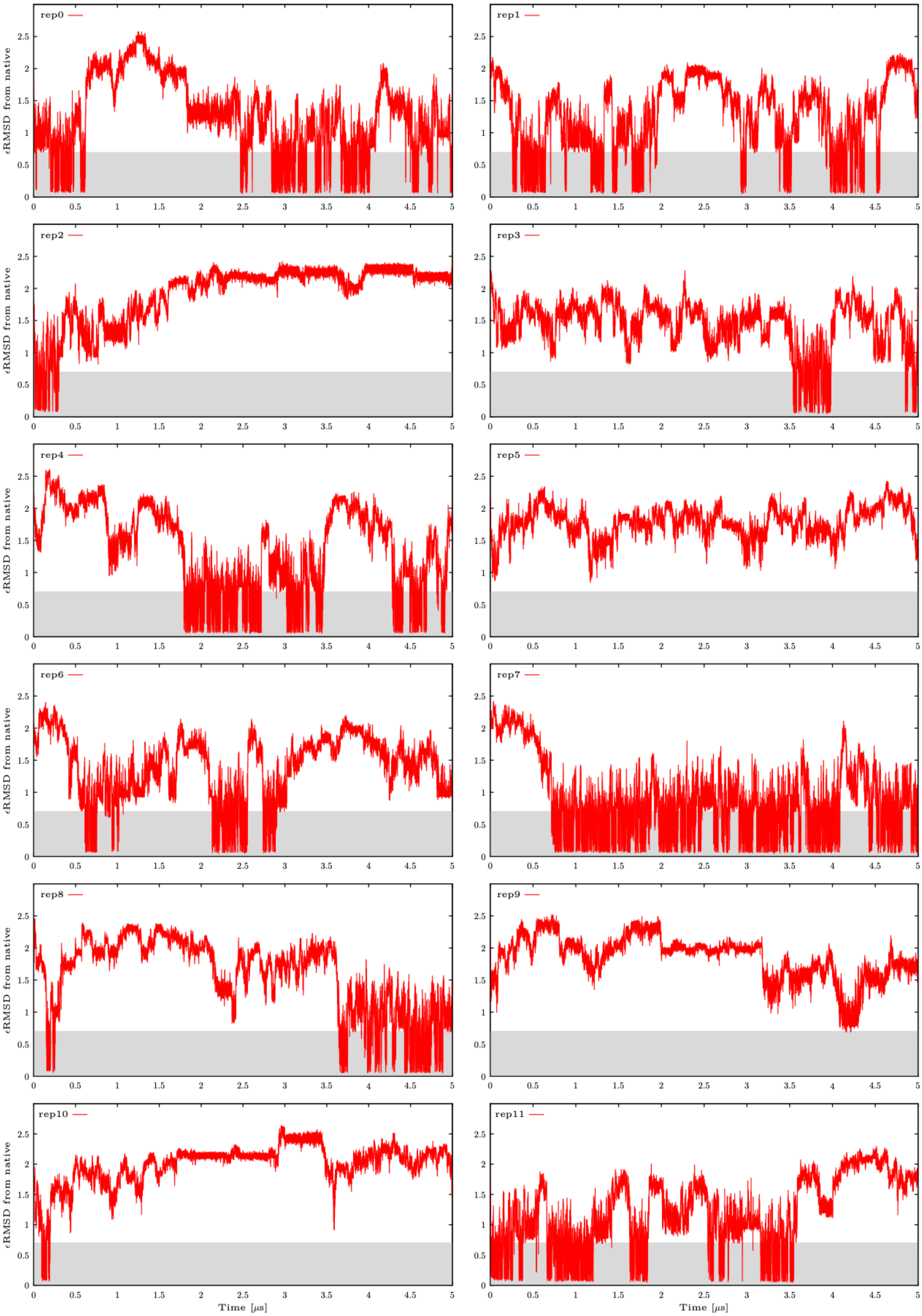

**Figure S14:** Calculated εRMSD from the native state in all twelve continuous (demultiplexed) replicas from the first ST-MetaD simulation of 8-mer UUCG TL in OL3 *ff*. See Figure S2 for more details.

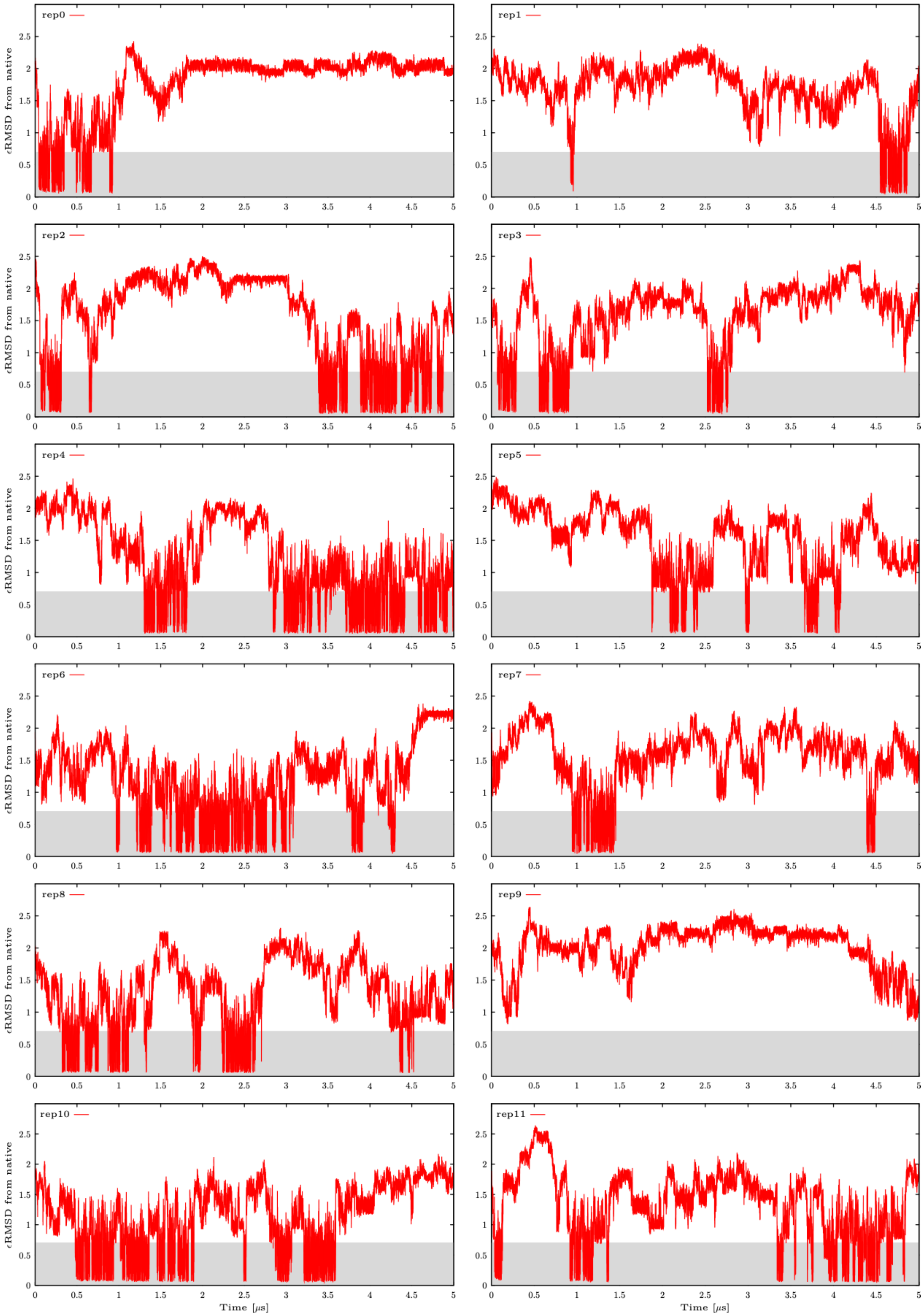

**Figure S15:** Calculated εRMSD from the native state in all twelve continuous (demultiplexed) replicas from the second ST-MetaD simulation of 8-mer UUCG TL in OL3 *ff*. See Figure S2 for more details.

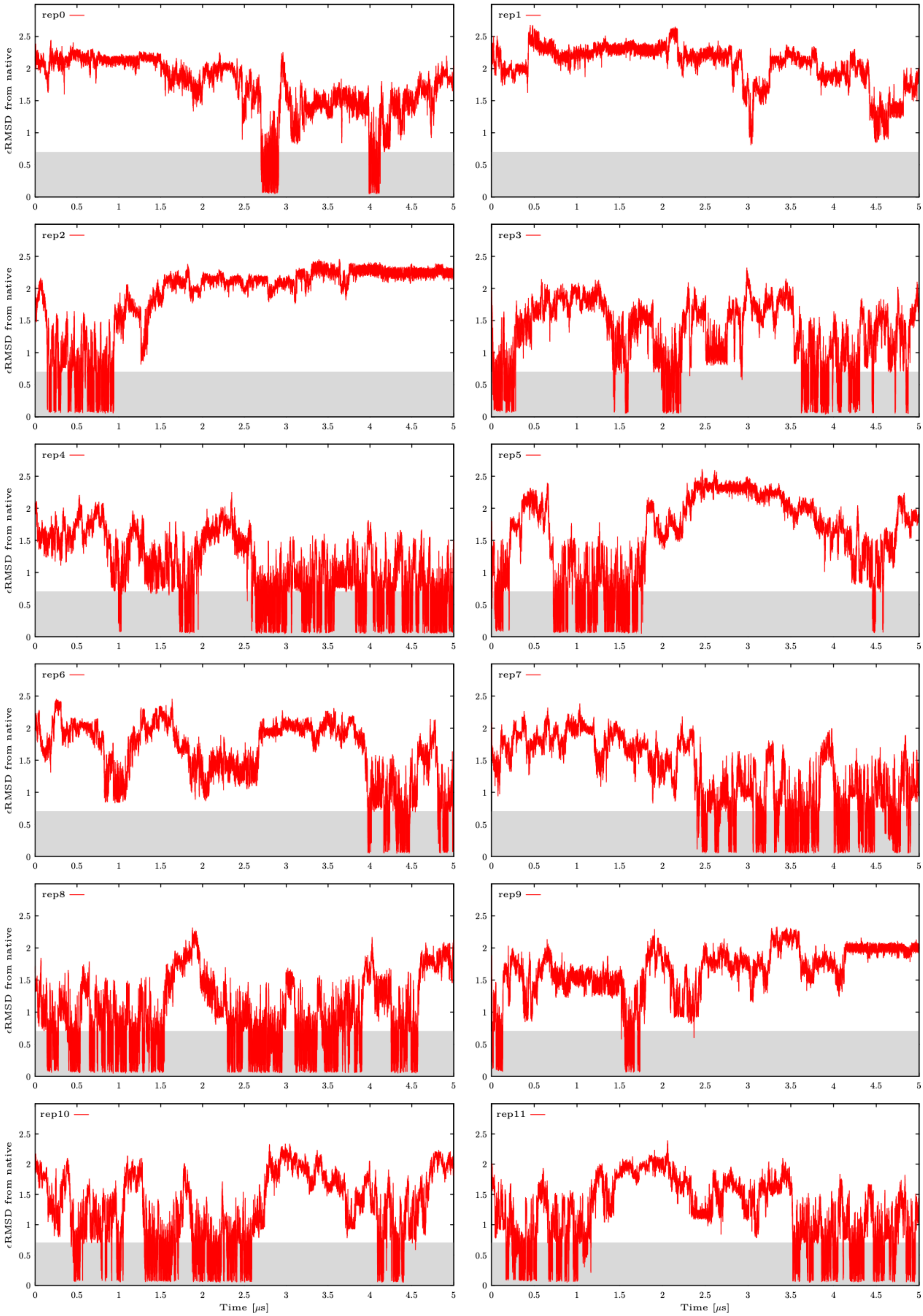

**Figure S16:** Calculated εRMSD from the native state in all twelve continuous (demultiplexed) replicas from the third ST-MetaD simulation of 8-mer UUCG TL in OL3 *ff*. See Figure S2 for more details.

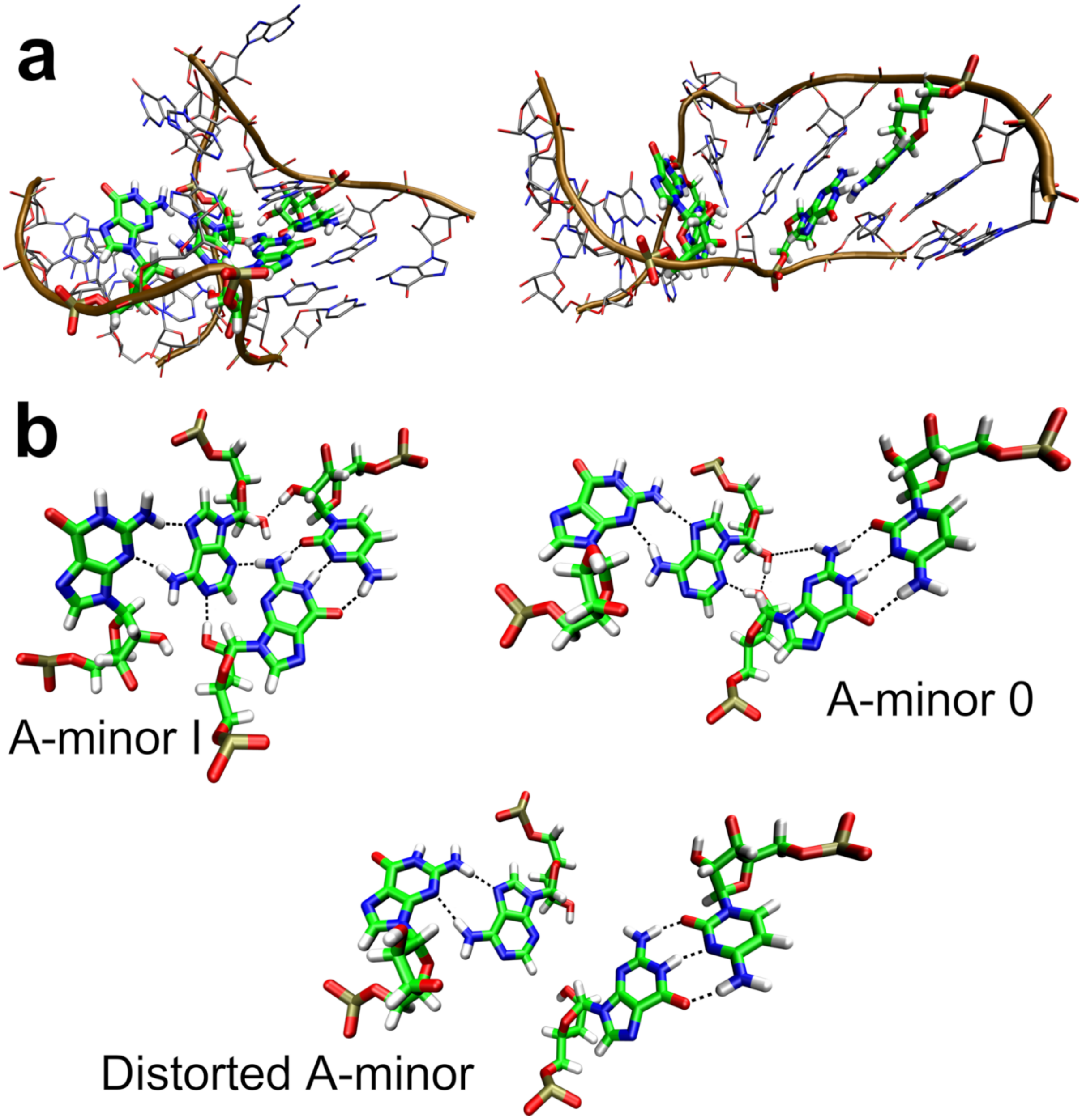

**Figure S17:** MD simulations of the Kink-turn 7 (Kt-7) from *Haloarcula marismortui* large ribosomal subunit using the DESAMBER *ff*. **a)** The native kink-turn (left) and the permanently distorted structure (right) observed in some simulations. **b)** The A-minor interaction (green carbons) variants observed in the simulations. A-minor interactions type 0 and type I are both possible for Kt-7. In addition to these, the DESAMBER *ff* exhibits a population of "distorted" or broken A-minor interaction in majority of the simulation frames. This distortion then tends to progress to global kink-turn degradation shown in panel A.

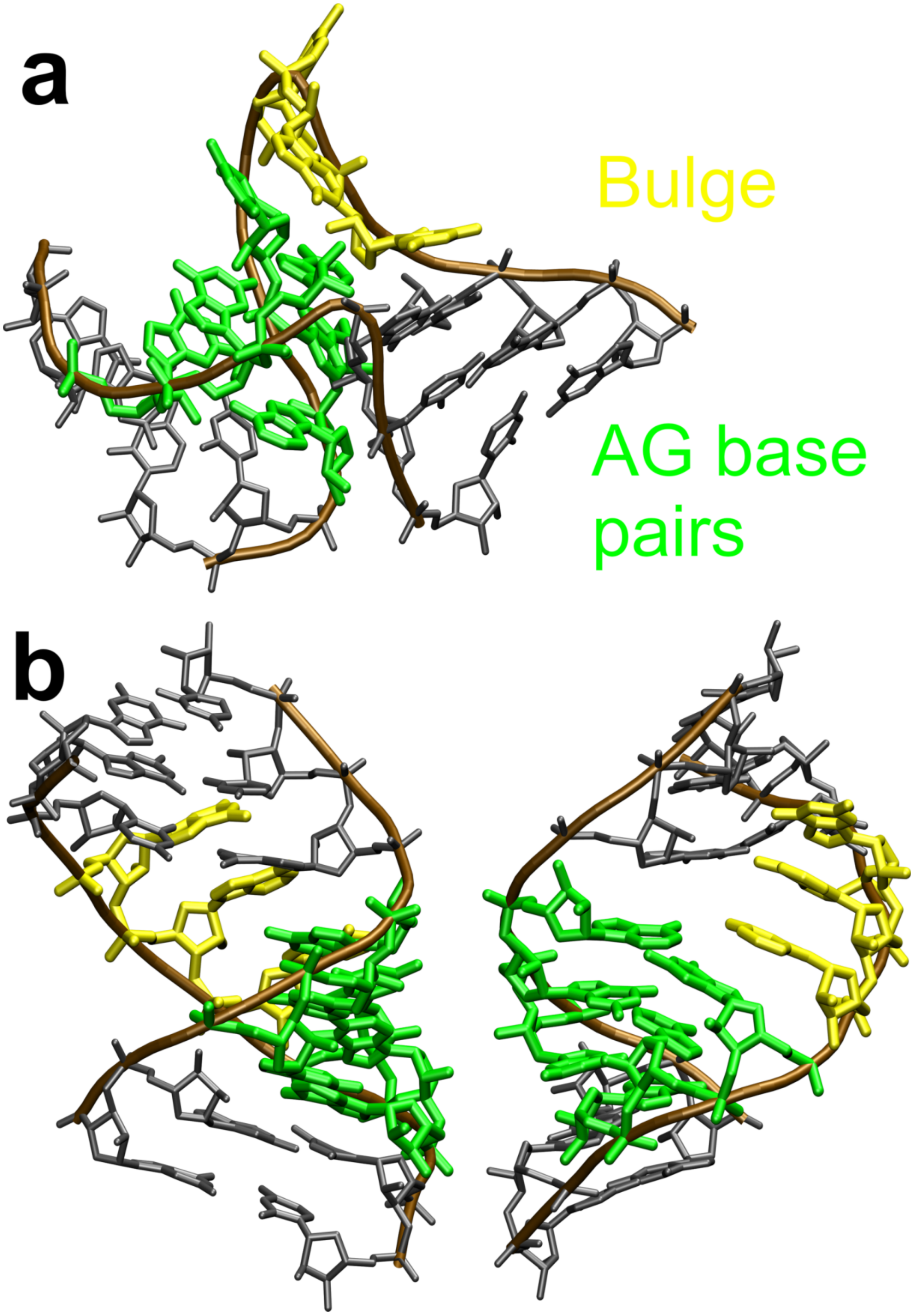

**Figure S18:** Transition of the Kt-7 into an A-form-like structure by the DESAMBER *ff*. **a)** Native kink-turn structure with the *trans* Hoogsteen/Sugar-edge AG base pairs and the bulge nucleotides marked in green and yellow, respectively. **b)** The misfolded A-RNA-like structure as observed in one of the simulations, visualized from two different angles. In two out of five 20 μs MD simulations, where we observed straightening of the kink-turn (Figure S17a, structure on the right), the RNA subsequently misfolded into a structure closely resembling a canonical A-form. In one simulation, the *trans* Hoogsteen/Sugar-edge AG base pairs even transitioned into a distorted *cis* Watson-Crick/Watson-Crick arrangement. The bulge nucleotides were pulled into the structure in both simulations, stacking on top of each other and interacting with the neighboring base pairs. The A-form-like structures, albeit fluctuating, were maintained for the rest of the simulations.

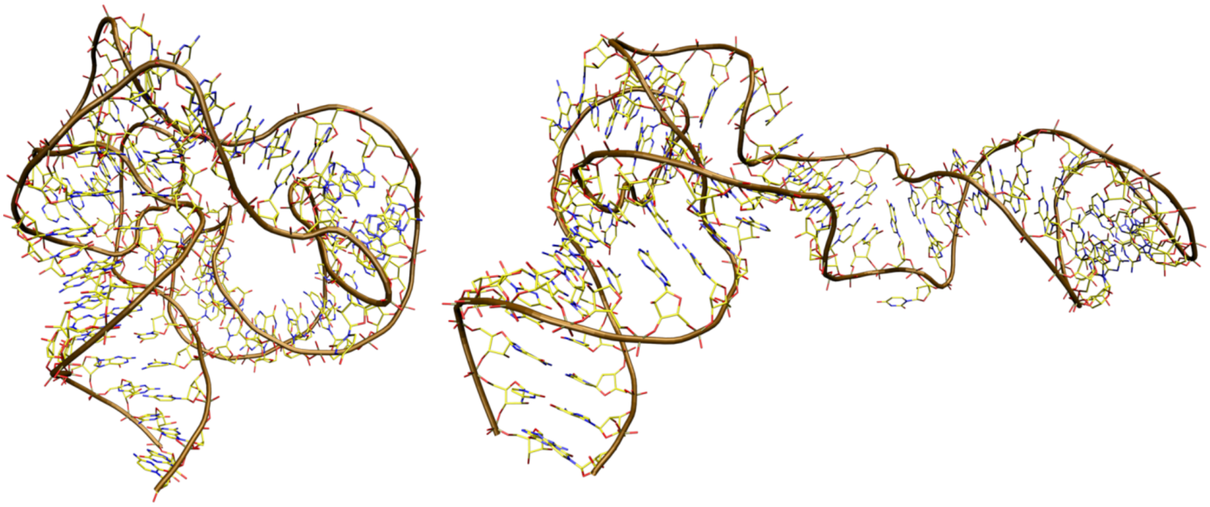

**Figure S19:** MD simulations of the L1-stalk rRNA from *Thermus thermophilus* using the DESAMBER *ff*. The native L1-stalk rRNA structure (left) and the permanently distorted structure (right) observed in all four MD simulations with the DESAMBER *ff*. A similar but even faster and more extensive structural collapse was observed in all four DESAMBER MD simulations of L1-stalk rRNA from *Haloarcula marismortui*.

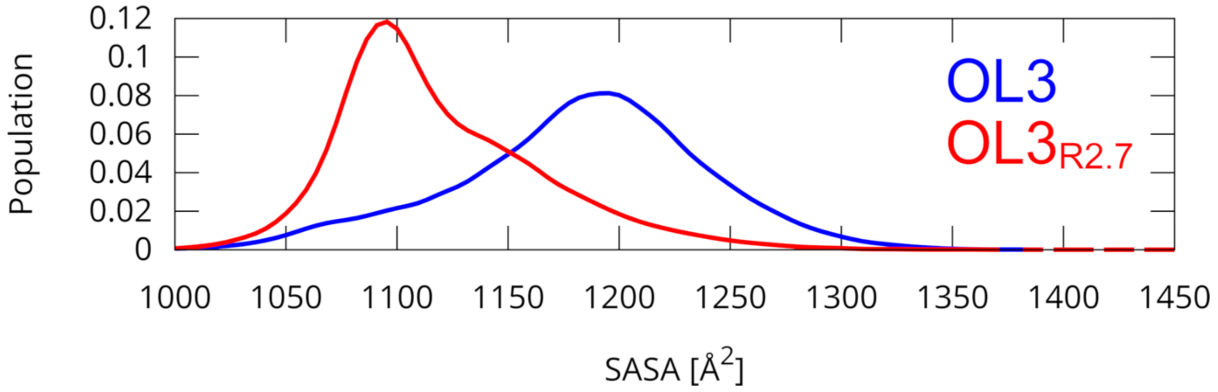

**Figure S20:** MD simulations of the r(UUUUU) PN. Distributions of the SASA (solvent-accessible surface area) measured in combined simulation ensembles of three 2 μs-long MD simulations using the standard OL3 *ff* (blue) and the $OL3_{R2.7}$ *ff* (red).

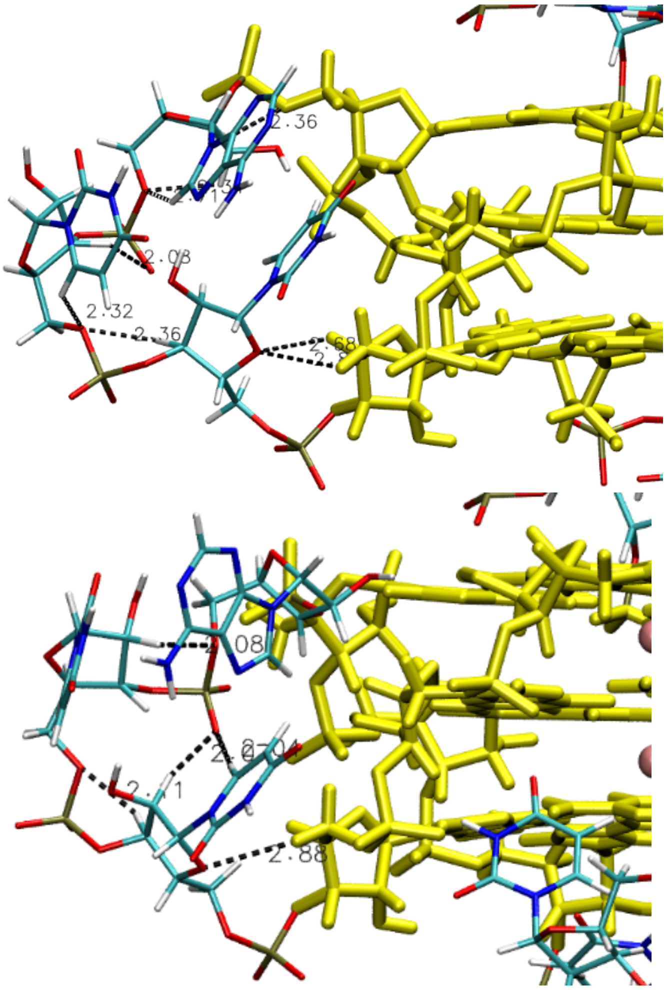

**Figure S21:** Two propeller loops in the 3IBK RNA GQ crystal structure with labeled native –CH…O– interactions. Non-G nucleotides are highlighted as lines colored specifically for C (cyan), O (red), N (blue), H (white) and P (sand) atoms, G-stem is highlighted as yellow sticks with cations in channel as pink spheres. Selected –CH…O– interactions are labeled by black dashed lines with measured distances in Å. A complete list of all –CH…O– interactions potentially affected by the HRM is given below:

The first loop (top) contains a total of **12 1…5** –CH…O– interactions potentially affected by the HRM (shorter than 3.0 Å): A8(H3')...G9(O5') 2.36 Å, A8(H3')...G9(O2P) 2.24 Å, G5(H3')...U6(O5') 2.60 Å, G5(H3')...U6(O1P) 2.04 Å, U6(H3')...U7(O5') 2.36 Å, U6(H3')...U7(O2P) 2.33 Å, U7(H3')...A8(O5') 2.37 Å, U7(H3')...A8(O2P) 2.08 Å, U6(H6)...U6(O4') 2.97 Å, U7(H6)...U7(O4') 2.19 Å, U7(H3')...U7(O5') 2.98 Å, A8(H8)...A8(O4') 2.91 Å, and a total of **13 other** –CH...O– interactions potentially affected by the HRM (shorter than 3 Å): U6(H5'')...G5(O3') 2.92 Å, G5(H5'')...U6(O4') 2.80 Å, G5(H4')...U6(O4') 2.68 Å, U6(H5')...U7(O2P) 2.99 Å, U7(H2')...A8(O5') 2.70 Å, A8(H5')...U7(O3') 2.96 Å, U7(H6)...U7(O5') 2.32 Å, A8(H2')...A8(O5') 2.34 Å, A8(H8)...A8(O5') 2.72 Å, U6(H5')...U6(O2P) 2.81 Å, U6(H5'')...U6(O2P) 2.68 Å, U7(H5'')...U7(O2P) 2.86 Å, A8(H5'')...A8(O1P) 2.42 Å.

The second loop (bottom) contains a total of **10 1…5** –CH…O– interactions potentially affected by the HRM (shorter than 3.0 Å): A20(H3')...G21(O2P) 2.71 Å, G17(H3')...U18(O5') 2.93 Å, G17(H3')...U18(O1P) 1.77 Å, U18(H3')...U19(O5') 2.11 Å, U18(H3')...U19(O2P) 2.47 Å, U19(H3')...A20(O5') 2.77 Å, U19(H3')...A20(O2P) 2.19 Å, U19(H6)...U19(O4') 2.37 Å, U19(H3')...U19(O5') 2.74 Å, A20(H3')...A20(O5') 2.85 Å, and a total of **17 other** –CH...O– interactions potentially affected by the HRM (shorter than 3.0 Å): U18(H5'')...G17(O3') 2.46 Å, A20(H5'')...G21(O2P) 2.18 Å, A20(H4')...G21(O1P) 2.25 Å, A20(H4')...G21(O2P) 2.30 Å, A20(H2')...G16(O4') 2.97 Å, G17(H5'')...U18(O4') 2.88 Å, U18(H2')...U19(O5') 2.94 Å, U18(H2')...A20(O2P) 2.08 Å, U18(H6)...A20(O2P) 2.04 Å, U19(H2')...A20(O5') 2.09 Å,

A20(H5'')...U19(O3') 2.86 Å, A20(H8)...A20(O5') 2.57 Å, U19(H5')...U19(O1P) 2.69 Å, U19(H5'')...U19(O1P) 2.94 Å, A20(H5'')...A20(O1P) 2.66 Å, A20(H3')...A20(O1P) 2.58 Å, A20(H8)...A20(O2P) 2.91 Å.

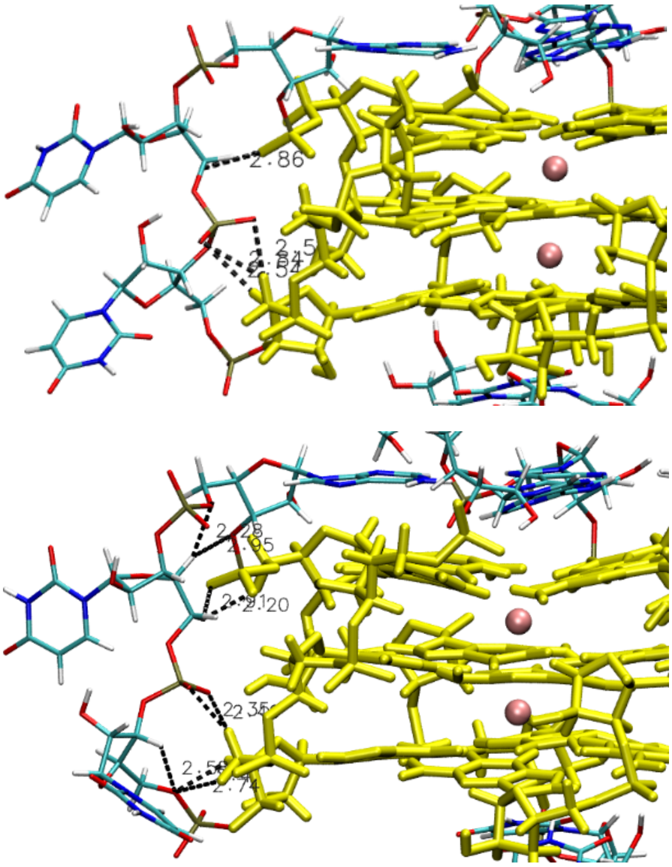

**Figure S22:** Comparison of the loop conformations (snapshots) with A stacked atop the 5'-quartet in 3IBK RNA GQ simulations. See Figure S21 for the color legend details. The standard OL3 *ff* (top panel) forms fewer –CH…O– interactions than the OL3$_{R2.7}$ *ff* (example of a sampled structure, bottom panel). As a result, the loop is placed closer to the groove with the OL3$_{R2.7}$ *ff* variant. Snapshots of both complete GQ structures are attached as the accompanying PDB files *figureS22top.pdb* and *figureS22bottom.pdb*. Only selected HRM-affected –CH…O– interactions are highlighted in panels, see the complete list below:

The top loop structure (OL3 simulation) contains a total of **9** **1…5** –CH…O– interactions potentially affected by the HRM (shorter than 3.0 Å): A8(H3')...G9(O2P) 2.81 Å, G5(H3')...U6(O1P) 2.64 Å, U6(H3')...U7(O2P) 2.82 Å, U7(H3')...A8(O1P) 2.77 Å, U6(H5'')...U6(O3') 2.75 Å, U6(H3')...U6(O5') 2.62 Å, U7(H3')...U7(O5') 2.66 Å, A8(H8)...A8(O4') 2.87 Å, A8(H3')...A8(O5') 2.60 Å, and a total of **13 other** –CH…O– interactions potentially affected by the HRM (shorter than 3.0 Å): U6(H5'')...G5(O3') 2.94 Å, U7(H5')...G9(O1P) 2.87 Å, G5(H5')...U7(O1P) 2.52 Å, G5(H5')...U7(O2P) 2.84 Å, G5(H4')...U6(O5') 2.69 Å, G5(H4')...U6(O3') 2.55 Å, G9(H5')...A8(O3') 2.92 Å, U6(H2')...U7(O2P) 2.66 Å, A8(H5'')...U7(O3') 2.81 Å, U7(H2')...U7(O5') 2.62 Å, U6(H5')...U6(O2P) 2.75 Å, U7(H5'')...U7(O1P) 2.63 Å, A8(H5')...A8(O2P) 2.64 Å.

The bottom loop structure (OL3$_{R2.7}$ simulation) contains a total of **8** HRM-affected **1…5** –CH…O– interactions (shorter than 3.0 Å): A8(H3')...G9(O2P) 2.55 Å, U6(H3')...U7(O2P) 2.48 Å, U7(H3')...A8(O1P) 2.40 Å, U6(H6)...U6(O4') 2.63 Å, U6(H3')...U6(O5') 2.58 Å, A8(H8)...A8(O4') 2.53 Å, A8(H5'')...A8(O3') 2.91 Å, A8(H3')...A8(O5') 2.48 Å, and a total of **22** HRM-affected **other** –CH…O– interactions (shorter than 3.0 Å): U6(H5)...G5(O1P) 2.99 Å, U7(H5')...G9(O1P) 2.90 Å, U7(H5')...G9(O2P) 2.21 Å, U7(H5'')...G9(O2P) 2.85 Å, A8(H2')...G9(O5') 2.93 Å,

G5(H5’)...U7(O1P) 2.45 Å, G5(H5’)...U7(O2P) 2.35 Å, G5(H5”)...U6(O5’) 2.74 Å, G5(H4’)...U6(O5’) 2.41 Å, G5(H4’)...U6(O2P) 2.71 Å, G5(H4’)...U7(O1P) 2.72 Å, U6(H2’)...U7(O2P) 2.94 Å, U7(H4’)...A8(O5’) 2.28 Å, U7(H4’)...A8(O3’) 2.95 Å, U7(H6)...U6(O3’) 2.35 Å, U6(H6)...U6(O5’) 2.31 Å, U7(H2’)...U7(O5’) 2.15 Å, U7(H6)...U7(O5’) 2.15 Å, U6(H5’)...U6(O1P) 2.50 Å, U6(H5”)...U6(O2P) 2.80 Å, U7(H5’)...U7(O1P) 2.86 Å, A8(H5’)...A8(O2P) 2.78 Å.

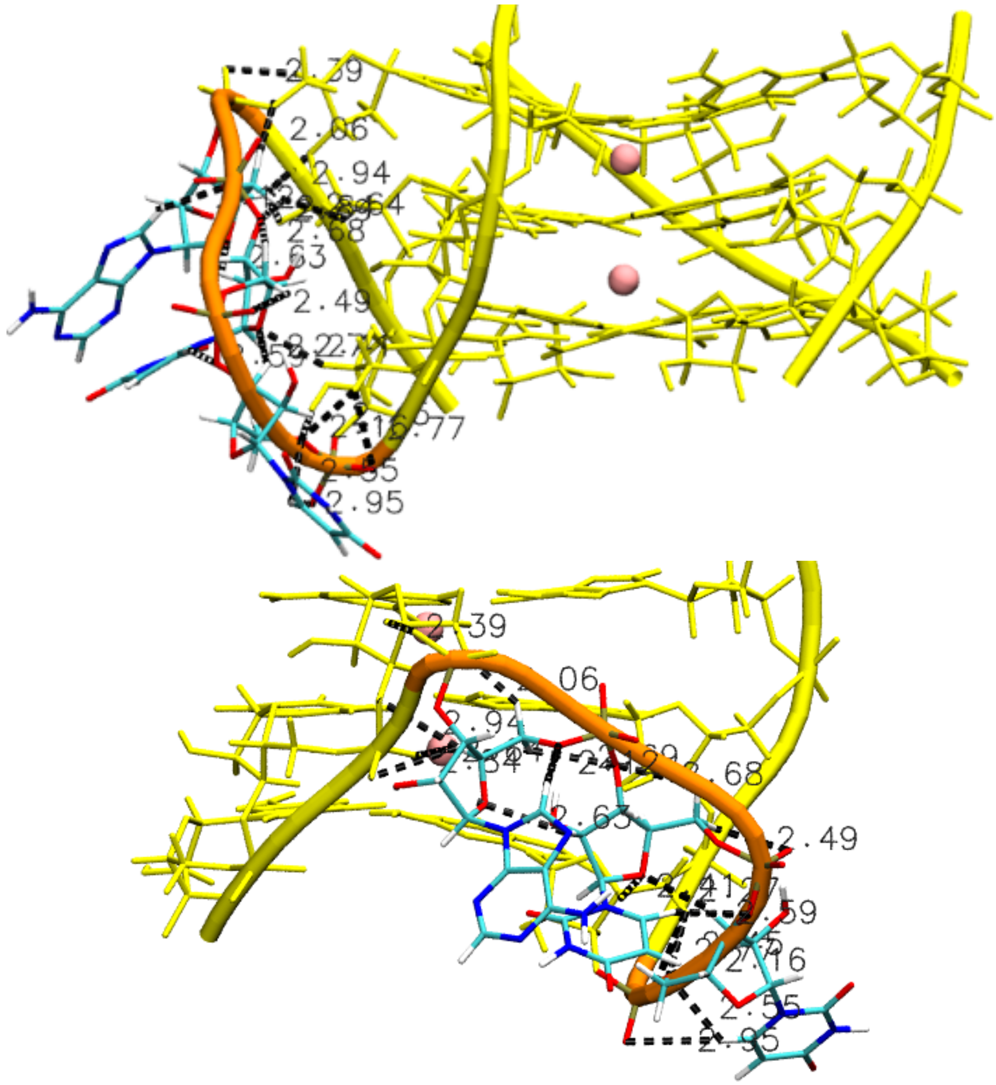

**Figure S23:** Propeller loop backbone interacting with the G-stem backbone in the simulation with $\text{OL21}_{\text{R2.7}}$ *ff* (1KF1 DNA GQ simulation snapshot), stuck in the position by several likely spurious –CH…O– interactions. Only the G-stem with the loop is shown for clarity. In the bottom figure, only the adjacent G-columns are depicted. See Figure S21 for the color legend details. In addition, the backbone is traced by the yellow and orange tube in the stem and loop, respectively. The snapshot of complete GQ structure is attached in the accompanying PDB file *figureS23.pdb*. Only selected HRM-affected –CH…O– interactions are depicted in panels, see the complete list below:

The loop contains a total of **11** HRM-affected **1…5** –CH…O– interactions (shorter than 3.0 Å): A8(H3’)...G9(O1P) 2.52 Å, G5(H3’)...U6(O1P) 2.46 Å, U6(H3’)...U7(O5’) 2.73 Å, U7(H3’)...A8(O2P) 2.58 Å, U6(H6)...U6(O4’) 2.44 Å, U6(H3’)...U6(O5’) 2.84 Å, U7(H6)...U7(O4’) 2.39 Å, U7(H5”)...U7(O3’) 2.67 Å, U7(H3’)...U7(O5’) 2.75 Å, A8(H8)...A8(O4’) 2.91 Å, A8(H3’)...A8(O5’) 2.68 Å, and a total of **25** HRM-affected **other** –CH…O– interactions (shorter than 3.0 Å): U6(H5”)...G5(O3’) 2.91 Å, A8(H5”)...G9(O5’) 2.06 Å, A8(H5”)...G9(O1P) 2.92 Å,

A8(H4')...G9(O5') 2.81 Å, A8(H4')...G9(O3') 2.95 Å, A8(H4')...G10(O1P) 2.85 Å, A8(H4')...G10(O2P) 2.64 Å, G5(H5'')...U6(O5') 2.45 Å, G5(H5'')...U6(O1P) 2.77 Å, G5(H4')...U6(O5') 2.82 Å, G5(H4')...U7(O4') 2.41 Å, U6(H5'')...U7(O4') 2.83 Å, U6(H3')...U7(O4') 2.27 Å, U7(H2')...A8(O5') 2.75 Å, U7(H2')...A8(O4') 2.64 Å, U7(H6)...U6(O3') 2.59 Å, A8(H5')...U7(O3') 2.70 Å, U6(H2')...U6(O5') 2.16 Å, U6(H6)...U6(O5') 2.56 Å, U7(H6)...U7(O5') 2.63 Å, A8(H8)...A8(O5') 2.12 Å, U6(H5')...U6(O2P) 2.40 Å, U6(H6)...U6(O2P) 2.94 Å, U7(H5')...U7(O1P) 2.48 Å, A8(H5'')...A8(O1P) 2.70 Å.

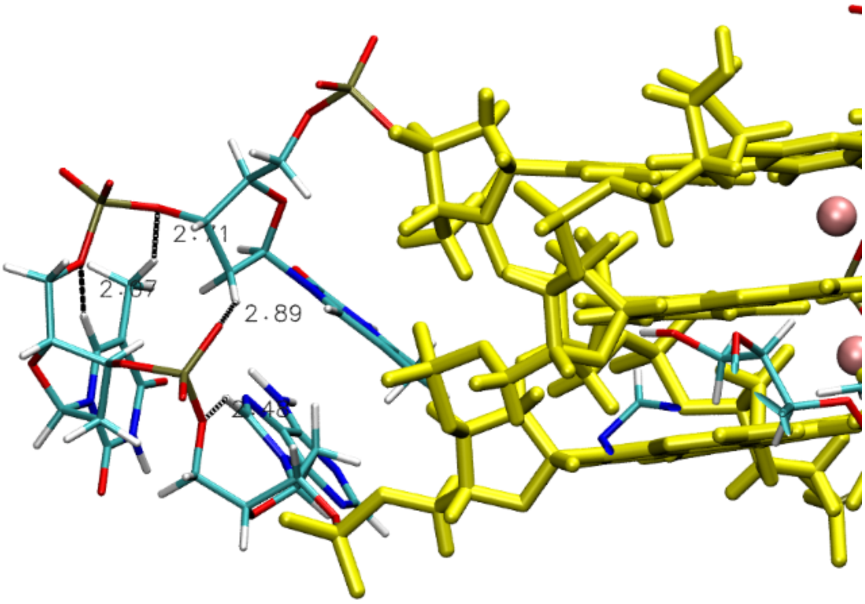

**Figure S24:** Selected native –CH…O– interactions found in the last propeller loop of 1KF1 DNA GQ crystal structure. See Figure S21 for the color legend details. A complete list of all –CH…O– interactions potentially affected by the HRM is given below:

The loop contains a total of **10 1…5** –CH…O– interactions potentially affected by the HRM (shorter than 3.0 Å): A19(H3')...G20(O2P) 2.59 Å, G16(H3')...T17(O5') 2.75 Å, G16(H3')...T17(O1P) 2.76 Å, T17(H3')...T18(O2P) 2.42 Å, T18(H3')...A19(O5') 2.87 Å, T18(H3')...A19(O2P) 2.98 Å, T18(H6)...T18(O4') 2.31 Å, T18(H5'')...T18(O3') 2.97 Å, T18(H3')...T18(O5') 2.60 Å, A19(H3')...A19(O5') 2.72 Å, and a total of **11 other** –CH…O– interactions potentially affected by the HRM (shorter than 3.0 Å): T17(H5'')...G16(O3') 2.77 Å, A19(H4')...G20(O2P) 2.96 Å, T17(H2')...A19(O2P) 2.89 Å, T18(H2')...A19(O5') 2.13 Å, T18(H71)...T17(O3') 2.72 Å, T18(H6)...T18(O5') 2.07 Å, A19(H2')...A19(O5') 2.87 Å, A19(H8)...A19(O5') 2.48 Å, T17(H5')...T17(O2P) 2.68 Å, T18(H5'')...T18(O2P) 2.36 Å, A19(H5'')...A19(O1P) 2.15 Å.

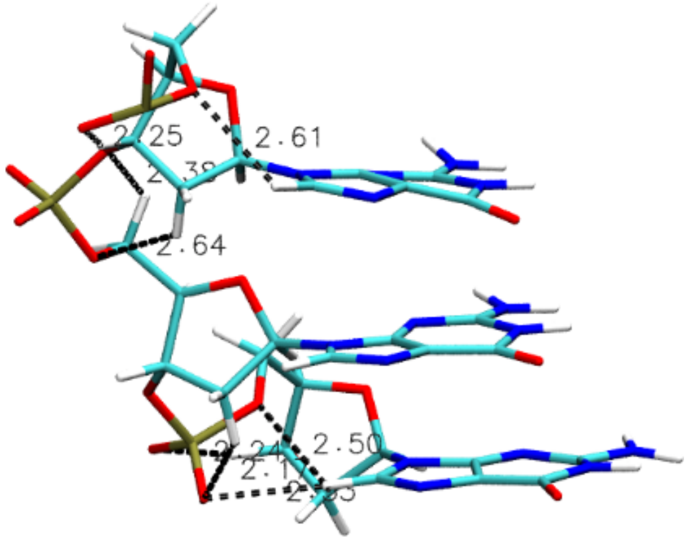

**Figure S25:** Non-native –CH…O– interactions in the G-stem of 1KF1 DNA GQ established in the simulation with $OL21_{R2.7}$ *ff* (simulation snapshot; second G-column). Only one G-column is shown for clarity. Such –CH…O– interactions were formed in all simulations with OL21 augmented by all HRM variants. The snapshot of complete GQ structure is attached in the accompanying PDB file *figureS25.pdb*. The bond lengths from top to bottom are 2.25 Å, 2.61 Å, 2.38 Å, 2.64 Å, 2.50 Å, 2.24 Å, 2.17 Å and 2.53 Å. Note that the lengths are instantaneous values for the snapshot and fluctuate during the simulation.

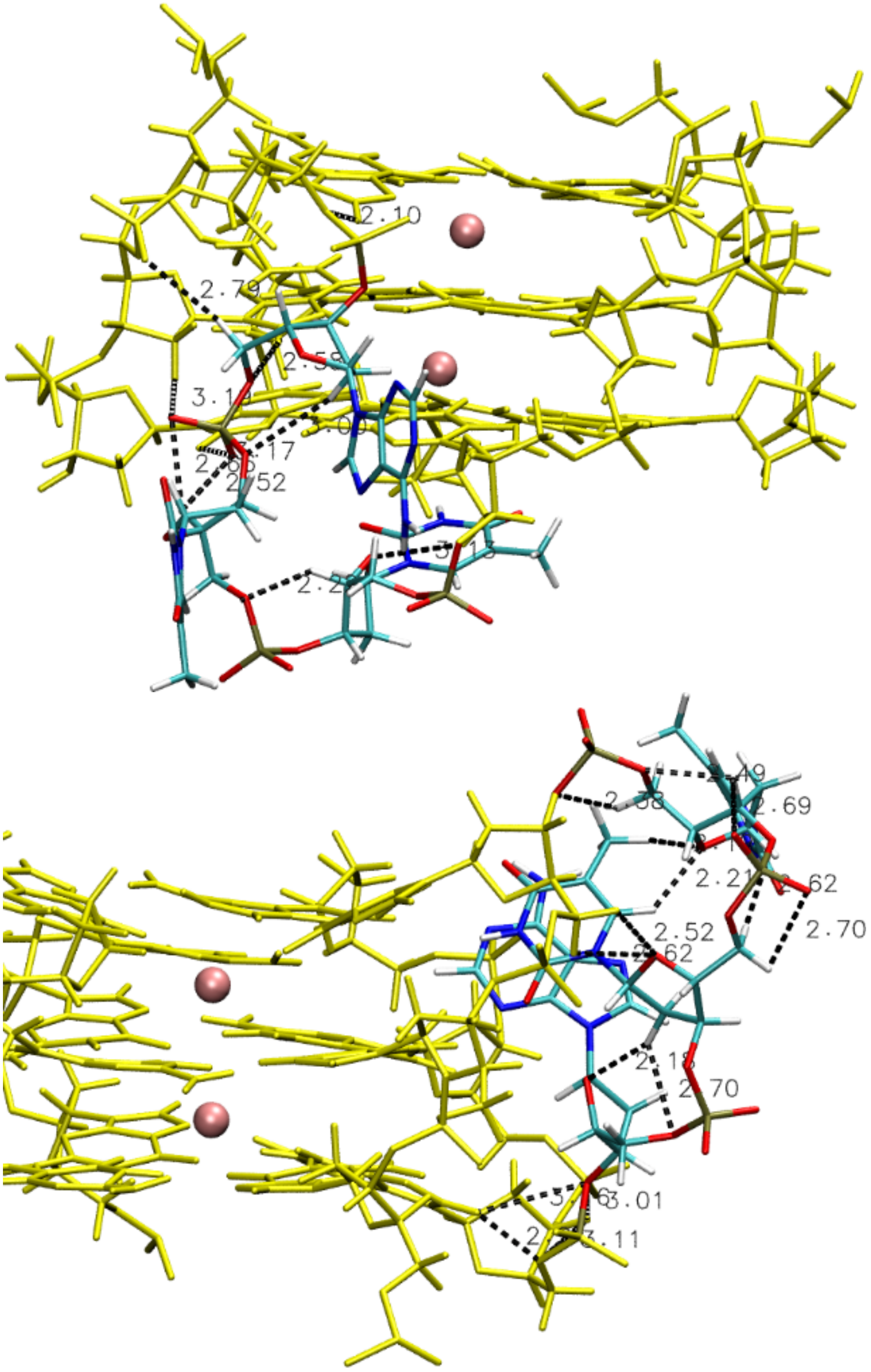

**Figure S26:** Example of two non-native loop conformations (snapshots) in 1KF1 DNA GQ supported by a network of likely spurious –CH…O– interactions in the simulation with $OL21_{R2.7}$ *ff*. Terminal (flanking) A and the other loops are not shown for clarity. Such –CH…O– interactions were found in all simulations with OL21 augmented by all HRM variants. See Figure S21 for more details. Snapshots of both complete GQ structures are attached as the accompanying PDB files *figureS26top.pdb* and *figureS26bottom.pdb*. Only selected HRM-affected –CH…O– interactions are depicted in panels, see the complete list below:

The top loop structure contains a total of **6** HRM-affected **1…5** –CH…O– interactions (shorter than 3.0 Å): T17(H3')...T18(O1P) 2.58 Å, T17(H6)...T17(O4') 2.65 Å, T17(H3')...T17(O5') 2.63 Å, T18(H6)...T18(O4') 2.55 Å, T18(H3')...T18(O5') 2.46 Å, A19(H3')...A19(O5') 2.51 Å, and a total of **20** HRM-affected **other** –CH…O– interactions (shorter than 3.0 Å): A19(H5")...G21(O2P) 2.79 Å, A19(H4')...G20(O2P) 2.14 Å, G16(H2")...T17(O5') 2.09 Å, G20(H8)...A19(O3') 2.72 Å,

G21(H2')...A19(O5') 2.83 Å, G21(H8)...A19(O5') 2.59 Å, T17(H5")...T18(O1P) 2.72 Å, T17(H4')...T18(O5') 2.22 Å, T18(H5")...T17(O3') 2.98 Å, T18(H4')...A19(O1P) 2.67 Å, T18(H4')...A19(O2P) 2.53 Å, A19(H5')...T18(O3') 2.57 Å, A19(H2')...T18(O3') 2.99 Å, A19(H8)...T18(O3') 2.68 Å, T18(H2')...T18(O5') 2.52 Å, T18(H6)...T18(O5') 2.59 Å, A19(H2')...A19(O5') 2.37 Å, T17(H5")...T17(O1P) 2.56 Å, T18(H5')...T18(O2P) 2.49 Å, A19(H5')...A19(O1P) 2.55 Å.

The bottom loop structure contains a total of **7** HRM-affected **1…5** –CH…O– interactions (shorter than 3.0 Å): A13(H3')...G14(O2P) 2.68 Å, G10(H3')...T11(O1P) 2.29 Å, T11(H3')...T12(O1P) 2.68 Å, T12(H3')...A13(O2P) 2.51 Å, T11(H3')...T11(O5') 2.50 Å, T12(H6)...T12(O4') 2.70 Å, A13(H3')...A13(O5') 2.47 Å, and a total of **17** HRM-affected **other** –CH…O– interactions (shorter than 3.0 Å): T11(H5")...G10(O3') 2.38 Å, A13(H5")...G14(O1P) 2.68 Å, A13(H4')...G14(O1P) 2.31 Å, G10(H5')...T12(O4') 2.62 Å, G10(H5")...T12(O4') 2.52 Å, G14(H2')...A13(O3') 2.33 Å, T11(H4')...T12(O5') 2.55 Å, T11(H4')...T12(O4') 2.81 Å, T12(H5")...T11(O3') 2.63 Å, T12(H2")...A13(O5') 2.70 Å, T12(H2")...A13(O4') 2.17 Å, T12(H6)...T11(O4') 2.21 Å, T12(H71)...T11(O4') 2.19 Å, A13(H5')...T12(O3') 2.84 Å, T11(H5')...T11(O2P) 2.92 Å, T11(H5")...T11(O2P) 2.99 Å, T12(H5')...T12(O2P) 2.70 Å.

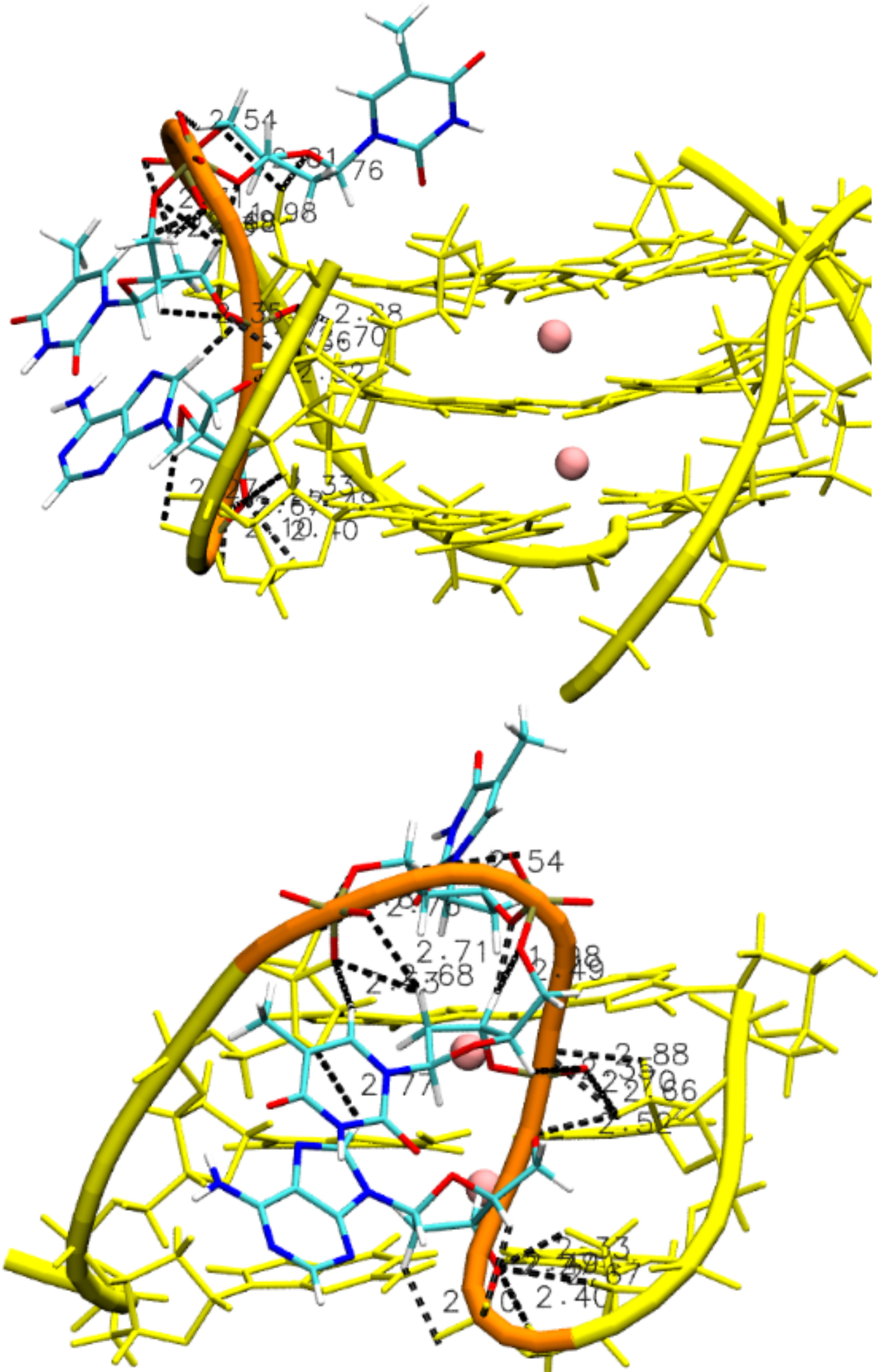

**Figure S27:** Propeller loop collapsed into the grove in the simulation with OL21$_{R2.8}$ *ff* (snapshot, two different views), stuck in the position by a series of spurious –CH…O– interactions. Only the G-stem with the loop is shown for clarity. In the bottom figure, only the adjacent G-columns are depicted. See Figure S21 for the color legend details. In addition, the backbone is traced by the yellow and orange tube in the stem and loop, respectively. Snapshot of the complete GQ structure is attached in the accompanying PDB file *figureS27.pdb*. Only selected HRM-affected –CH…O– interactions are depicted in panels, see the complete list below:

The loop contains a total of **5** HRM-affected **1…5** –CH…O– interactions (shorter than 3.0 Å): G16(H3')...T17(O2P) 2.52 Å, T17(H3')...T18(O1P) 2.69 Å, T18(H3')...T18(O5') 2.49 Å, A19(H8)...A19(O4') 2.99 Å, A19(H3')...A19(O5') 2.40 Å, and a total of **30** HRM-affected **other** –CH…O– interactions (shorter than 3.0 Å): T17(H4')...G16(O3') 2.67 Å, T18(H2')...G16(O3') 2.68 Å, T18(H2'')...G16(O3') 2.92 Å, T18(H6)...G16(O3') 2.53 Å, A19(H4')...G20(O2P) 2.27 Å, A19(H2'')...G20(O1P) 2.10 Å, A19(H8)...G16(O4') 2.76 Å, G16(H2'')...T17(O5') 2.81 Å,

G16(H2'')...T17(O4') 2.76 Å, G20(H5')...A19(O3') 2.40 Å, G20(H3')...A19(O3') 2.67 Å, G20(H2')...A19(O3') 2.33 Å, G20(H8)...A19(O3') 2.48 Å, G21(H2')...A19(O5') 2.51 Å, G21(H2')...A19(O1P) 2.66 Å, G21(H2')...A19(O2P) 2.71 Å, G21(H2'')...A19(O2P) 2.88 Å, G21(H8)...A19(O5') 2.28 Å, G21(H8)...A19(O2P) 2.65 Å, T17(H5')...T18(O1P) 2.55 Å, T17(H2'')...A19(O2P) 2.92 Å, T18(H5'')...A19(O1P) 2.71 Å, T18(H4')...A19(O1P) 2.36 Å, T18(H3')...T17(O3') 1.97 Å, T18(H2')...T17(O1P) 2.72 Å, T18(H6)...T17(O1P) 2.58 Å, A19(H5')...T18(O3') 2.69 Å, T17(H5')...T17(O1P) 2.31 Å, T18(H5'')...T18(O2P) 2.53 Å, A19(H5')...A19(O1P) 2.50 Å.
